# The anti-angiogenic compound dimethyl fumarate inhibits the serine synthesis pathway and increases glycolysis in endothelial cells

**DOI:** 10.1101/2021.12.13.472337

**Authors:** Maria Carmen Ocaña, Chendong Yang, Manuel Bernal, Beatriz Martínez-Poveda, Hieu S. Vu, Casimiro Cárdenas, Ralph J. DeBerardinis, Ana R. Quesada, Miguel Ángel Medina

## Abstract

A pathological and persistent angiogenesis is observed in several diseases like retinopathies, diabetes, psoriasis and cancer. Dimethyl fumarate, an ester from the Krebs cycle intermediate fumarate, is approved as a drug for the treatment of psoriasis and multiple sclerosis, and its anti-angiogenic activity has been reported *in vitro* and *in vivo*. However, it is not known whether dimethyl fumarate is able to modulate endothelial cell metabolism, considered an essential feature for the angiogenic switch. By means of different experimental approximations, including proteomics, isotope tracing and metabolomics experimental approaches, in this work we studied the possible role of dimethyl fumarate in endothelial cell energetic metabolism. We demonstrate for the first time that dimethyl fumarate promotes glycolysis and diminishes cell respiration, which could be a consequence of a down-regulation of serine and glycine synthesis through inhibition of PHGDH activity in endothelial cells. This new target can open a new field of study regarding the mechanism of action of dimethyl fumarate.

## Introduction

The fumaric acid ester (FAE) dimethyl fumarate (DMF) is a methyl ester of fumaric acid (FA) that has been broadly studied in several models of disease, such as inflammatory diseases, dermatological lesions and cancer. In 2013, DMF was approved by the US Food and Drug Administration (FDA) and the European Medicines Agency (EMA) for the treatment of relapsing forms of multiple sclerosis (MS), marketed under the name of Tecfidera (previously called BG-12) (Saidu et al., 2019). DMF has also been used as an anti-psoriatic drug for more than 50 years, under the brand names Fumaderm and Skilarence (Linker & Haghikia, 2016; Mrowietz et al., 2018).

Psoriasis is an inflammatory disorder that has been associated with a persistent and maintained angiogenesis (Heidenreich, Rocken, & Ghoreschi, 2009). Angiogenesis is the formation of new blood vessels from pre-existing ones. It is a natural process during wound repair, embryonic development and the reproductive cycle. However, pathological, exacerbated and deregulated angiogenesis is related to several diseases besides psoriasis, such as retinopathies, rheumatoid arthritis and cancer (Carmeliet, 2005). Some years ago, we hypothesized that DMF anti-psoriatic effect could be related somehow to modulation of angiogenesis. Interestingly, our group characterized DMF as an anti-angiogenic compound using *in vitro* and *in vivo* models (Garcia-Caballero, Mari-Beffa, Medina, & Quesada, 2011). Simultaneously, DMF was demonstrated to exert its anti-angiogenic activity through inhibition of vascular endothelial growth factor receptor 2 (VEGFR2) expression, the main receptor for VEGF-A (Meissner et al., 2011). As mentioned by Jack Arbiser, based on these data, it seems safe to say that angiogenesis inhibition plays a role in the activity of DMF and further studies on DMF mechanisms of action seem warranted (Arbiser, 2011).

Due to their potential usefulness in the treatment of several diseases, many anti-angiogenic compounds have been characterized in the last decades (Folkman, 2007; Quesada, Munoz-Chapuli, & Medina, 2006; Ronca, Benkheil, Mitola, Struyf, & Liekens, 2017). Most of these compounds target VEGF signaling pathways, and they have demonstrated to present clinical efficacy. However, in some cases an evasive resistance to VEGF pathway inhibitors is developed (Bergers & Hanahan, 2008). Therefore, therapeutical approximations in angiogenesis-dependent diseases should rely on combined targeting of different pathways (Quesada, Medina, & Alba, 2007). Not surprisingly, endothelial cell (EC) energetic metabolism was short after shown to be essential for correct function of ECs, and hence for correct angiogenesis trigger (Eelen, Treps, Li, & Carmeliet, 2020). In consequence, targeting EC metabolism was proposed as a novel strategy for the treatment of angiogenesis-dependent pathologies (Goveia, Stapor, & Carmeliet, 2014; Ocana, Martinez-Poveda, Quesada, & Medina, 2019b).

DMF is a cell permeable FAE that can be converted into fumarate inside the cell, thus feeding the tricarboxylic acid (TCA) cycle. Diverse, cell- and dose-dependent effects of DMF on global cell metabolism have been found in different cell types. For instance, DMF exerted a differential effect on the energetics metabolism of mouse embryonic fibroblasts depending on Nrf-2 expression and time incubation (Ahuja et al., 2016). Other authors found lower respiration rates in human retinal epithelial cells treated with DMF (Foresti et al., 2015). Additionally, DMF was shown to inhibit glyceraldehyde 3-phosphate dehydrogenase (GAPDH), a glycolytic enzyme, thus down-regulating aerobic glycolysis in murine activated myeloid and lymphoid cells (Kornberg et al., 2018). Moreover, DMF was found to induce cell metabolism dysfunction in human pancreatic cells through inhibition of mitochondrial respiration, aerobic glycolysis and folate metabolism, possibly by targeting the enzyme methylenetetrahydrofolate dehydrogenase 1 (MTHFD1) (Chen et al., 2021). Nevertheless, as far as we are concerned, no studies have been performed regarding the possible role of DMF in EC energetic metabolism.

In this work, we wanted to explore the potential capacity of DMF to modulate microvascular EC glucose and/or glutamine metabolism in an *in vitro* model of microvascular ECs. We found that DMF diminishes cell respiration while it upregulates glycolysis in human dermal microvascular ECs (HMECs). Interestingly, our results show that DMF downregulates the serine and glycine synthesis pathway in these cells through inhibition of phosphoglycerate dehydrogenase (PHGDH) activity. To our knowledge, the results presented herein are the first experimental evidence showing that DMF downregulates the serine and glycine synthesis pathway in ECs. The observed alteration of EC metabolism exerted by DMF could open new horizons for further characterization of its mechanism of action in angiogenesis-dependent diseases.

## Results

### DMF inhibits capillary tube formation in microvascular endothelial cells

The anti-angiogenic activity of DMF has been previously described in an *in* vitro model of macrovascular ECs, but its effect on microvascular ECs has not been assessed before (Garcia-Caballero et al., 2011). In order to determine the concentrations of DMF that interfere with angiogenesis *in vitro* in HMECs (microvascular ECs), we evaluated the dose of DMF able to totally inhibit tube formation in these cells. For this aim, we performed a capillary tube formation assay on Matrigel using increasing DMF concentrations. Consistent with the previously published effect on macrovascular ECs, we confirmed the total inhibition of tubular-like morphogenesis on Matrigel by 50 and 100 µM DMF in microvascular ECs (Figure 1). Then we decided to use 50 and 100 µM for further experiments.

**Figure 1.**
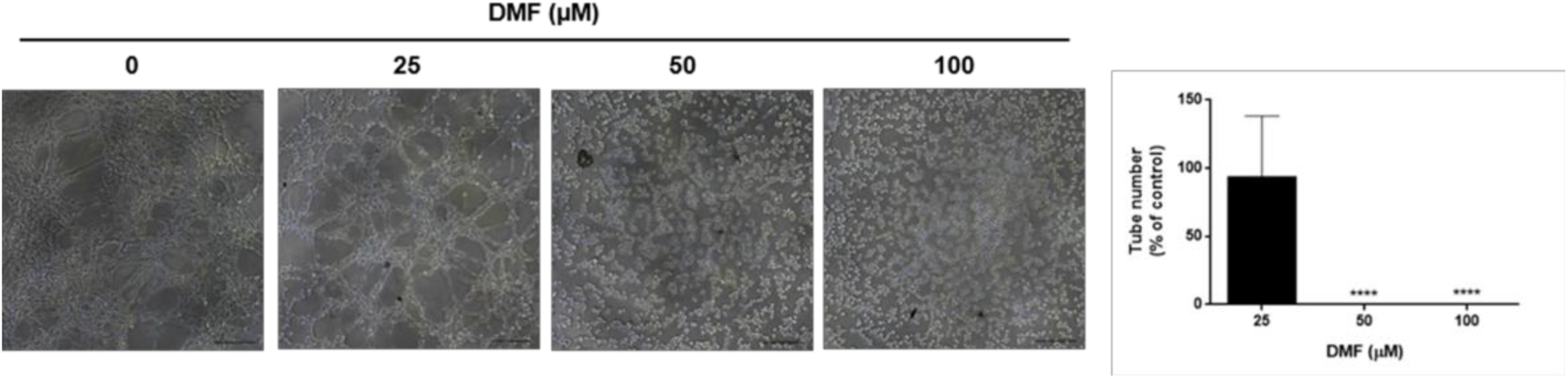
DMF inhibits tube formation in HMECs. Representative pictures and quantification of tube formation on Matrigel in HMECs treated with different concentrations of DMF. Bar scales = 183.25 μm. Data are expressed as means ± SD. ****p<0.0001 versus untreated control.

### DMF diminishes respiration while favors glycolysis in HMECs

As a first approximation to test the capacity of DMF to modulate global energetic metabolism in HMECs, oxygen consumption rate (OCR) and extracellular acidification rate (ECAR), as estimators of oxidative phosphorylation (OXPHOS) and glycolysis, respectively, were measured using a Seahorse flux analyzer. Obtained data showed that cells incubated for 20 h with 50 µM DMF had a higher glycolytic rate than control cells (p<0.001) (Figures 2A-B).

**Figure 2.**
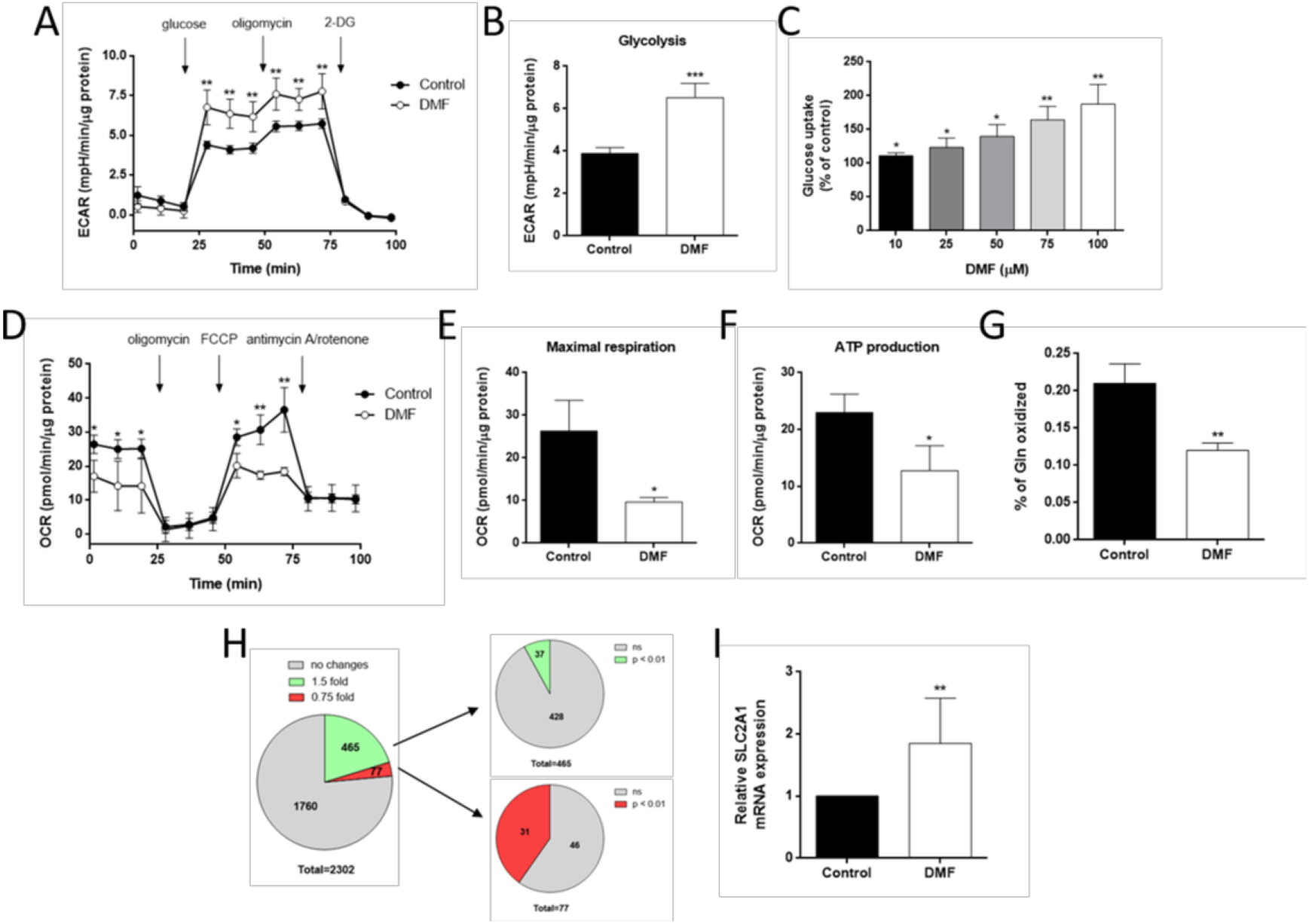
DMF increases glycolysis and diminishes OXPHOS in HMECs. (A) ECAR was measured in cells treated with 50 μM DMF and (B) glycolytic rate was calculated. (C) Glucose uptake after 30 minutes incubation with 5 mM glucose and 0.5 mM glutamine in cells treated with different doses of DMF for 16 h. (D) OCR was measured in cells treated with 50 μM DMF and (E) maximal respiration and (F) ATP production were calculated. (G) Glutamine oxidation after 30 minutes incubation with 5 mM glucose, 0.5 mM glutamine and 0.5 µCi/mL L-[U-^14^C]-glutamine after treatment with 100 μM DMF for 16 h. (H) Proteomics analysis of cells treated with 100 μM DMF for 24 h. (I) *SLC2A1* mRNA expression in cells treated with 100 μM DMF for 24 h. Data are expressed as means ± SD. *p<0.05; **p<0.01; ***p<0.001 versus untreated control.

Interestingly, this increased glycolytic rate observed in DMF-treated HMECs was correlated with an increased glucose uptake in these cells. HMECs cultured overnight with several concentrations of DMF were exposed to 30 minutes fasting in DPBS in the presence of DMF, followed by additional 30 minutes incubation in the presence of glucose and glutamine. Glucose taken up during those 30 minutes in the presence of DMF was measured, revealing that DMF increased glucose uptake in HMECs in a dose-dependent manner (p<0.05) (Figure 2C). Since 100 µM exerted a greater effect than 50 µM without compromising cell viability, we decided to keep using 100 µM DMF for further experiments.

In order to elucidate if the observed effect of DMF on glucose metabolism could be dependent on transcriptional regulation, we compared results of glucose uptake after overnight treatment with DMF with shorter incubation time (DMF added during the last 30 minutes incubation with glucose). Our data showed that 100 µM DMF incubated in short time (30 min before measurements) increased glucose uptake in HMECs to a lesser extent than after overnight incubation, yet the increase was statistically significant in both conditions (p<0.05) (116,2 ± 4,2% and 187,6 ± 29,1% data of fold compared to the control condition in short time and overnight DMF treatments, respectively). These results suggest the implication of transcriptional regulation in the DMF-enhanced glucose metabolism in ECs, although additional shorter-term mechanisms may also contribute to this effect.

Additionally, we determined that OCR was lower in DMF-treated cells (Figures 2D-F). Since intracellular glutamine is mainly incorporated into the TCA and oxidized, we tested glutamine oxidation in HMECs in presence of 100 µM DMF after overnight incubation, using the same experimental conditions described in glucose uptake assays. As shown in Figure 2G, 100 µM DMF halved glutamine oxidation in HMECs (p<0.01), indicating an effect in the metabolic use of this amino acid.

Previously described experiments were performed in nutrient-limited conditions, assuring the detection of direct effects of DMF on the metabolic substrates of interest. On the other hand, in order to assess glucose and glutamine uptake, as well as lactate, glutamate and ammonia secretion in a more complex mixture of nutrients, we cultured cells overnight or for 24 h incubation in the presence of 100 µM DMF in complete medium. Either way, glucose uptake and lactate secretion were increased in HMECs treated with DMF (p<0.01) (Figure S1A-B). On the contrary, DMF reduced glutamine uptake in HMECs (p<0.05), whereas glutamate release to the medium was slightly higher in treated cells (p<0.001) (Figure S1C-D). Regarding ammonia production, no differences were found in presence of DMF (Figure S1C-D). All these results point to a DMF-mediated upregulation of glycolysis in HMECs, whereas oxidative metabolism seems compromised in presence of this compound. Obtained results were similar in both timepoints, and hence 24 h incubation was preferred for next experiments.

### DMF has differential effects in different cell lines

In order to see whether the observed effects of DMF were specific to HMECs, we also tested glucose uptake and glutamine oxidation in different cell lines treated with DMF, including macro- and mesovascular ECs (BAECs and HUVECs), two breast adenocarcinoma cell lines (MDA-MB-231 and MCF7), a cervix adenocarcinoma cell line (HeLa) and fibroblasts (HGF), in order to cover non-microvascular ECs, different tumor cell lines and a non-transformed cell line different from endothelium.

As shown in Figure S2A, the effect of 100 µM DMF overnight incubation on glucose uptake was also found in all the tested cell lines (p<0.05), suggesting that DMF could be targeting a common mechanism in all of them. Regarding glutamine oxidation, a slight inhibitory effect, yet not statistically significant, was found in HUVECs after DMF treatment, whereas this DMF-induced reduction was significant in tumor MDA-MB-231 cells (p<0.05) and no effect was found in HeLa cells (Figure S2B).

### DMF upregulates GLUT1 expression without affecting HIF-1α

Due to the greater effect of DMF on glucose uptake after longer incubation, we hypothesized that this compound might modulate glucose and/or glutamine metabolism through modulation of gene and/or protein expression. To test this premise, a quantitative proteomics analysis was performed in samples from HMECs treated with 100 µM DMF for 24 h. We considered an upregulation on protein expression when at least a 1.5-fold increase in the abundance ratio (DMF/DMSO) was found and a downregulation of those proteins with a 0.75-fold or lower expression in the abundance ratio (DMF/DMSO). A total of 2302 proteins were identified with a high confidence level and at least two peptides detected. Of those, 465 presented a ≥1.5-fold increase and 77 a ≤0.75-fold expression. However, we only considered statistically significant those changes with a p-value lower than 0.01, thus selecting 37 upregulated proteins and 31 downregulated after DMF treatment (Figure 2H and Tables S1-S3).

Among the upregulated proteins, glucose transporters GLUT1 and GLUT14, also known as solute carrier family 2, facilitated glucose transporter member 1 (*SLC2A1*) and member 14 (*SLC2A14*), respectively, expressions were found to be 3.72-fold and 4.06-fold compared to the control condition, respectively (p<0.01) (Table S2). Moreover, DMF also increased mRNA *SLC2A1* expression in these cells (Figure 2I). Since GLUT1 is under the transcriptional control of HIF-1α, and DMF was shown to stabilize HIF-1α in human embryonic kidney cells, we checked HIF-1α protein levels in HMECs treated with DMF (Koivunen et al., 2007). However, we did not detect any HIF-1α in normoxia with DMF (data not shown). Thus, this increase in GLUT1 expression was not likely the consequence of a stabilization of HIF-1α in normoxia in the presence of DMF, discarding this possibility and pointing to a different mechanism of action responsible of the observed increase in glucose transporters expression induced by DMF.

### DMF affects aspartate and TCA cycle intermediates levels

Next, we performed steady-state metabolomics in complete medium in order to study the possible changes in the intracellular pool of several metabolites as a consequence of the deregulated glycolytic and oxidative metabolism in DMF-treated cells. Among other changes, we observed that aspartate levels were drastically lower in DMF-treated HMECs (p<0.0001) (Figure 3A and Fig. S3). Of note, aspartate is absent in DMEM formulation, which we used for these experiments, and hence cells need to synthetize it. However, we also performed this experiment in cells cultured in RPMI-1640 medium, which contains aspartate and, in these conditions, aspartate levels in DMF-treated cells were also lower, but to a lesser extent than when cells were cultured in aspartate-free medium (data not shown). Interestingly, aspartate synthesis is regulated by the electron transport chain (ETC) activity (Birsoy et al., 2015). Since DMF treatment suppressed respiration and glutamine oxidation in HMECs, we also performed stable isotope-labeling studies using glutamine labeled with carbon-13 in its five carbons ([U-^13^C]-glutamine) to follow the labeling of TCA intermediates (Figure 3B). As expected, labeling of aspartate, fumarate, malate and citrate from glutamine was lower in DMF-treated cells (p<0.05) (Figures 3C-F), corroborating a lower incorporation of glutamine into the TCA cycle. The fumarate pool was not affected (Figure 3G), probably due to external addition of fumarate from DMF. Intracellular malate levels were lower (p<0.0001) (Figure 3H). However, citrate levels were increased after DMF treatment (p<0.001) (Figure 3I), which may reflect inhibition of the TCA cycle.

**Figure 3.**
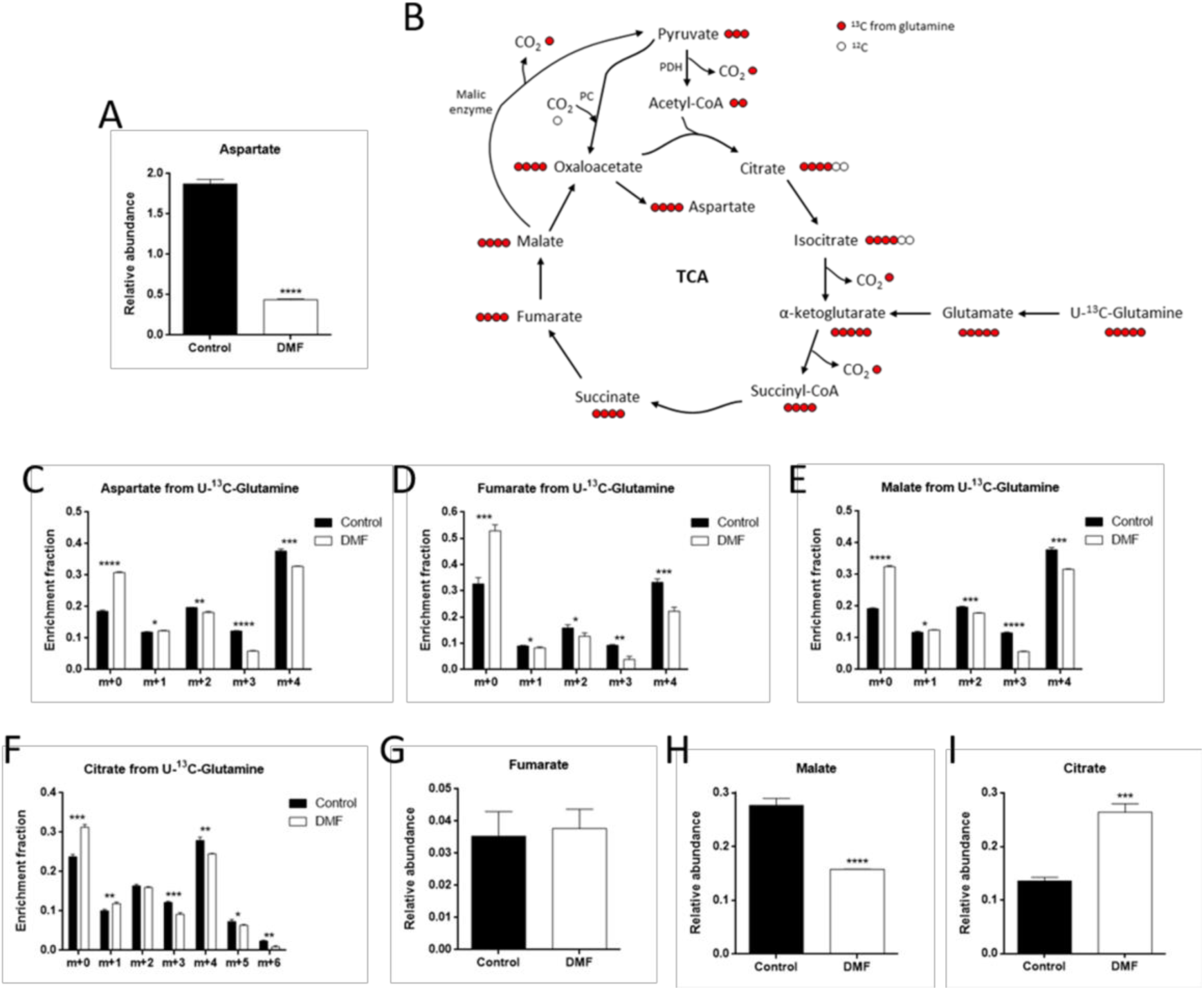
DMF diminishes incorporation of glutamine into the TCA cycle in HMECs. (A) Intracellular aspartate levels in cells treated with 100 μM DMF for 24 h. (B) Scheme of TCA cycle illustrating labeling from [U-^13^C]-glutamine. (C) Fractional labeling of aspartate, (D) fumarate, (E) malate and (F) citrate from [U-^13^C]-glutamine in cells treated with 100 μM DMF for 24 h. (G) Intracellular fumarate, (H) malate and (I) citrate levels in cells treated with 100 μM DMF for 24 h. Data are expressed as means ± SD. *p<0.05; **p<0.01; ***p<0.001; ****p<0.0001 versus untreated control.

### DMF decreases serine and glycine synthesis and favors extracellular serine and glycine uptake in HMECs

Interestingly, by means of the steady-state metabolite experiment, higher levels of intracellular glycine were found in DMF-treated cells (p<0.001) (Figure 4A and Fig. S2). Serine levels were slightly higher, yet not statistically significant (Figure 4B). Again, we performed stable-isotope-labeling studies, this time using glucose labeled at all six carbons with carbon-13 ([U-^13^C]-glucose), in order to check whether serine and glycine synthesis from glucose was boosted in the presence of DMF. The endogenous synthesis of serine and glycine starts from glucose, which through glycolysis, converts after several steps into 3-phosphoglycerate (3-PG). This glycolytic intermediate is the substrate of phosphoglycerate dehydrogenase (PHGDH). The resultant 3-phosphohydroxypyruvate (PHP) is then converted into 3-phosphoserine (P-Ser) through phosphoserine aminotransferase (PSAT), and this P-Ser is finally the substrate of phosphoserine phosphatase (PSPH), resulting in the synthesis of serine. Finally, glycine is the product of the enzyme serine hydroxymethyltransferase (SHMT) from serine (Figure 4C).

**Figure 4.**
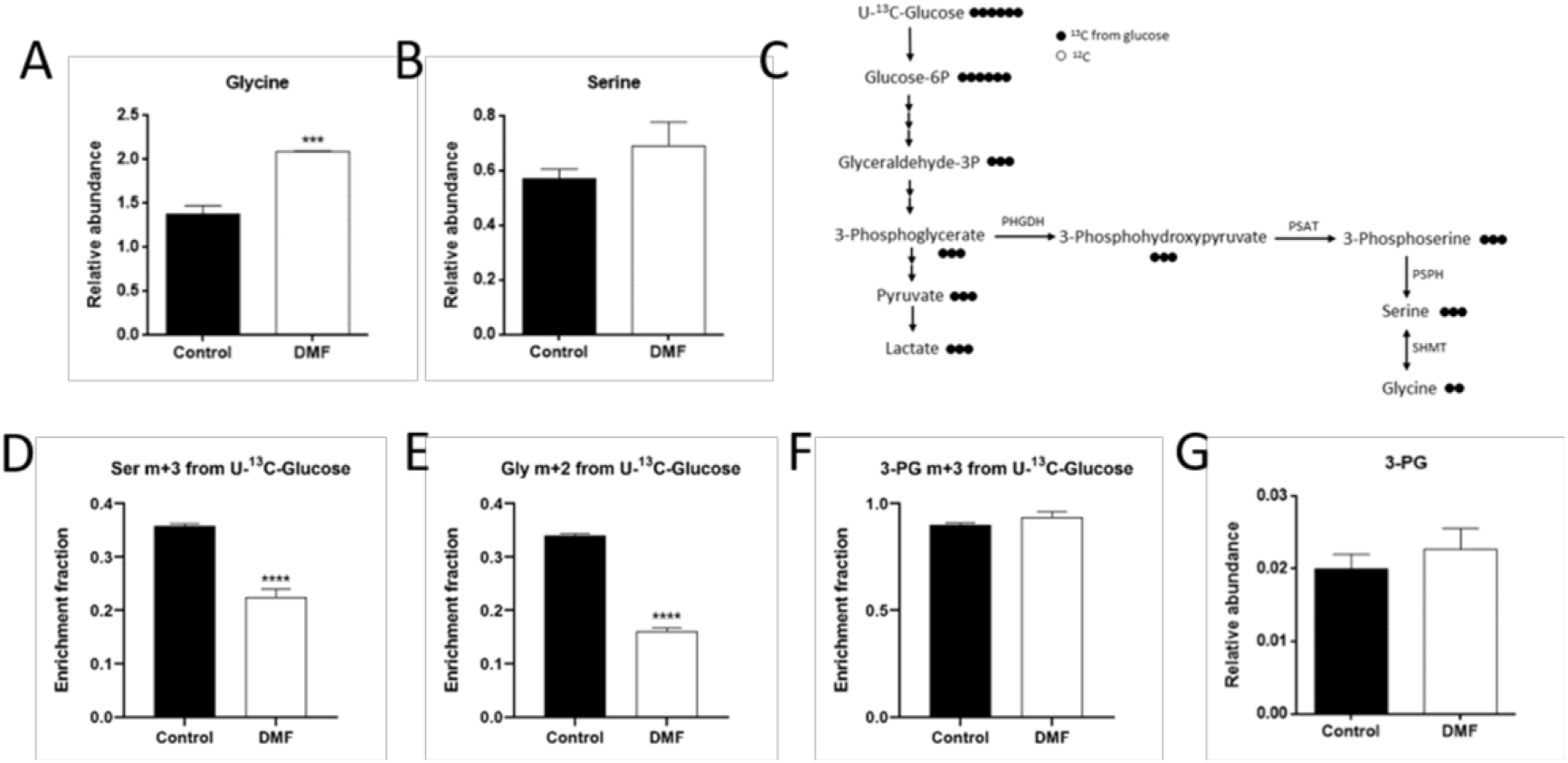
DMF decreases serine and glycine synthesis from glucose in HMECs. (A) Intracellular glycine and (B) serine levels in cells treated with 100 μM for 24 h. (C) Scheme of glycolysis and the serine and glycine synthesis pathway illustrating labeling from [U-^13^C]-glucose. (D) Fractional labeling of serine, (E) glycine and (F) 3-PG from [U-^13^C]-glucose in cells treated with 100 μM DMF for 24 h. (G) Intracellular 3-PG levels in cells treated with 100 μM DMF for 24 h. Data are expressed as means ± SD. ***p<0.001; ****p<0.0001 versus untreated control.

Surprisingly, we found that not only serine m+3 from glucose was lower after 24 h incubation with 100 µM DMF (p<0.0001) (Figure 4D), but an even greater decrease in glycine m+2 was detected (p<0.0001) (Figure 4E). No changes in 3-PG m+3 labeling or intracellular levels were found (Figures 4F-G), which could have been expected due to the higher glycolytic activity in DMF-treated cells. Similar effects were found in HMECs treated with 100 µM DMF overnight or with 50 µM DMF for 24 h (Figures S4 and S5).

Due to the presence of extracellular serine and glycine in the medium we used for metabolomics and isotope tracing analysis, we checked whether depleting serine and glycine from the medium affected metabolism of HMECs. Our data showed that serine and glycine withdrawal did not affect the observed effect of DMF on glucose uptake or lactate production in HMECs (neither level in control conditions) compared to conditions with extracellular serine and glycine (Figure S6).

Regarding serine and glycine synthesis from glucose, on the one hand serine m+3 was higher in control and DMF-treated HMECs incubated in the absence of both serine and glycine (p<0.0001) (Figure 5A), indicating the need for serine synthesis when no extracellular serine is available. Serine pools were higher in DMF-treated cells when serine and glycine were present in the medium (p<0.001), whereas serine levels were low in serine and glycine depleted medium in both conditions (p<0.001) (Figure 5B). Glycine m+2 was also higher in control HMECs when no extracellular serine and glycine was available (p<0.01) (Figure 5C). However, DMF-treated cells failed to increase glycine m+2 labeling from glucose in the absence of these two amino acids to the same extent as they did with serine (p<0.01 compared to cells treated with DMF in the presence of serine and glycine) (Figure 5C). Furthermore, intracellular glycine levels were increased in DMF-treated cells when there was serine and glycine in the medium (p<0.001), but the glycine pool was almost totally depleted during serine and glycine withdrawal (p<0.01) (Figure 5D). No remarkable changes were found in either 3-PG labeling or intracellular pool between control and DMF-treated cells in conditions with or without extracellular serine and glycine (Figures 5E-F). Together, these results suggest that DMF-treated ECs had their *de novo* synthesis pathway compromised, while the intracellular pool of these two amino acids was higher compared to control cells.

**Figure 5.**
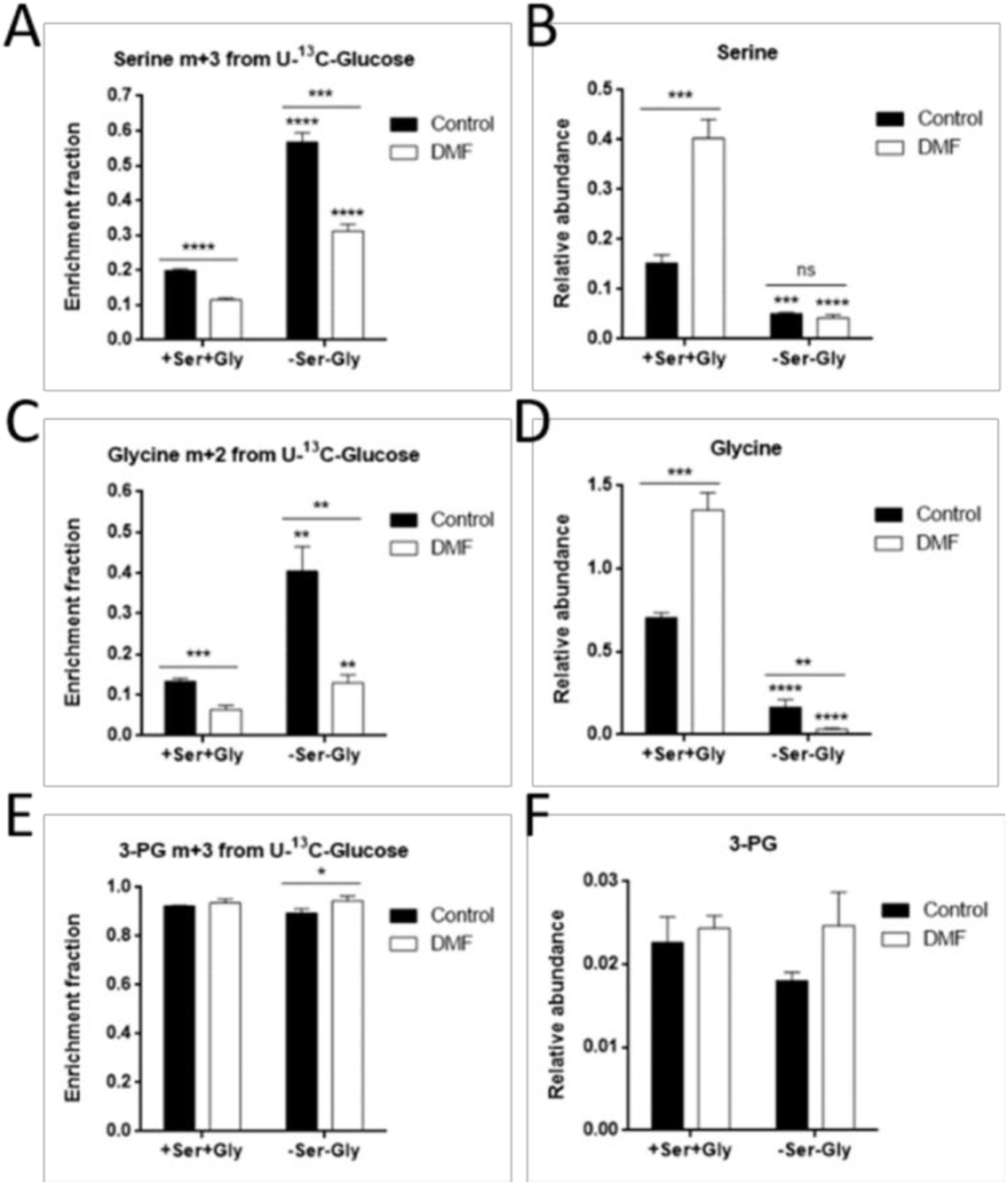
DMF favors serine and glycine synthesis from glucose in the absence of extracellular serine and glycine in HMECs. (A) Fractional labeling of serine from [U-^13^C]-glucose, (B) intracellular serine levels, (C) fractional labeling of glycine from [U-^13^C]-glucose, (D) intracellular glycine levels, (E) fractional labeling of 3-PG from [U-^13^C]-glucose and (F) intracellular 3-PG levels in cells treated with 100 μM DMF for 24 h in the presence or absence of extracellular serine and glycine. Data are expressed as means ± SD. *p<0.05; **p<0.01; ***p<0.001; ***p<0.0001 versus condition with extracellular serine and glycine. ns: non-significant.

Therefore, we next checked if an increase in extracellular serine and glycine uptake was taking place in DMF-treated cells. For that aim, we used serine and glycine labeled with carbon-13 in all their carbons ([U-^13^C]-serine and [U-^13^C]-glycine). In order to avoid interferences, extracellular glycine was absent in medium supplemented with labeled serine, whereas serine was not added to the medium with labeled glycine, since these two amino acids can be converted into each other through SHMT activity (Figure 6A). Not surprisingly, HMECs treated with 100 µM DMF presented higher serine m+3 labeling from labeled serine (p<0.0001) (Figure 6B). Glycine m+2, which comes from this extracellular labeled serine, was also higher after DMF treatment (p<0.05) (Figure 6C). Nevertheless, the contribution of labeled glycine to the intracellular glycine pool was not as high in DMF-treated cells respect to the control condition compared to serine uptake (p<0.05) (Figure 6D). No significant differences were found in serine labeling from glycine (data not shown). These data indicate that uptake of these two amino acids is increased in DMF-treated HMECs.

**Figure 6.**
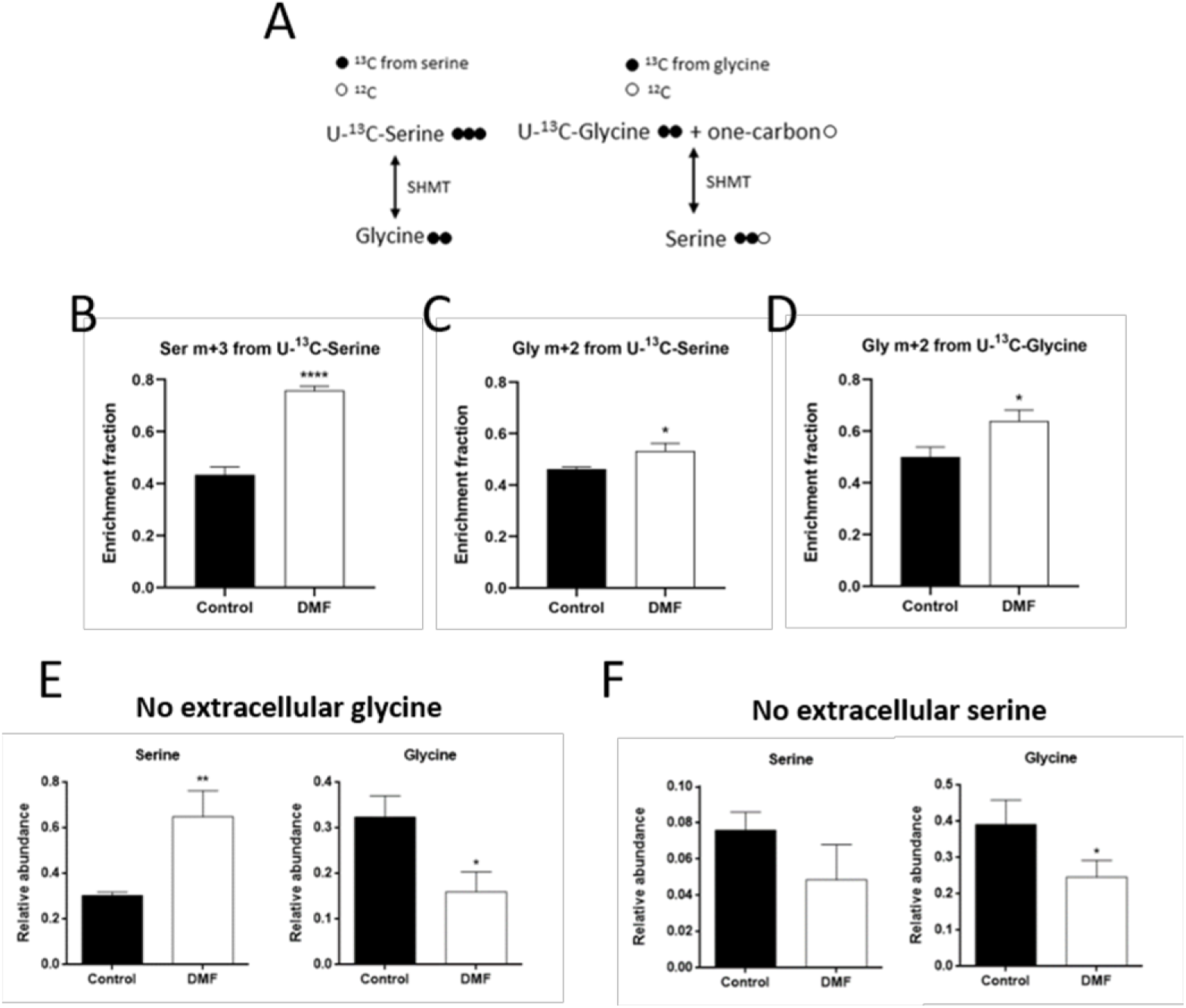
DMF favors extracellular serine and glycine uptake in HMECs. (A) Scheme of serine and glycine interconversion illustrating labeling from [U-^13^C]-serine and [U-^13^C]-glycine. (B) Fractional labeling of serine and (C) glycine from [U-^13^C]-serine, and of (D) glycine from [U-^13^C]-glycine in cells treated with 100 μM DMF for 24 h. (E) Intracellular serine and glycine levels in medium without glycine or (F) without serine in cells treated with 100 μM for 24 h. Data are expressed as means ± SD. *p<0.05; **p<0.01; ***p<0.0001 versus untreated control.

Regarding intracellular serine and glycine pools, when serine, but not glycine, was present in the medium, intracellular serine levels were higher in DMF-treated cells (p<0.01), but glycine levels were lower (p<0.05) (Figure 6E), indicating an increase in extracellular serine uptake, which could not be converted to glycine. However, depleting serine from the medium while extracellular glycine is available diminished serine levels after DMF treatment, yet not significantly, whereas the glycine pool was unexpectedly decreased (p<0.05) (Figure 6F).

### DMF down-regulated PHGDH activity without affecting protein levels

So far, our data indicated a major role of serine and, specially, glycine synthesis in HMECs compared to their extracellular uptake, pointing to DMF as an inhibitor of this biosynthetic pathway. This was surprising, since our proteomics analysis revealed a 4.5-fold expression of PSPH, the third and last enzyme in the synthesis pathway (p<0.0001) (Table S2). However, the rate-limiting enzyme under cell culture conditions is PHGDH, which catalyzes the committed step in the serine synthesis pathway (Scerri et al., 2017). Based on proteomics results, PHGDH protein expression was not significantly changed in DMF-treated ECs (Table S1) and we further validated this data by Western-blot, assessing that PHGDH protein expression was unaffected by DMF treatment in HMECs (Figures 7A-B). Nonetheless, treatment with 100 µM DMF decreased PHGDH activity in HMECs (p<0.0001) (Figure 7C), and this partial inhibition matches the percentage of serine m+3 labeling from glucose (Figure 4D). Therefore, even if PSPH protein levels are higher with DMF, a lower PHGDH activity is most likely limiting the serine and glycine synthesis rate in HMECs. Interestingly, DMF failed to decrease PHGDH activity in cell lysates from the control condition (1.30 ± 0.13 mU/mg *vs*. 1.32 ± 0.13 mU/mg in cell lysates from control cells treated *on site* with DMSO or 100 µM DMF, respectively). These data suggest that the PHGDH inhibition exerted by DMF in ECs was not a direct effect on the enzyme, and was probably dependent on some cellular processes.

**Figure 7.**
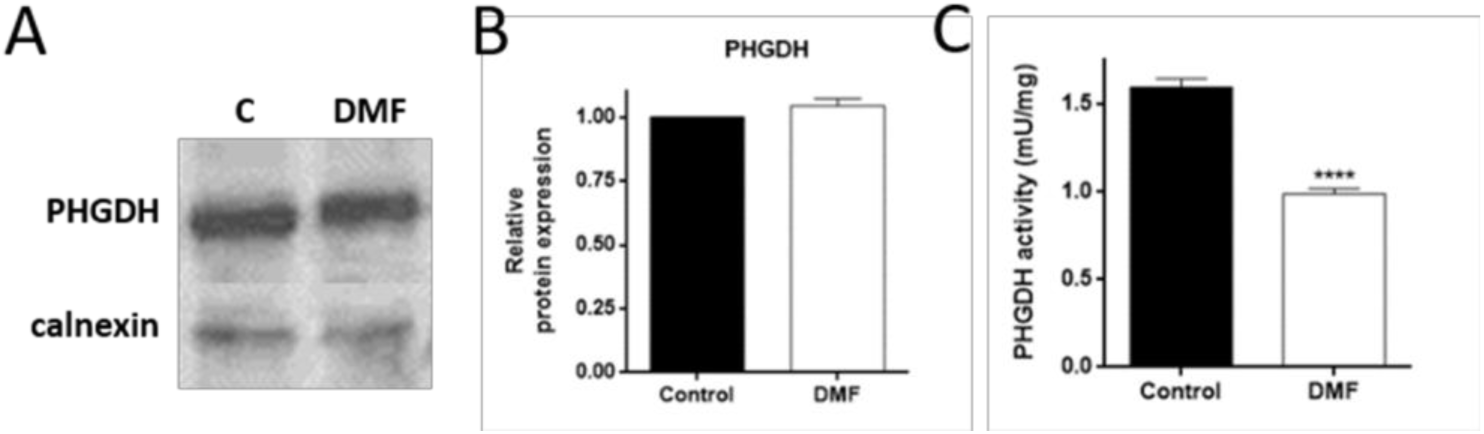
DMF inhibits PHGDH activity in HMECs. (A) Representative Western blot for PHGDH and (B) quantification in cells treated with 100 μM DMF for 24 h. (C) PHGDH activity in cells treated with 100 μM for 24 h. Data are expressed as means ± SD. ****p<0.0001 versus untreated control.

## Discussion

Study of EC metabolism has gained importance in the search of new targets for the treatment of angiogenesis-dependent diseases (Goveia et al., 2014; Ocana et al., 2019b). As far as we are concerned, this is the first work demonstrating the reduction on PHGDH activity by the anti-angiogenic compound DMF. The serine synthesis pathway has been described to be essential for the progression of several types of tumors, and the development of PHGDH inhibitors has emerged as a promising cancer therapy (Ravez, Spillier, Marteau, Feron, & Frederick, 2017). One of the most studied pathways regulated by DMF is the Nrf2 pathway. Interestingly, Nrf2 has been shown to induce protein expression of genes from the serine and glycine synthesis pathway, including PHGDH, through ATF4 in cancer cells (DeNicola et al., 2015). These results are converse to those obtained with ECs. Other studies have reported a transcriptional regulation of PHGDH by other molecules also affected by DMF (Liu et al., 2017; Ou, Wang, Jiang, Zheng, & Gu, 2015). Nevertheless, our results suggest that the effects of DMF on EC energetic metabolism are not likely controlled by a transcriptional regulation but by direct enzymatic inhibition by an unknown mechanism which is likely to require some cellular process.

DMF is known to be an electrophilic molecule able to bind to protein cysteine residues in a process called succination, hence modifying their activity (Frycak, Zdrahal, Ulrichova, Wiegrebe, & Lemr, 2005). Indeed, many of the DMF molecular targets suffer cysteine succination (Saidu et al., 2019). This fact makes studying the exact mechanism of action of this compound a big challenge due to this lack of specificity, which could affect many molecules through a cascade of different dysfunctional signaling pathways. Interestingly, a global analysis of cysteine ligandability performed in cancer cells revealed that Cys369 of PHGDH can react with electrophilic small molecules (Bar-Peled et al., 2017). Whether the modulation of PHGDH activity exerted by DMF involves succination of cysteine residues of this protein remains unstudied. However, since DMF failed to inhibit PHGDH activity in control cell lysates *in vitro*, it is likely that some cellular process participates in the regulation of PHGDH activity mediated by DMF and that cysteine succination might not be enough for repressing the PHGDH activity in ECs.

Our data point out that suppression of PHGDH activity in ECs seems to have a greater effect on glycine synthesis compared to serine synthesis, since glycine levels were more compromised after DMF treatment than the serine pool even when extracellular glycine was available. It is known that *de novo* mitochondrial glycine synthesis is highly active in ECs and more important than cellular uptake (Hitzel et al., 2018). Conversely, extracellular glycine has been shown to stimulate VEGF signaling and angiogenesis *in vitro* and *in vivo* by promoting mitochondrial function, and VEGF was found to promote the expression of the glycine transporter GlyT1 in ECs, while it did not affect the levels of the enzymes involved in glycine synthesis (Guo et al., 2017). DMF was reported ten years ago to exert an anti-angiogenic activity *in vitro* and *in vivo*, at least partially due to VEGFR2 suppression (Garcia-Caballero et al., 2011; Meissner et al., 2011). The exact mechanism by which DMF represses VEGFR2 expression remains unexplored and would require further research. Since DMF inhibits VEGFR2 and, therefore, the VEGF signaling pathway in ECs, it could be possible that glycine uptake is suppressed in DMF-treated ECs. In any case, both extracellular glycine and *de novo* synthetized glycine seem important to EC metabolism and function, and DMF plays a major role in glycine metabolism through PHGDH inhibition. Nonetheless, the existence of an interplay between the DMF-induced PHGDH inhibition and the anti-angiogenic activity of this molecule are related remains unclear.

Among other metabolic pathways, glycolysis has been described to be essential for vessel sprouting (De Bock et al., 2013). Some years ago, DMF was found to inhibit glycolysis in immune cells through targeting of GAPDH activity mediated by succination of several cysteine residues (Kornberg et al., 2018). Conversely, the results found in this work show that DMF increases glycolysis whereas diminishes OXPHOS in ECs. Why this compound affects energetic metabolism differently in different cell types has yet to be answered. Remarkably, using ECs, Vandekeere and colleagues found out that silencing PHGDH impaired angiogenesis, even when the glycolytic rate of PHGDH knock-down cells was higher than those whose levels of PHGDH remained intact (Vandekeere et al., 2018). These results are similar to those obtained in this work for the inhibition of PHGDH after DMF treatment in HMECs. Thus, it is likely that PHGDH inhibition boosts glycolytic activity in order to compensate the reduction in serine synthesis rate. However, we found an increased glucose uptake in several cell lines treated with DMF, including MDA-MB-231, a triple negative breast cancer cell line which lacks PHGDH (Possemato et al., 2011). Therefore, additional mechanisms must regulate the increase in the glycolytic rate exerted by DMF.

It is worthy of note the different effects of DMF on glutamine oxidation in different cell lines. DMF diminished glutamine oxidation in different EC lines and in cancer MDA-MB-231 cells. All these cell lines are known to be highly glycolytic (Gaglio et al., 2011; Ocana, Martinez-Poveda, Quesada, & Medina, 2019a; Peters et al., 2009). However, DMF failed to alter glutamine oxidation in the highly glutamine-dependent, oxidative cell line HeLa (Reitzer, Wice, & Kennell, 1979). This differential effect points out to a different regulation of energetic metabolism depending on the metabolic preferences of the cells. This makes an interesting point in using DMF as a therapeutic tool in different pathological contexts, since this compound may exert different effects in different cell types such as ECs, cancer cells and immune cells.

Altogether, our data suggest a complex regulation of EC metabolism by DMF (Figure 8). Elucidating the exact mechanism of action of DMF requires a vast and comprehensive experimental design, due to its reported high molecular reactivity and wide regulation of transcription factors, such as Nrf2 and NF-κB. Thus, it remains unclear whether impairment of angiogenesis through inhibition of VEGFR2 and the decrease in serine and glycine synthesis pathway by reduced PHGDH activity exerted by DMF are causally related. Although glycolysis has been shown to be essential for angiogenesis, inhibition of PHGDH impaired the angiogenic process but increased the glycolytic rate in ECs (De Bock et al., 2013; Vandekeere et al., 2018). Therefore, the results published by Vandekeere *et al*. and the data presented in this work reinforces the complexity of metabolic regulation in ECs and its relation with the angiogenic switch.

**Figure 8.**
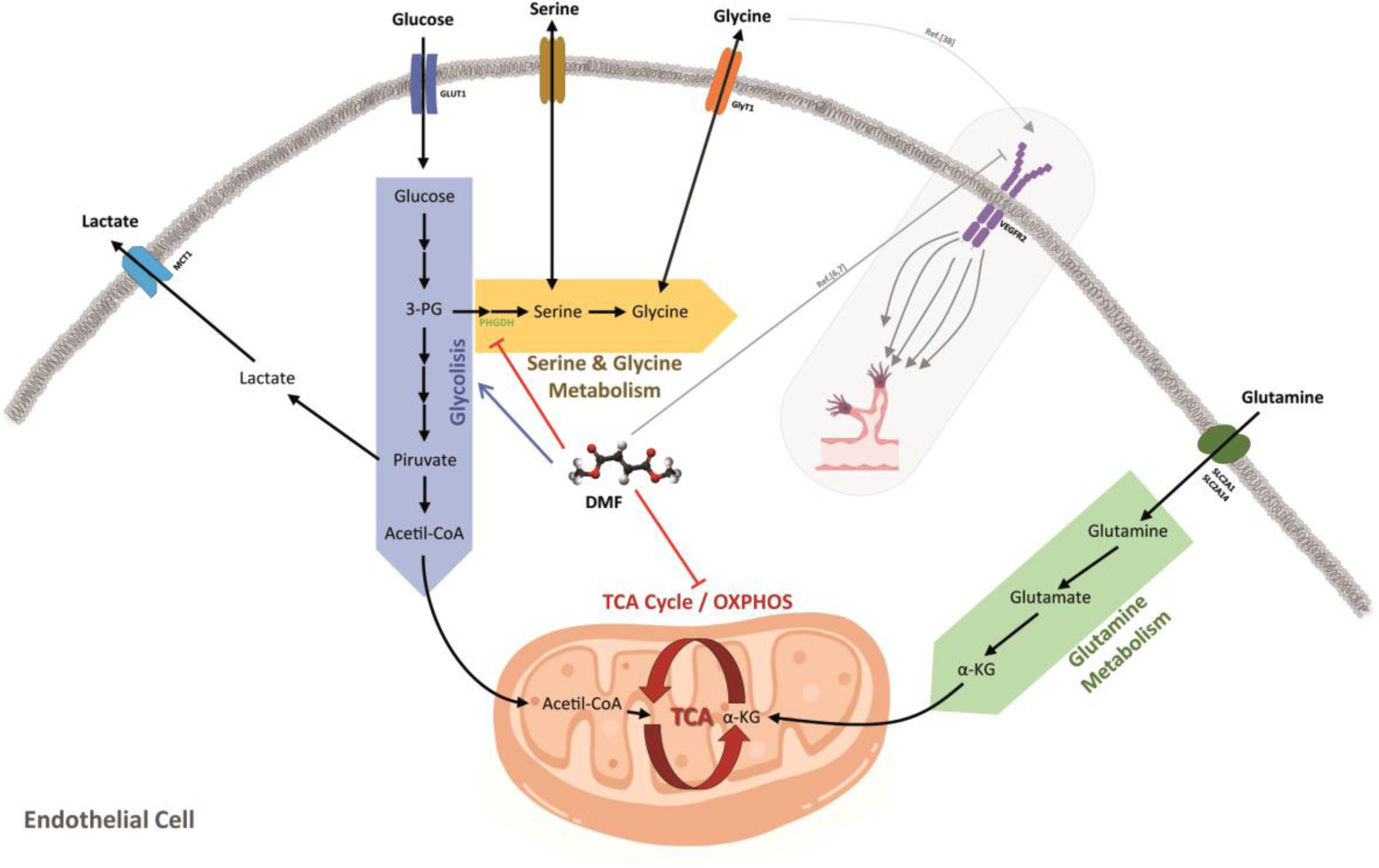
Summary of DMF effects on energetic EC metabolism. The results of this work show how DMF induces glucose uptake through upregulation of GLUT1 expression. This increase in glucose uptake leads to a greater glycolytic rate and lactate secretion to the extracellular media. Conversely, glutamine uptake is diminished in cells treated with DMF, and its oxidation through the TCA cycle is reduced. DMF diminished PHGDH activity and synthesis of serine and glycine from glucose. It is described in the bibliography that DMF inhibits VEGFR2 expression, and that extracellular glycine has been seen to promote VEGF signaling and angiogenesis through upregulation of mitochondrial activity. Whether inhibition of glycine metabolism is related with the anti-angiogenic activity of DMF remains unclear.

## Materials and methods

### Materials

MCDB-131 cell culture medium was obtained from Gibco (Paisley, Scotland, UK). Glucose, glutamine, serine and glycine free media were from Teknova (Hollister, CA, USA) and from US Biological Life Sciences (Salem, MA, USA). Other cell culture media, penicillin and streptomycin and trypsin were purchased from BioWhittaker (Verviers, Belgium). Fetal bovine serum (FBS) was purchased from Biowest (Kansas, USA). Dialyzed FBS (dFBS) was from Gemini Bioproducts (West Sacramento, CA, USA) and from Capricorn (Ebsdorfergrund, Germany). Matrigel was purchased from Becton-Dickinson (Bedford, MA, USA). Material for Seahorse experiments were from Agilent Technologies (Santa Clara, CA, USA). 2-NBDG was supplied by Molecular Probes (Eugene, OR, USA). L-[U-^14^C]-Glutamine was acquired from Perkin Elmer (Waltham, MA, USA). L-glutamine/ammonia assay kit was from Megazyme (Bray, County Wicklow, Ireland). Anti-HIF1α antibody was from BD Biosciences (San Jose, CA, USA), anti-α-tubulin antibody was from Cell Signaling Technology (Danvers, MA, USA), anti-PHGDH antibody was from GeneTex (Irvine, CA, USA) and anti-calnexin was from Enzo Life Sciences (Farmingdale, NY, USA). D-[U-^13^C]-Glucose, L-[U-^13^C]-glutamine, L-[U-^13^C]-serine and L-[U-^13^C]-glycine were purchased from Cambridge Isotope Laboratories (Tewksbury, MA, USA). PHGDH activity assay kit was from BioVision (Milpitas, CA, USA). Plastic material for cell culture was from Nunc (Roskilde, Denmark). All other reagents not listed on this section, including DMF and glucose and glutamine free medium, were from Sigma-Aldrich (St. Louis, MO, USA).

### Cell culture

All cell culture media, unless otherwise specified, were supplemented with glutamine (2 mM), penicillin (50 U/mL) and streptomycin (50 U/mL). Human microvascular endothelial cells (HMECs) were kindly supplied by Dr. Arjan W. Griffioen (Maastricht University, Netherlands) and maintained in MCDB-131 medium supplemented with 10% FBS, hydrocortisone (1 μg/mL) and EGF (10 ng/mL). Human umbilical vein endothelial cells (HUVECs) were isolated by a modified collagenase treatment as previously reported and maintained in 199 medium supplemented with 20% fetal bovine serum, ECGS (30 µg/mL) and heparin (100 µg/mL) (Kubota, Kleinman, Martin, & Lawley, 1988). Bovine aortic endothelial cells (BAECs) were isolated from bovine aortic arches as previously described and maintained in Dulbecco’s modified Eagle’s medium (DMEM) containing glucose (1 g/L) and supplemented with 10% FBS (Cardenas, Quesada, & Medina, 2006). Primary human gingival fibroblasts (HGF) were maintained in DMEM containing glucose (4.5 g/L) and 10% FBS. Tumor cells used in this paper (human breast carcinoma MDA-MB-231 and MCF7, and human cervix adenocarcinoma HeLa) were purchased from the ATCC (Rockville, MD, USA) and maintained in RPMI-1640, DMEM containing glucose (4.5 g/L) and EMEM, respectively, all of them supplemented with 10% FBS. All cell lines were maintained at 37 °C under a humidified 5% CO2 atmosphere.

### Tube formation on Matrigel by endothelial cells

Each well of a 96-well plate was coated with Matrigel (50 µL of about 10.5 mg/mL) at 4 °C and polymerized at 37 °C for a minimum of 30 min. 7 x 10^4^ cells were seeded in 200 µL of medium without serum. 25, 50 and 100 µM DMF were added to the wells and incubated at 37 °C. After 5 hour incubation, cultures were observed and photographed with a microscope camera Nikon DS-Ri2 coupled to a Nikon Eclipse Ti microscope from Nikon (Tokyo, Japan). Closed “tubular” structures were counted using ImageJ software.

### Extracellular flux analyzer experiments

HMECs were cultured at a density of 3 x 10^4^ cells/well in 24-well Seahorse XFe24 plates (Agilent) and incubated at 37 °C under a humidified 5% CO2 atmosphere overnight in the presence or absence of DMF. Cells were washed twice with XF base medium (Agilent) and incubated with XF medium containing or not DMF and supplemented with 10 mM glucose and 4 mM glutamine or just with 4 mM glutamine at 37 °C without CO2 for one hour. Three initial measurements were made using XFe24 Seahorse analyzer. Then, 1 µM oligomycin, 0.6 µM carbonyl cyanide-4-(trifluoromethoxy)phenylhydrazone (FCCP), and 1 µM antimycin A and 1 µM rotenone for wells with both glucose and glutamine, or 10 mM glucose, 1 µM oligomycin and 25 mM 2-deoxyglucose (2-DG) for wells with only glutamine, were injected sequentially to each well with three measurements between each injection. Data were analyzed using Wave software and normalized to protein amount.

### Measurement of glucose, glutamine, lactate, glutamate and ammonia

For glucose uptake after short incubation time, cells cultured in 96-well plates were treated with DMF overnight. Then, cells were washed twice with PBS supplemented with calcium and magnesium (DPBS), and then starved for 30 min with this DPBS containing or not DMF. Cells were incubated for additional 30 min with DPBS containing or not DMF and supplemented with 5 mM glucose, 0.5 mM glutamine and 100 µM 2-NBDG (2-(N-(7-Nitrobenz-2-oxa-1,3-diazol-4-yl)Amino)-2-Deoxyglucose). Relative glucose uptake was determined using a FACS VERSE^TM^ cytometer from BD Biosciences (San Jose, CA, USA) as previously described (Zou, Wang, & Shen, 2005). Data were analyzed with BD FACSuite software. For longer incubation times, concentrations of extracellular glucose, lactate, glutamine, and glutamate of control and DMF-treated cells were determined from aliquots of media using an automated electrochemical analyzer (BioProfile Basic-4 analyzer; NOVA). Ammonia secretion was measured from a fraction of the same aliquots with a spectrophotometric assay (Megazyme) using a FLUOstar Omega microplate reader from BMG LABTECH (Ortenberg, Germany). Data were normalized to cell number.

### Measurement of glutamine oxidation

Cells cultured in 24-well plates and treated or not with 100 µM DMF overnight were washed twice with PBS supplemented with DPBS, and then starved for 30 min with this DPBS containing or not DMF. Cells were incubated for additional 30 min with DPBS containing or not DMF and supplemented with 25 mM HEPES, 0.5 mM glutamine, 5 mM glucose and 0.5 µCi/mL L-[U-^14^C]-glutamine. Media and cells were collected in round-bottom glass tubes with screw-caps. Each glass tube contained a Whatman^TM^ paperfold imbibed with benzethonium hydroxide (hyamine). 400 µL 10% (v/v) HClO4 were added to the closed glass tubes through the cap. Tubes were incubated for additional 30 minutes at 37 °C with agitation. Whatman^TM^ paperfolds with ^14^CO2 captured in them were mixed with scintillation liquid. A Beckman Coulter LS6500 liquid scintillation counter (Fullerton, CA, USA) was used for the measurements. All assays were performed at the Radioactive Facilities of the University of Málaga, authorized with reference IR/MA-13/80 (IRA-0940) for the use of non-encapsulated radionuclides.

### Proteomics analysis

Control and DMF treated cells were extensively washed and frozen for their analysis. Cells were lysed using RIPA buffer on ice for 5 minutes and scratched. Cell extracts were sonicated and centrifuged and supernatants were collected. For protein precipitation, a modified trichloroacetic acid–acetone precipitation method (Clean-Up Kit; GE Healthcare, Munich, Germany) was used. The resultant precipitate was dissolved in bidistilled water, sonicated and centrifuged and supernatants were collected in a clean tube. Then we carried out a gel-assisted proteolysis entrapping the protein solution in a polyacrylamide gel matrix. Peptides were extracted from the gel pieces with ACN/0.1% formic acid and the samples were dried in a SpeedVac^TM^ vacuum concentrator. The dried peptides from each sample were reconstituted in 0.1% formic acid and quantified in a NanoDrop^TM^ (Thermo Scientific) to equalize all samples at an identical protein concentration before analysis on the liquid chromatography-tandem mass spectrometry (LC-MS/MS) system. Mass spectrometry (MS) analysis was performed using an Easy nLC 1200 UHPLC system coupled to a hybrid linear trap quadrupole Orbitrap Q-Exactive HF-X mass spectrometer (ThermoFisher Scientific). Software versions used for the data acquisition and operation were Tune 2.9 and Xcalibur 4.1.31.9. The acquired raw data were analyzed using Proteome Discoverer^TM^ 2.2 (Thermo Fisher Scientific). Normalization was performed based on specific abundance of human β-actin protein and samples were scaled to controls average. The MS proteomics data have been deposited to the ProteomeXchange Consortium via the PRIDE partner repository with the dataset identifier PXD014489 (Vizcaino et al., 2014).

### RNA extraction and gene expression analysis

Total RNA from control and DMF-treated HMECs was extracted using Tri Reagent^TM^ (Sigma-Aldrich), and RNA was purified with the Direct-zol™ RNA MiniPrep Kit (Zymo Research) according to the manufacturer’s instructions. RNA quality and amount was measured using a NanoDrop ND-1000 (Thermo Scientific). cDNA synthesis was performed using the PrimeScript™ RT reagent Kit (Takara) following purchaser’s instructions. mRNA expression analysis was determined using KAPA SYBR Fast Master Mix (2×) Universal (KAPA Biosystems) in an Eco Real-Time PCR System (Illumina). The following thermal cycling profile was used: 95 °C, 3 min; 40 cycles of 95 °C, 10 s and 54 °C, 30s. Primer sequences were as follows: *ACTB* forward: GACGACATGGAGAAAATCTG; *ACTB* reverse: ATGATCTGGGTCATCTTCTC; *SLC2A1* forward: ACCTCAAATTTCATTGTGGG; *SLC2A1* reverse: GAAGATGAAGAACAGAACCAG. qPCR data were normalized to *ATCB* expression.

### Western blot

Cells were lysed with RIPA (50 mM Tris-HCl pH 7.4, 150 nM NaCl, 1% Triton X-100, 0.25% sodium deoxycholate and 1 mM EDTA) or with 2x denaturing loading buffer. Samples were heated at 95 °C during 5 min and separated on 10% polyacrylamide gels. Proteins were transferred to nitrocellulose membranes and blocked with 10% (w/v) semiskimmed dried milk. Blocked membranes were incubated overnight with primary antibodies (anti-HIF-1α 1:500, anti-PHGDH 1:1000, anti-α-tubulin 1:10000, anti-calnexin 1:1000), washed and incubated with the peroxidase-linked secondary anti-rabbit or anti-mouse antibody for 1 h at room temperature. Membranes were washed and finally incubated with the Supersignal^®^ West Pico chemiluminescent substrate system (Thermo Scientific). Image captions were taken with the ChemiDoc^TM^ XRS+ System (Bio-Rad) using Image Lab software or either films were revealed using a Medical film processor from Konica Minolta (Tokyo, Japan). Densitometry analyses were made with Image J software.

### Metabolomics and labeling experiments using stable isotopes

Cells grown in 6-cm dishes to 80-90% confluence were washed and 10 mM glucose or [U-^13^C]-glucose and 4 mM glutamine or [U-^13^C]-glutamine were added for the indicated times in DMEM supplemented with 10% dFBS in the presence or absence of DMF. For experiments of serine and glycine withdrawal or labeling, 10 mM glucose, 4 mM glutamine, 0.4 mM serine or [U-^13^C]-serine and/or 0.4 mM glycine or [U-^13^C]-glycine were added for 24 h in serine and glycine free RPMI-1640 or DMEM supplemented with 10% dFBS containing or not DMF. For analysis of intracellular metabolites by gas chromatography/mass spectrometry (GC/MS), labeled cells were rinsed in ice-cold saline solution and lysed with three freeze-thaw cycles in cold 80% methanol. Debris was discarded by centrifugation and 50 nmol of sodium 2-oxobutyrate were added as internal standard to the supernatants. Samples were evaporated, derivatized with N-(Tert-Butyldimethylsilyl)-N-Methyltrifluoroacetamide (MTBSTFA) and analyzed on an Agilent 7890 gas chromatograph coupled to an Agilent 5975 mass selective detector as previously described (Faubert et al., 2017). Data were acquired using MSD ChemStation software (Agilent). For analysis of intracellular metabolites by liquid chromatography/mass spectrometry (LC/MS), samples obtained in the same way but without addition of sodium 2-oxobutyrate were evaporated. Dried samples were reconstituted in 0.1% formic acid in water and 5 µL were injected into a 1290 UHPLC liquid chromatography (LC) system interfaced to a high-resolution mass spectrometry (HRMS) 6550 iFunnel Q-TOF mass spectrometer (MS) (Agilent). The MS was operated in both positive and negative (ESI+ and ESI-) modes. Analytes were separated on an Acquity UPLC® HSS T3 column (1.8 μm, 2.1 x 150 mm, Waters, MA). The column was kept at room temperature. Mobile phase A composition was 0.1% formic acid in water and mobile phase B composition was 0.1% formic acid in 100% acetonitrile. The LC gradient was 0 min: 1% B; 5 min: 5% B; 15 min: 99%; 23 min: 99%; 24 min: 1%; 25 min: 1%. The flow rate was 250 μL/min. Data were acquired using Profinder B.08.00 SP3 software (Agilent). Intracellular relative abundance of metabolites was normalized to cell number for GC/MS samples or by total ion current (TIC) normalization for LC/MS samples and represented in a heatmap using Heatmapper (Babicki et al., 2016).

### Determination of PHGDH activity in HMECs

PHGDH activity was measured in control and DMF treated cells using a spectrophotometric assay as indicated by the supplier (Phosphoglycerate Dehydrogenase (PHGDH) Activity Assay Kit (Colorimetric), BioVision). 100 µg protein of cell lysates were added to each well along with the reaction mix and PHGDH activity was monitored at 450 nm at 37 °C using a FLUOstar Omega microplate reader from BMG LABTECH (Ortenberg, Germany). Additionally, cell lysates from the control condition were added along with DMSO or 100 µM DMF to the reaction mix.

### Statistical analysis

All results are expressed as means ± SD. Data shown for extracellular flux analysis are for an experiment with 3-5 replicates, which was repeated three times with similar results. Metabolite quantification using NOVA/ammonia kit was done with three replicates, and repeated several times. Metabolomics and labeling experiments were performed once with 3-4 replicates. Data in the remaining figure panels reflect at least 3 independent experiments. For proteomics analysis, abundance ratio p-values were calculated by ANOVA based on background population of peptides and proteins, and values of p<0.01 were considered to be statistically significant. For the rest of the experiments, statistical significance was determined using the two-sided Student t-test and values of p<0.05 were considered to be statistically significant. In all figures, the p-values were shown as: *p<0.05; **p<0.01; ***p<0.001; ****p<0.0001.

## Funding

M.O. was recipient of a predoctoral FPU grant from the Spanish Ministry of Education, Culture and Sport. M.B. was supported by Juan de la Cierva – Incorporation Program (IJC2018-037657-I), Spanish Ministry of Science and Innovation”. This work was supported by grants PID2019-105010RB-I00 (MICINN and FEDER), UMA18-FEDERJA-220 and PY20_00257 (Andalusian Government and FEDER) and funds from group BIO 267 (Andalusian Government). The “CIBER de Enfermedades Raras” and “CIBER de Enfermedades Cardiovasculares” are an initiative from the ISCIII (Spain). R.J.D. is funded by Howard Hughes Medical Institute and the National Cancer Institute (Grant R35CA22044901). The funders had no role in the study design, data collection and analysis, decision to publish or preparation of the manuscript.

## Competing interests

The authors declare no conflict of interest.

## Author Contributions

M.O., C.Y., B.M-P.; R.J.D., A.R.Q. and M.A.M. designed the research; M.O. and C.Y. performed the experiments and analyzed the data; H.S.V. performed LC/MS labeling and metabolomics assays; C.C. performed proteomics assays and analysis; M.O. wrote the original draft; M. B., B.M-P., R.J.D., A.R.Q. and M.A.M. reviewed and edited the final version of the manuscript; all authors reviewed the results and approved the final version of the manuscript.

## Supplementary Figures

**Supplementary Figure S1.**
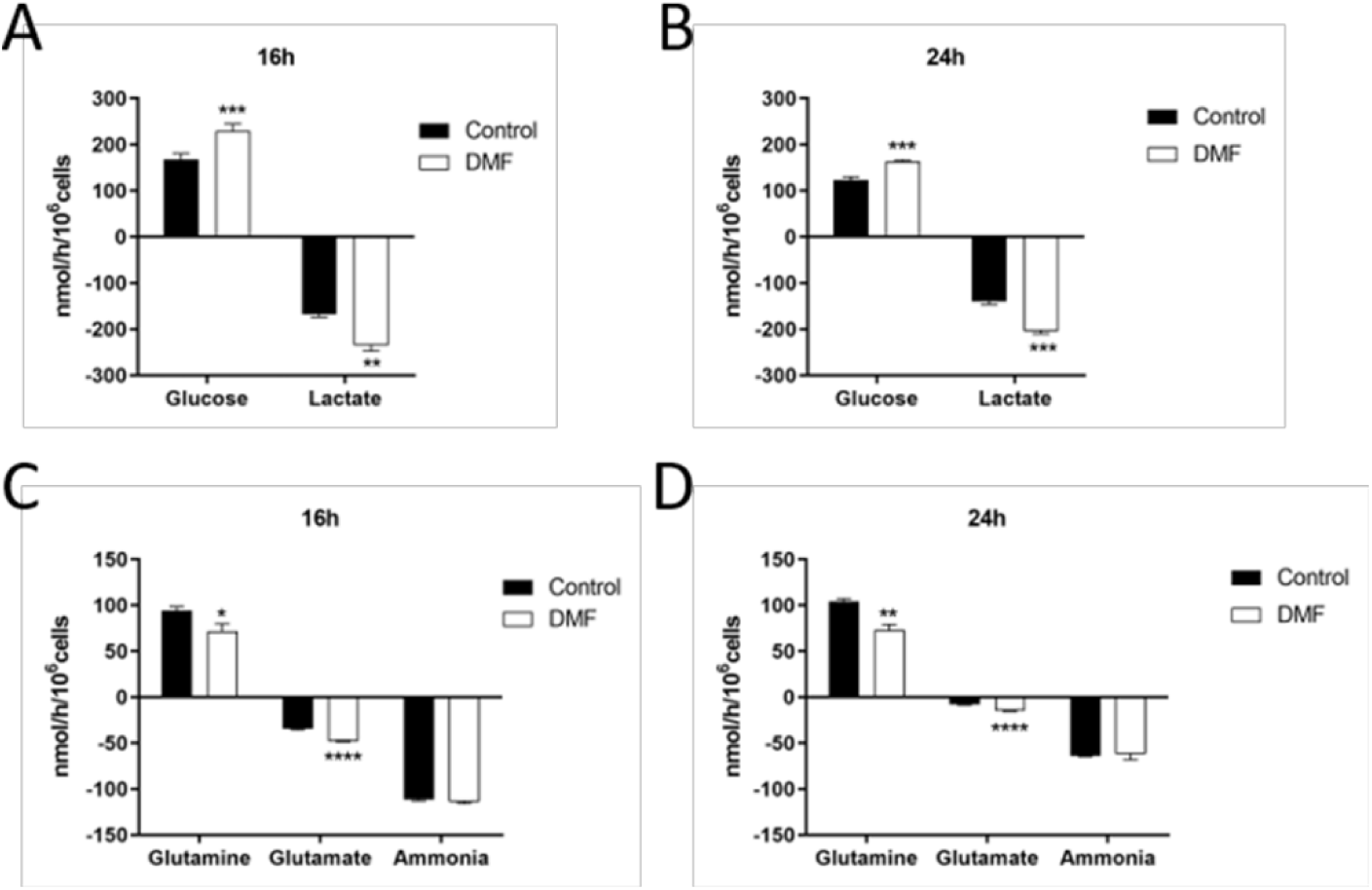
Effect of DMF on glucose uptake, lactate production, glutamine uptake and glutamate and ammonia production. (A) Glucose uptake and lactate secretion in cells treated with 100 µM DMF for 16 h or (B) 24 h in medium with 10 mM glucose and 4 mM glutamine. (C) Glutamine uptake and glutamate and ammonia secretion in cells treated with 100 µM DMF for 16 h or (D) 24 h in medium with 10 mM glucose and 4 mM glutamine. Data are expressed as means ± SD. *p<0.05, **p<0.01, ***p<0.001, ****p<0.0001 versus untreated control.

**Supplementary Figure S2.**
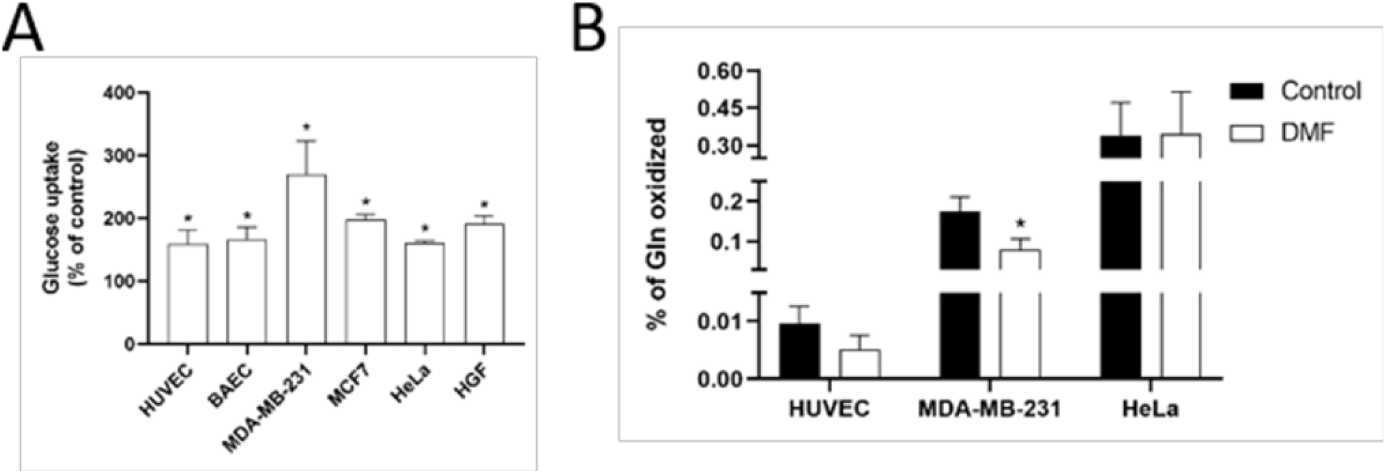
Effect of DMF on glucose uptake and glutamine oxidation in other endothelial cells, tumor cells and fibroblasts. (A) Glucose uptake and (B) glutamine oxidation after 30 minutes incubation with 5 mM glucose and 0.5 mM glutamine in different cell lines treated with 100 µM DMF for 16 h. Data are expressed as means ± SD. *p<0.05 versus untreated control.

**Supplementary Figure S3.**
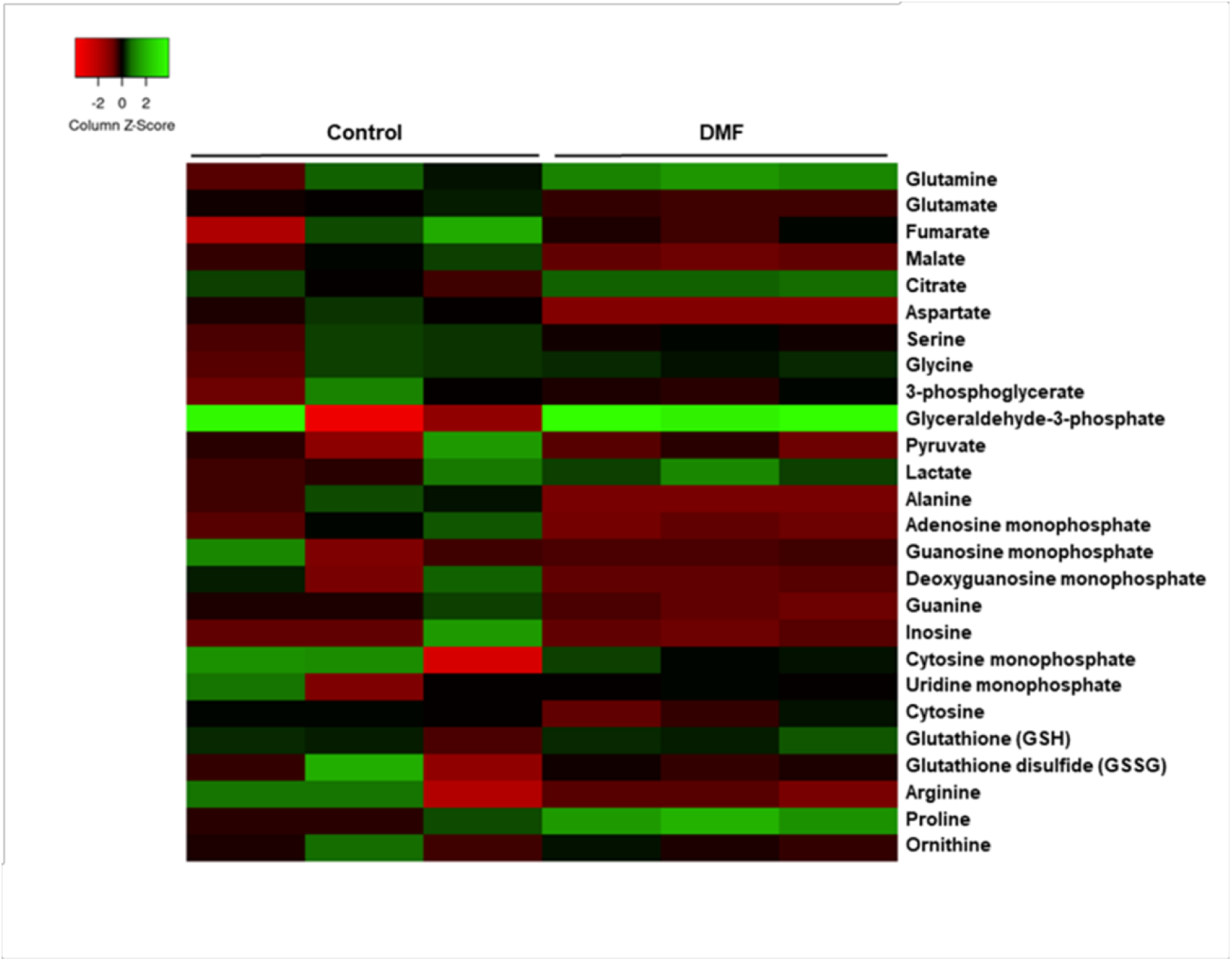
Heatmap of intracellular metabolite levels in HMECs treated or not with 100 µM DMF for 24 h. Data are expressed as fold from the control condition.

**Supplementary Figure S4.**
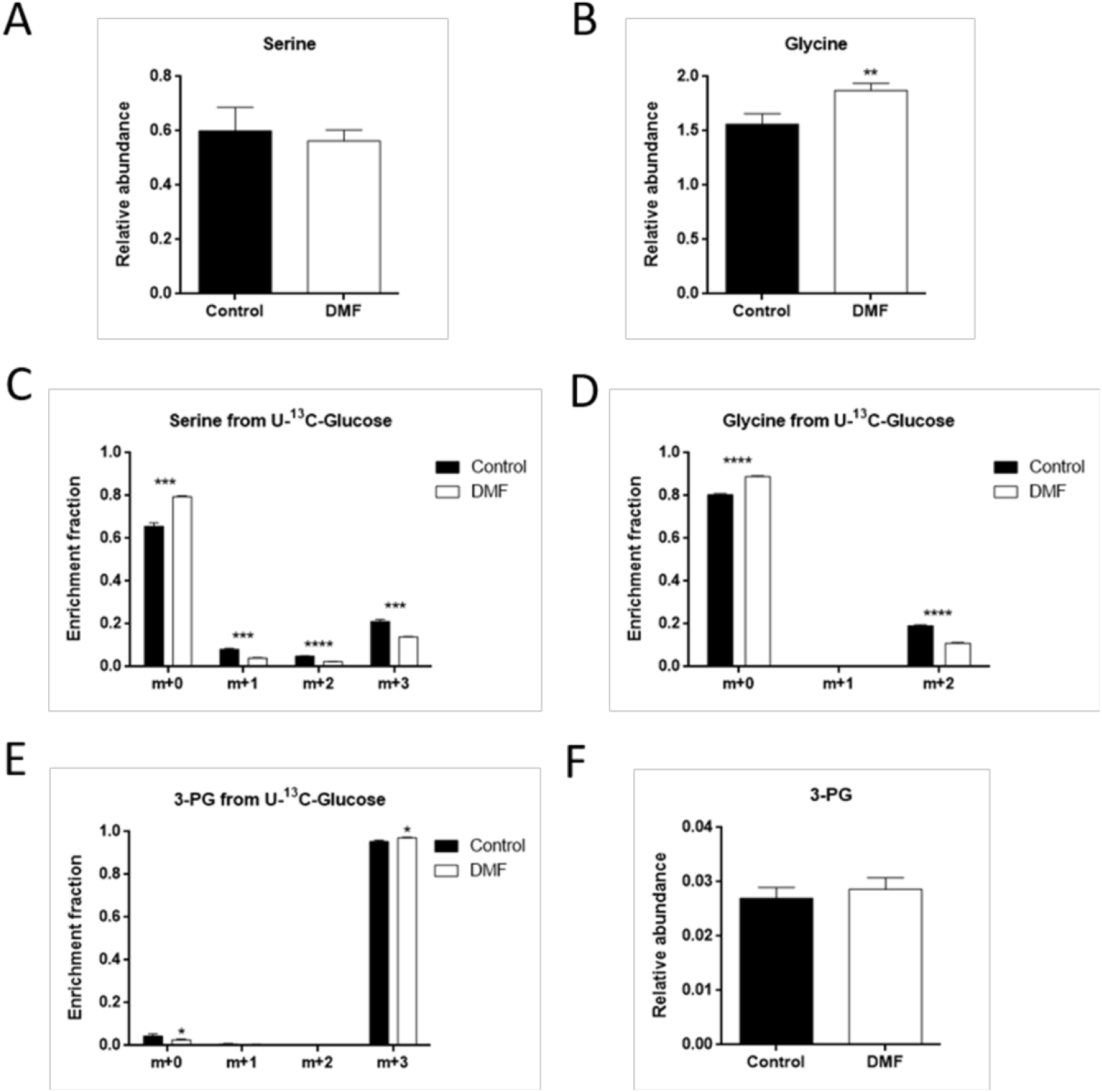
Effect of 100 µM DMF overnight incubation on serine and glycine synthesis pathway in HMECs. (A) Intracellular serine and (B) glycine levels in cells treated with 100 µM DMF for 16 h. (C) Fractional labeling of serine, (D) glycine and (E) 3-PG from [U-^13^C]-glucose and (F) intracellular 3-PG levels in cells treated with 100 µM DMF for 16 h. Data are expressed as means ± SD. **p<0.01, ***p<0.001, ****p<0.0001 versus untreated control.

**Supplementary Figure S5.**
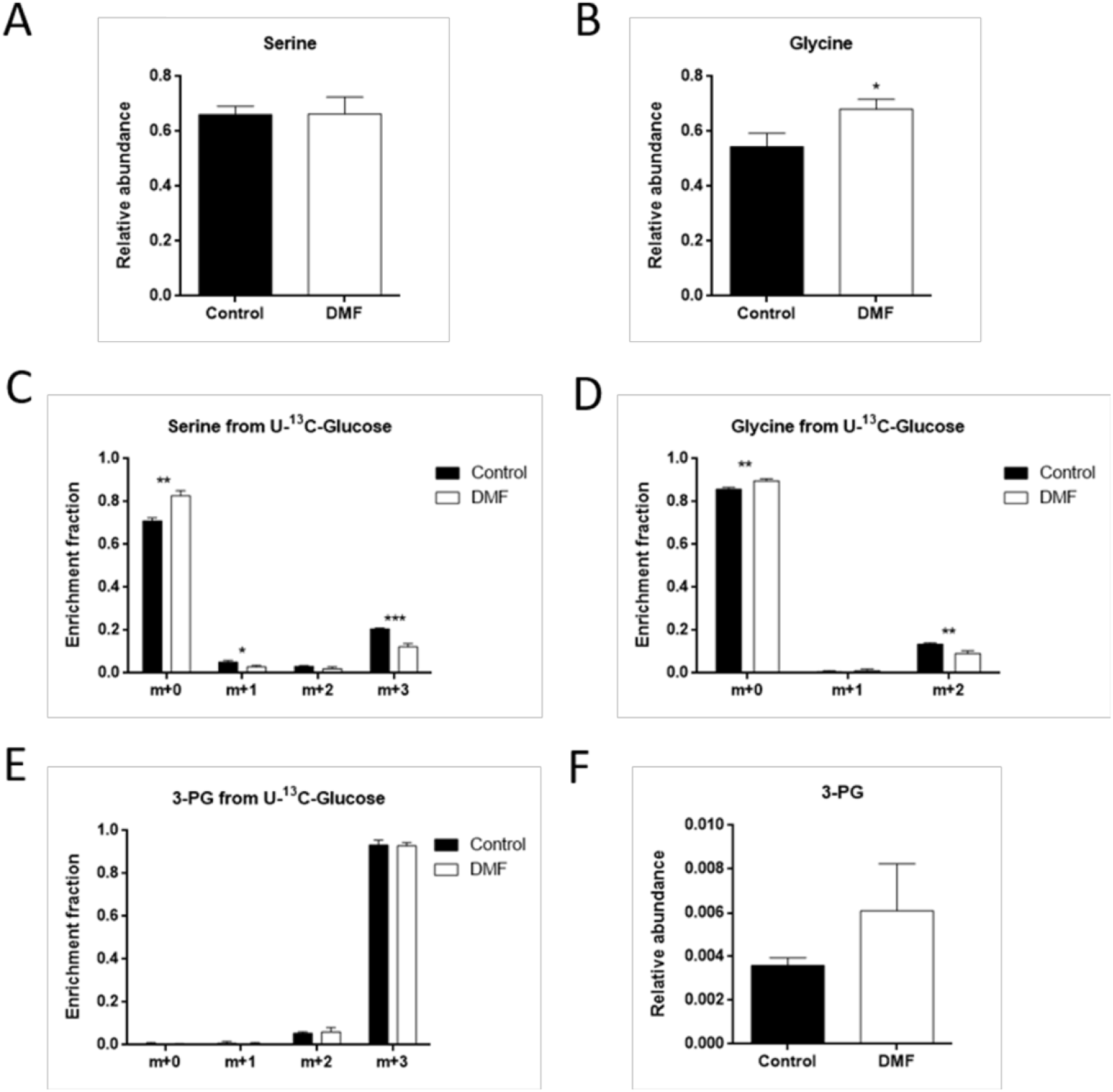
Effect of 50 µM DMF for 24 h on serine and glycine synthesis pathway in HMECs. (A) Intracellular serine and (B) glycine levels in cells treated with 50 µM DMF for 24 h. (C) Fractional labeling of serine, (D) glycine and (E) 3-PG from [U-^13^C]-glucose and (F) intracellular 3-PG levels in cells treated with 50 µM DMF for 24 h. Data are expressed as means ± SD. *p<0.05, **p<0.01, ***p<0.001 versus untreated control.

**Supplementary Figure S6.**
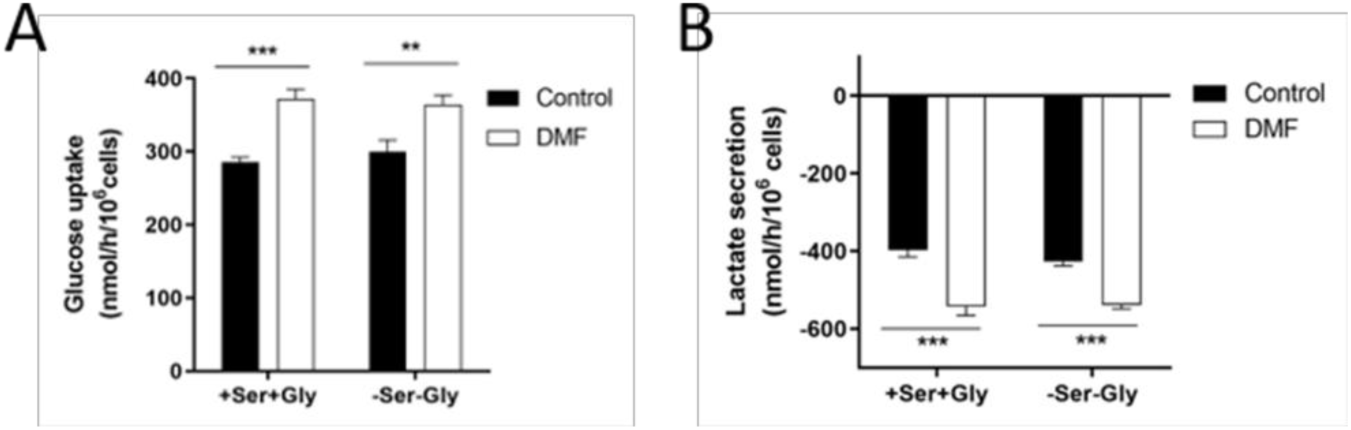
Effect of DMF on glucose uptake and lactate secretion in the absence of extracellular serine and glycine. (A) Glucose uptake and (B) lactate secretion in HMECs treated with 100 µM DMF for 24 h in medium with 10 mM glucose and 4 mM glutamine in the presence or absence of 0.4 mM extracellular serine and glycine. Data are expressed as means ± SD. **p<0.01, ***p<0.001 versus untreated control.

## Supplementary Tables

**Supplementary Table S1.**
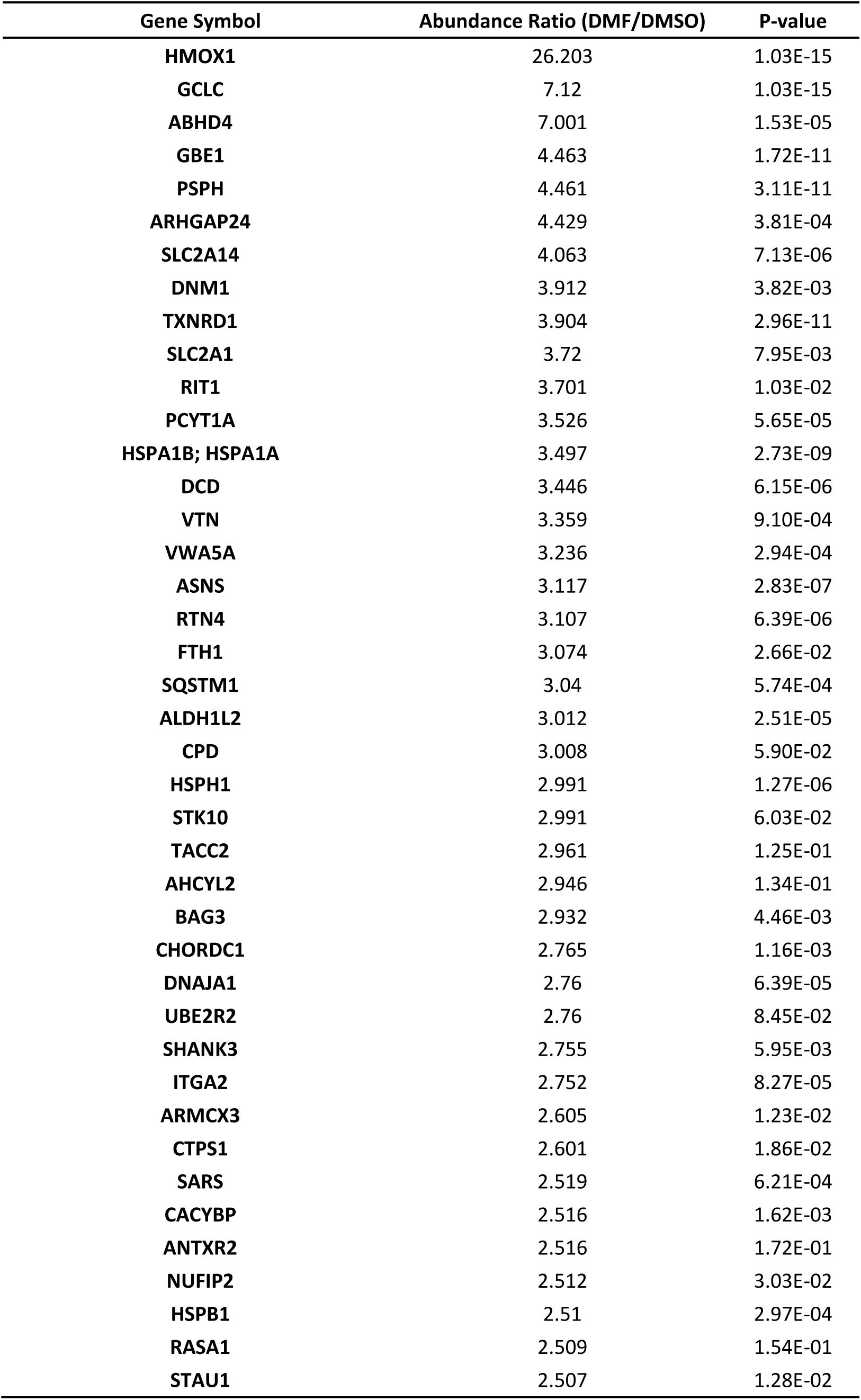
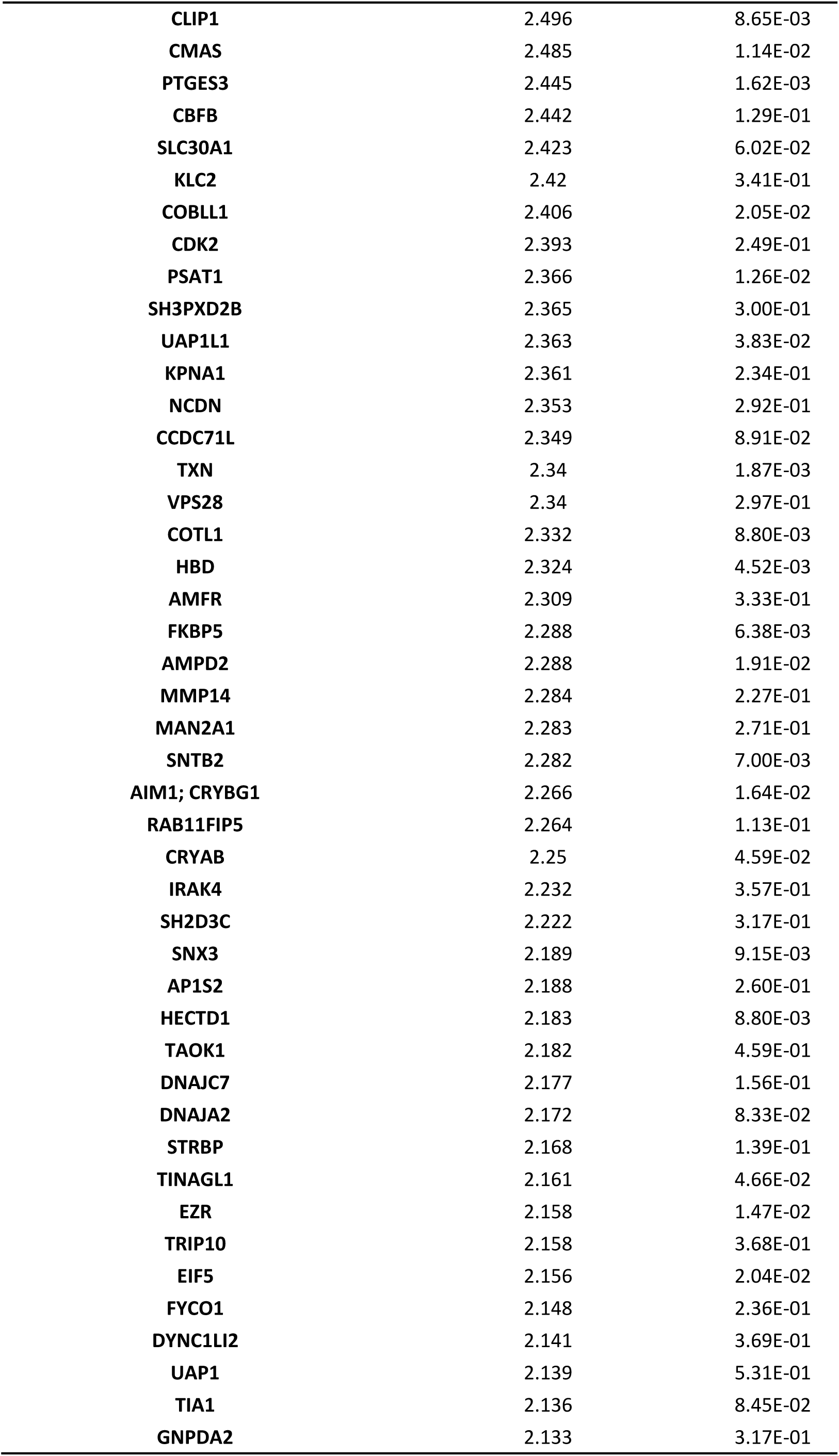
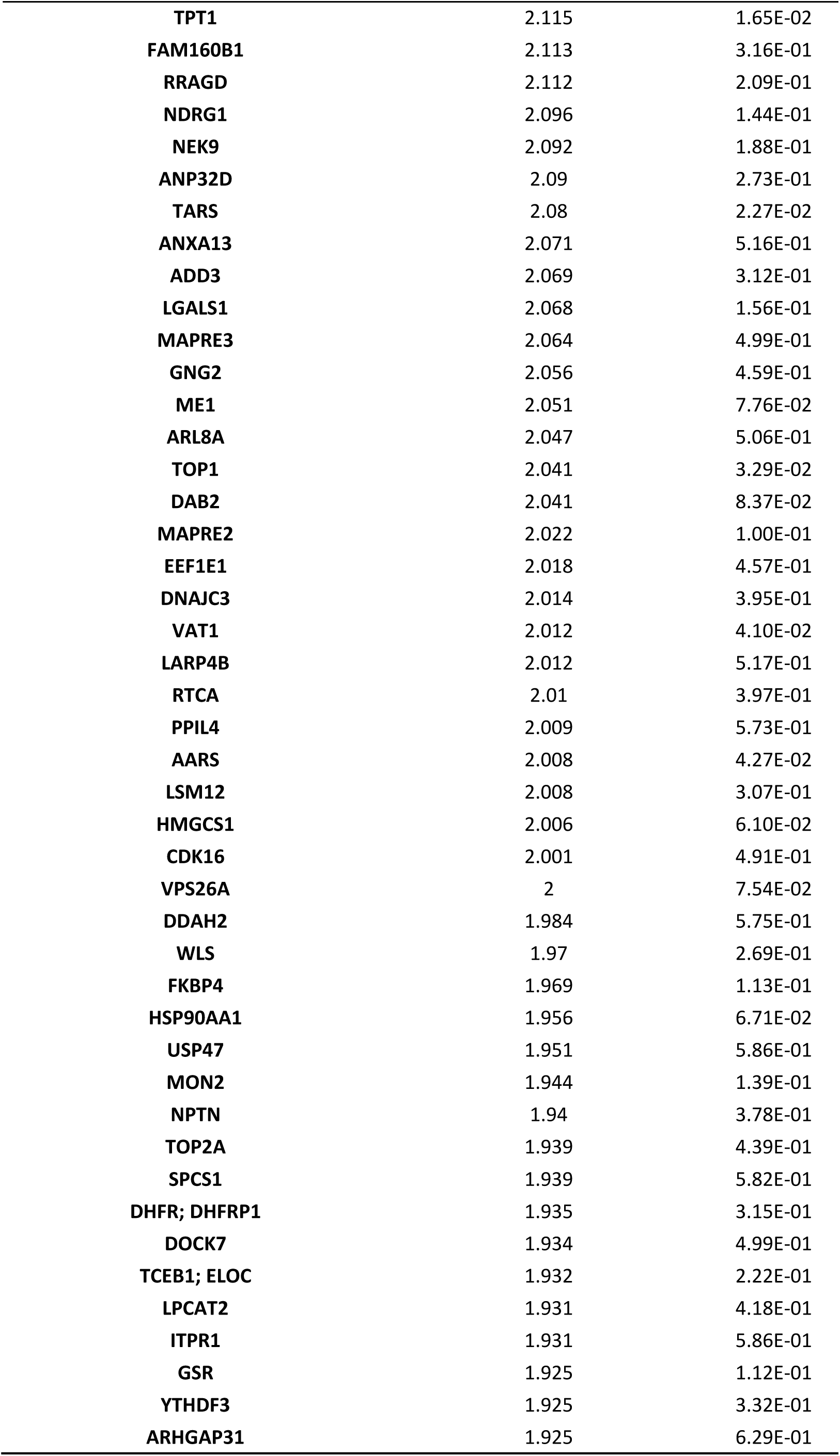
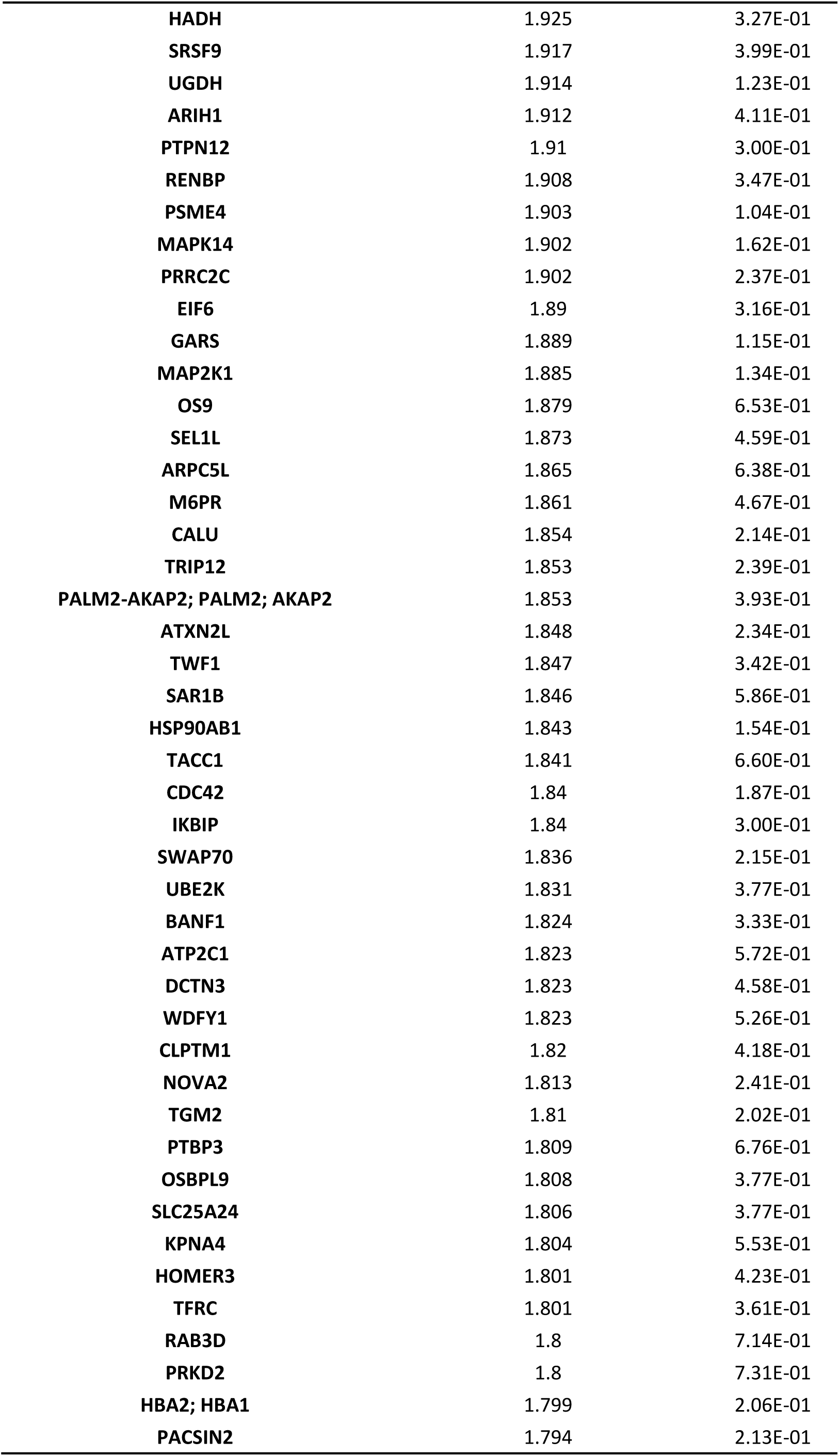
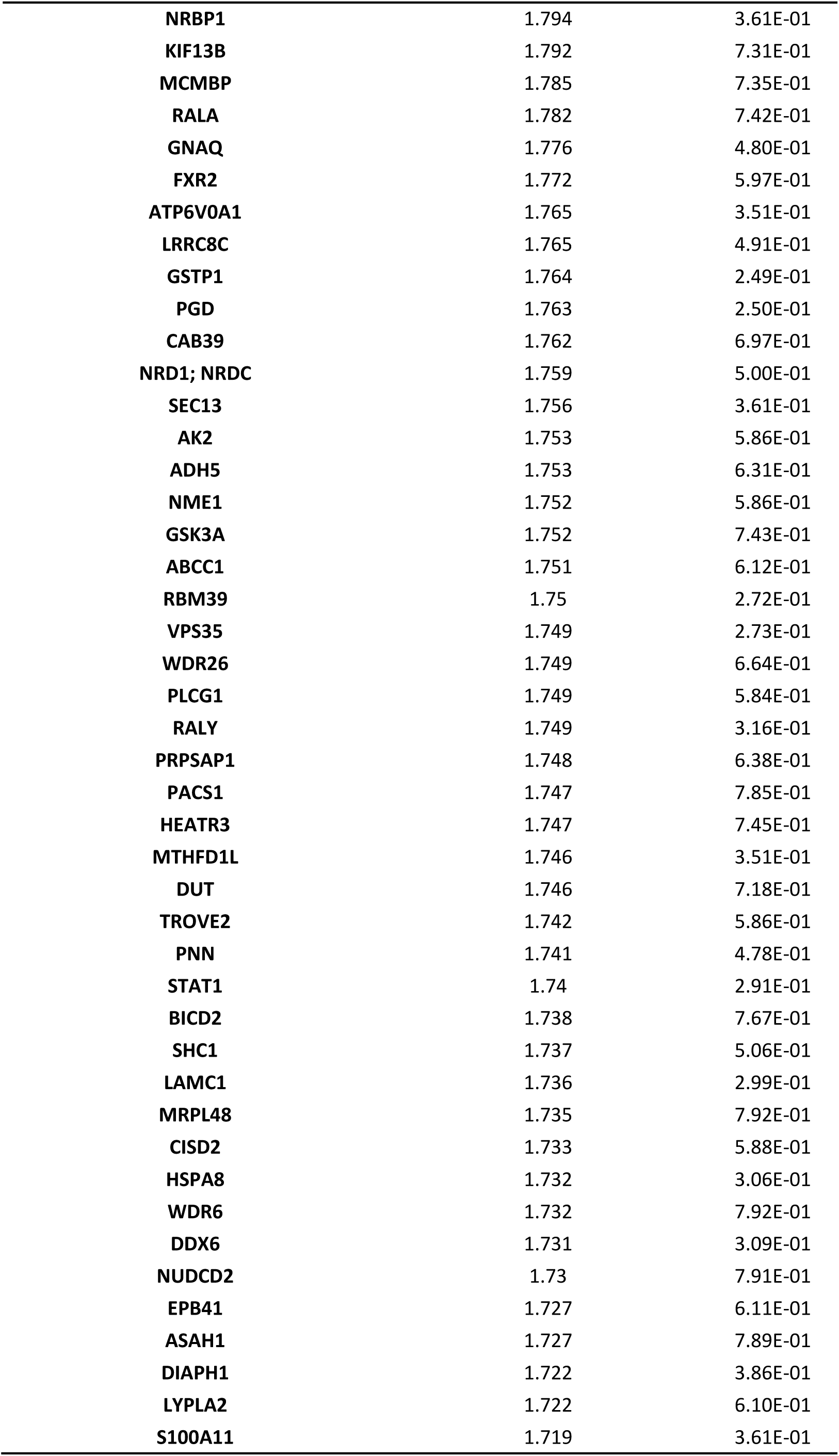
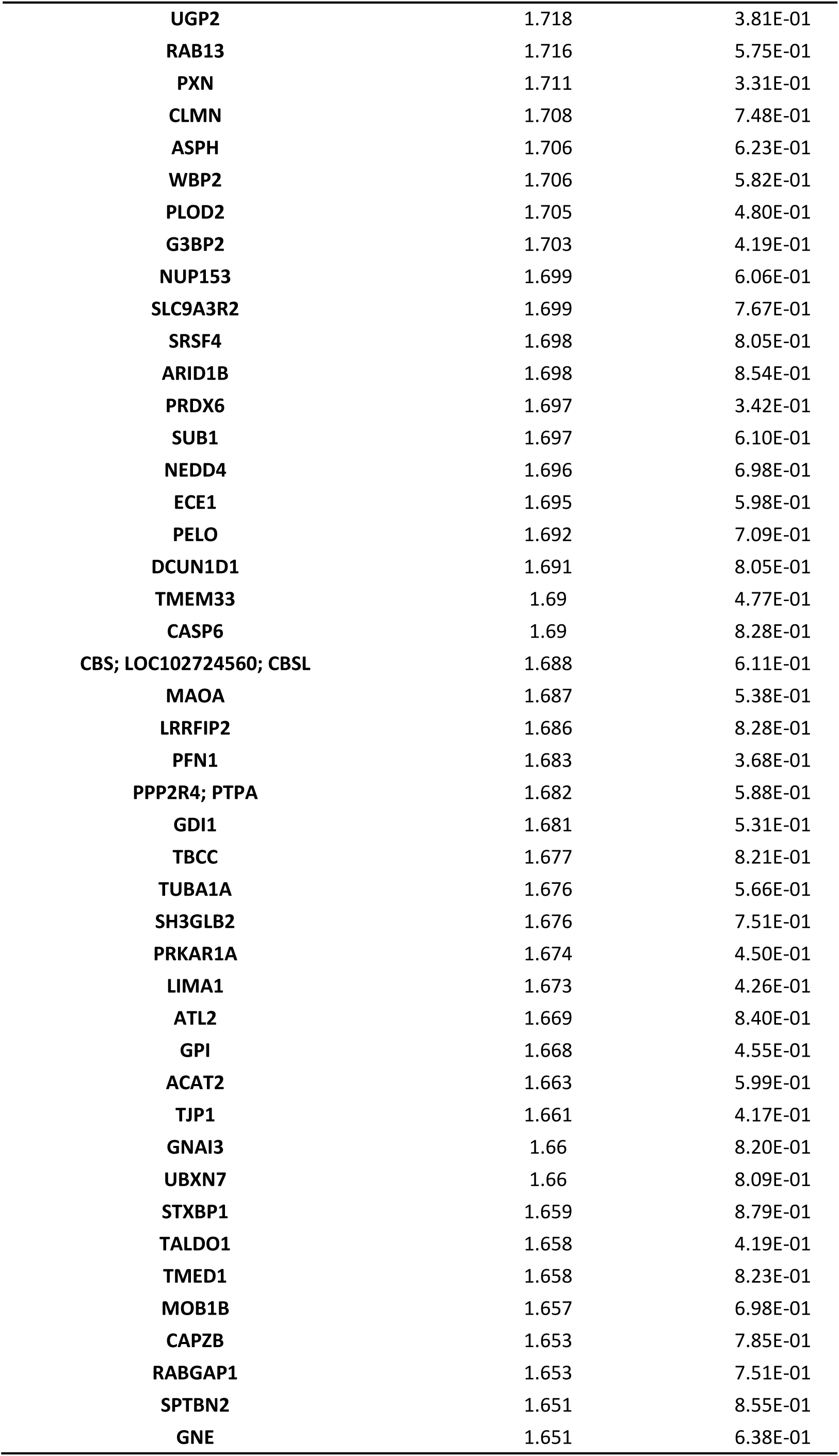
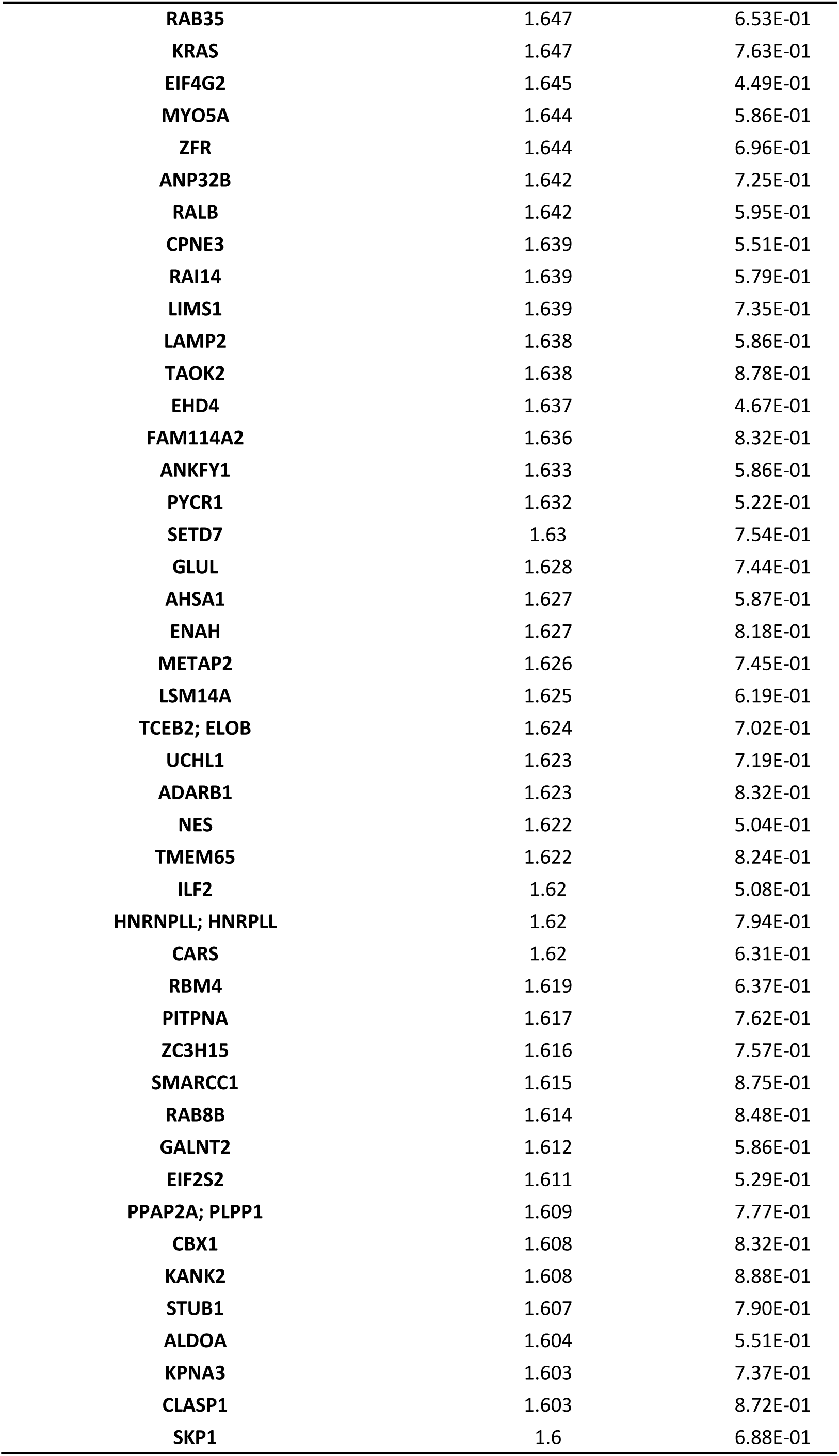
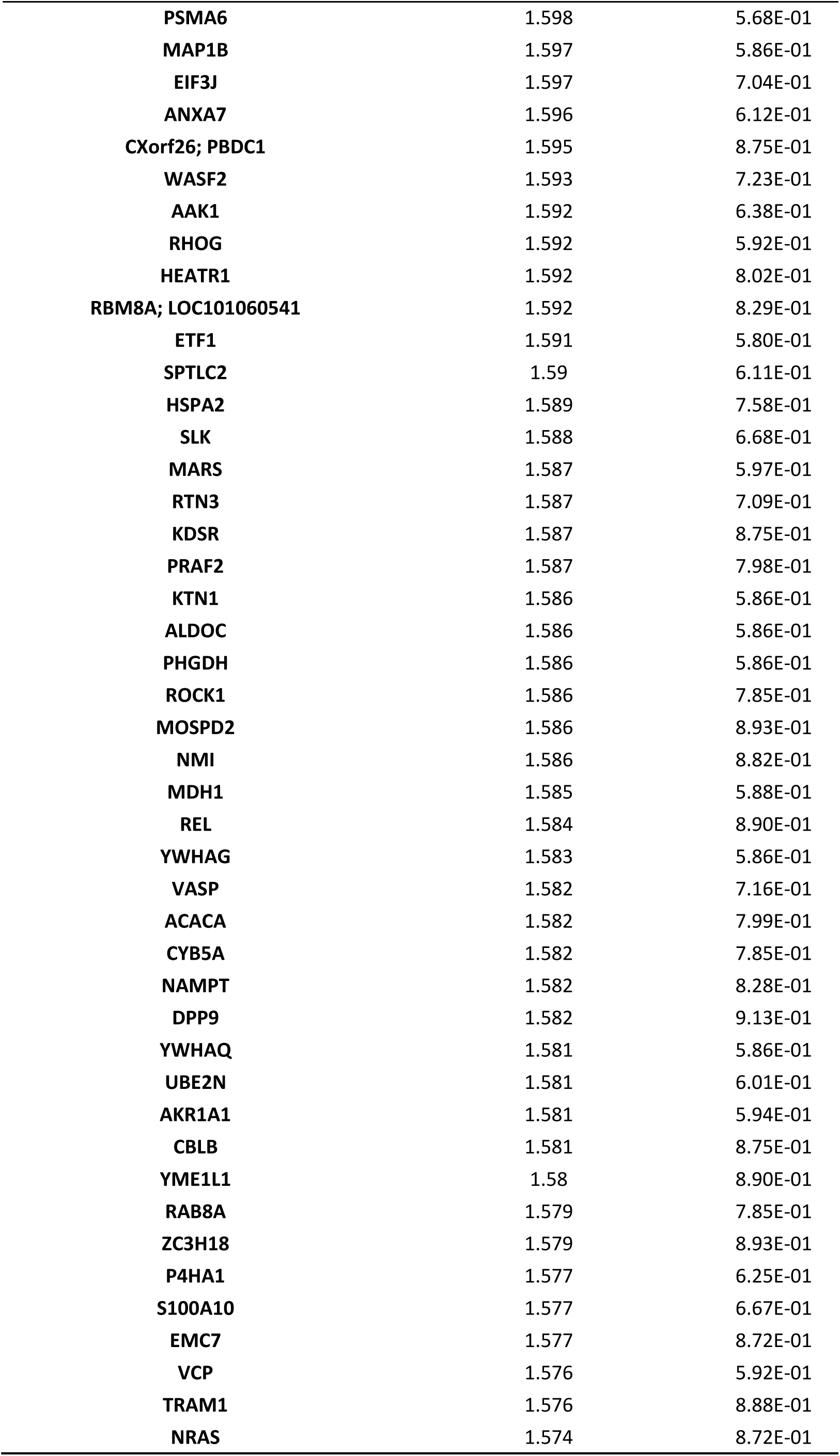
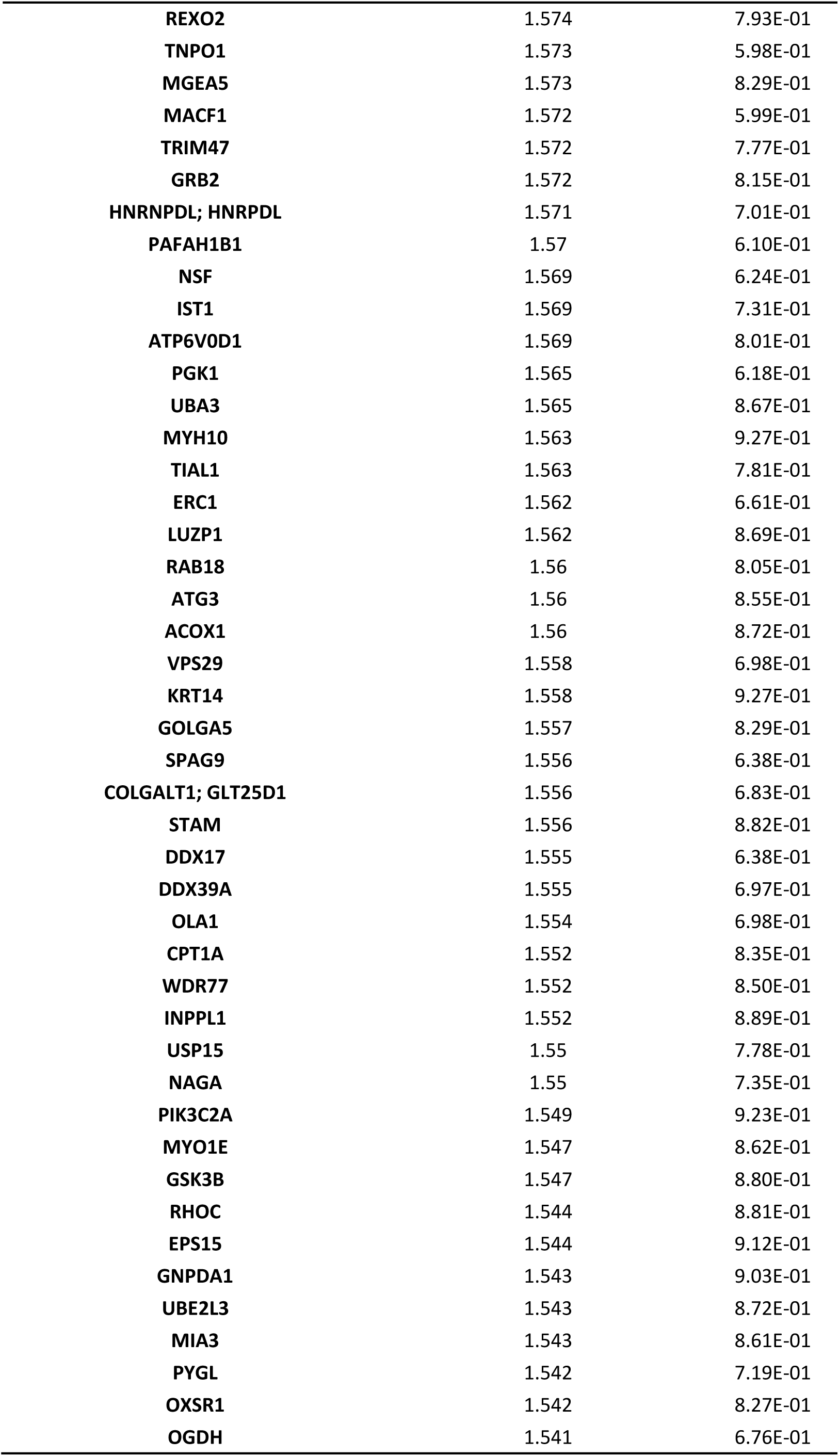
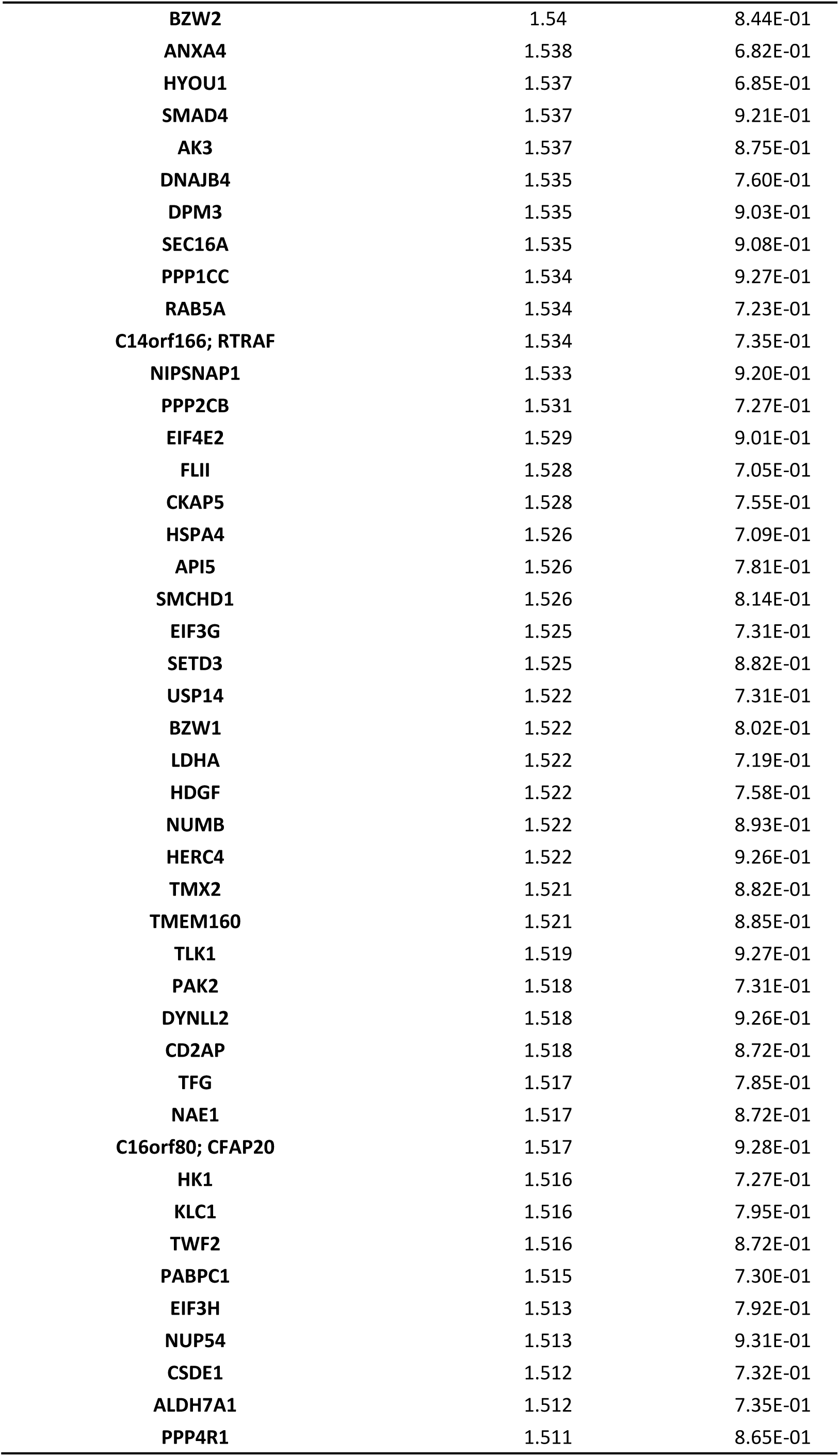
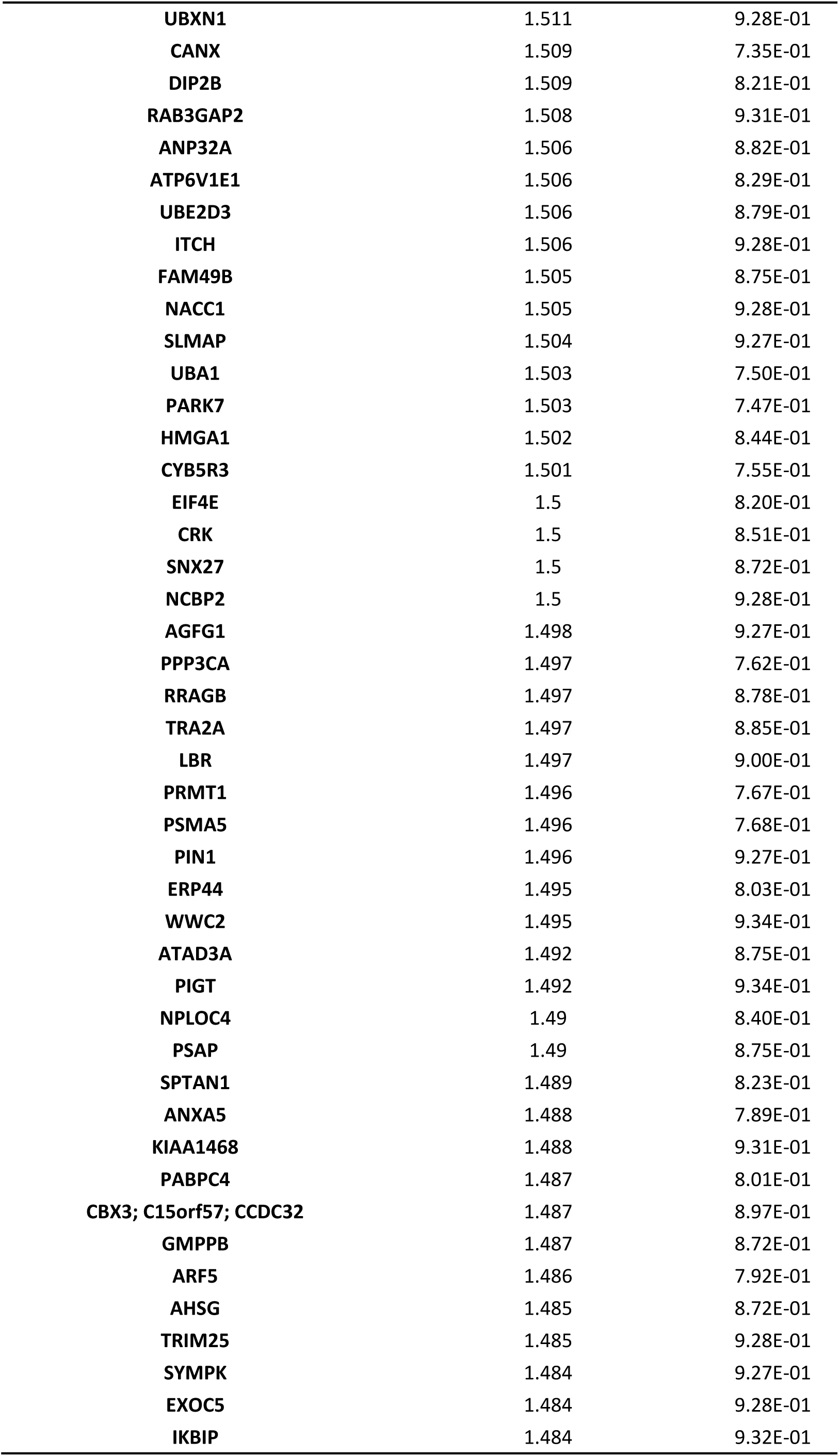
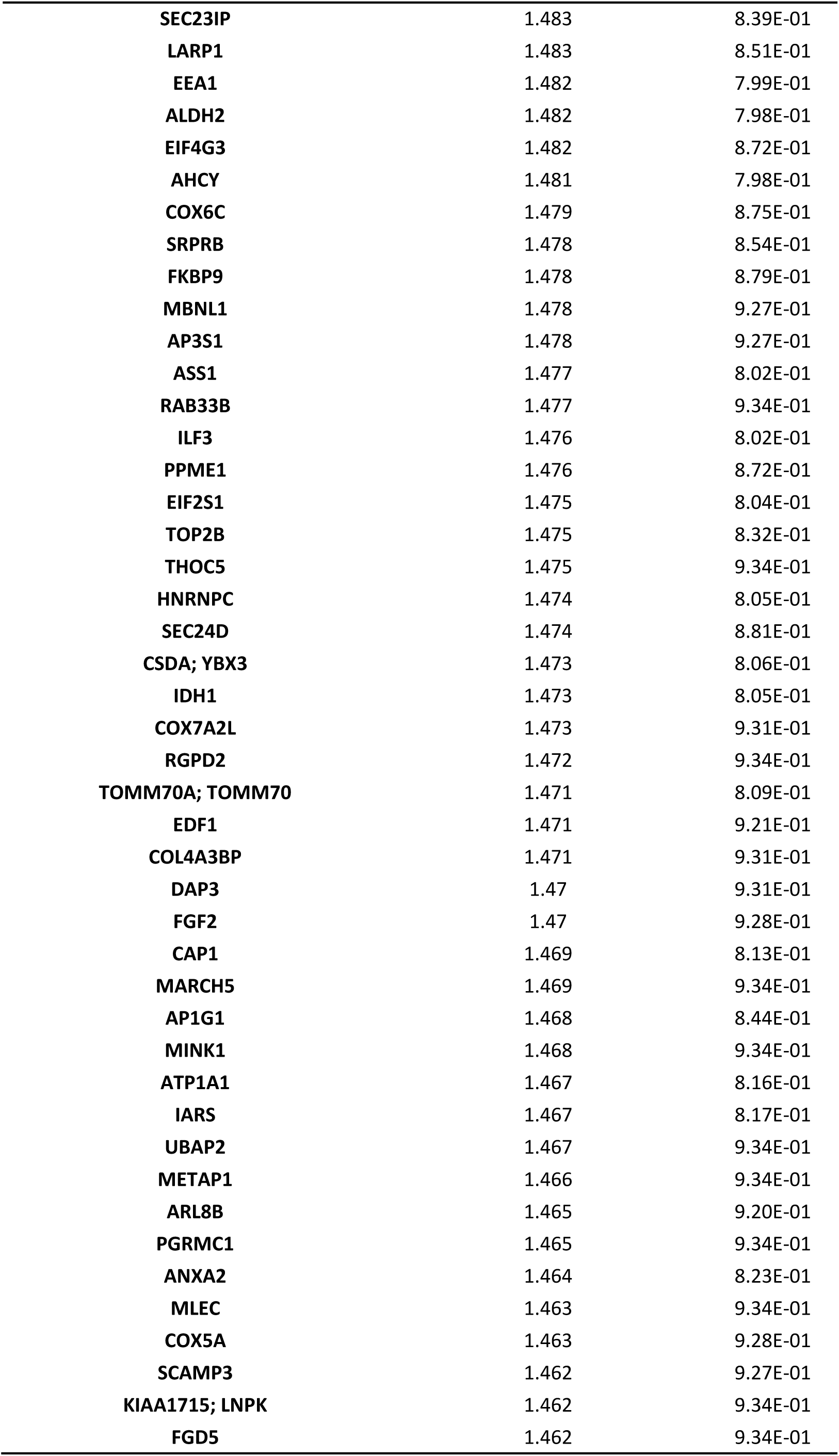
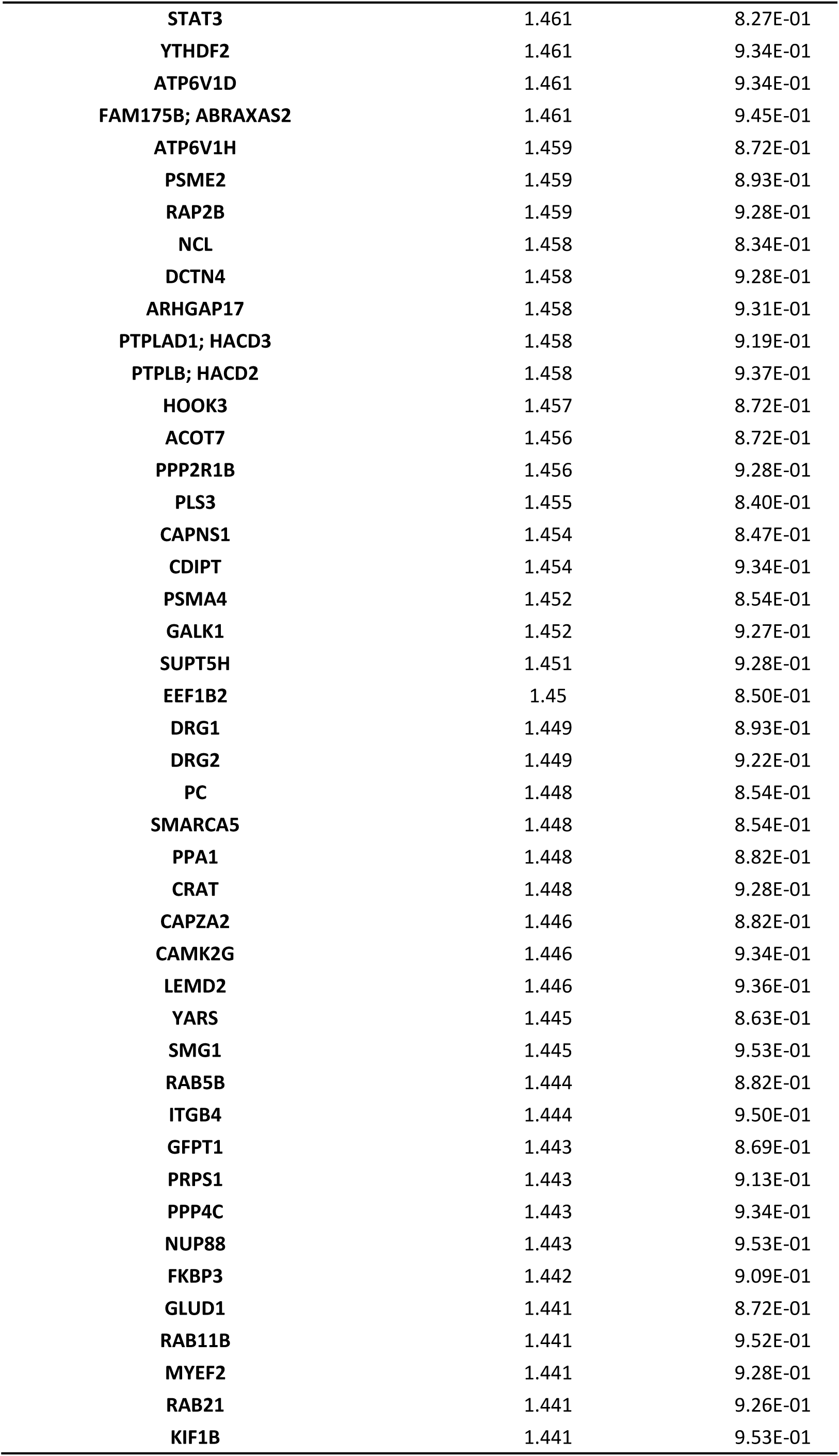
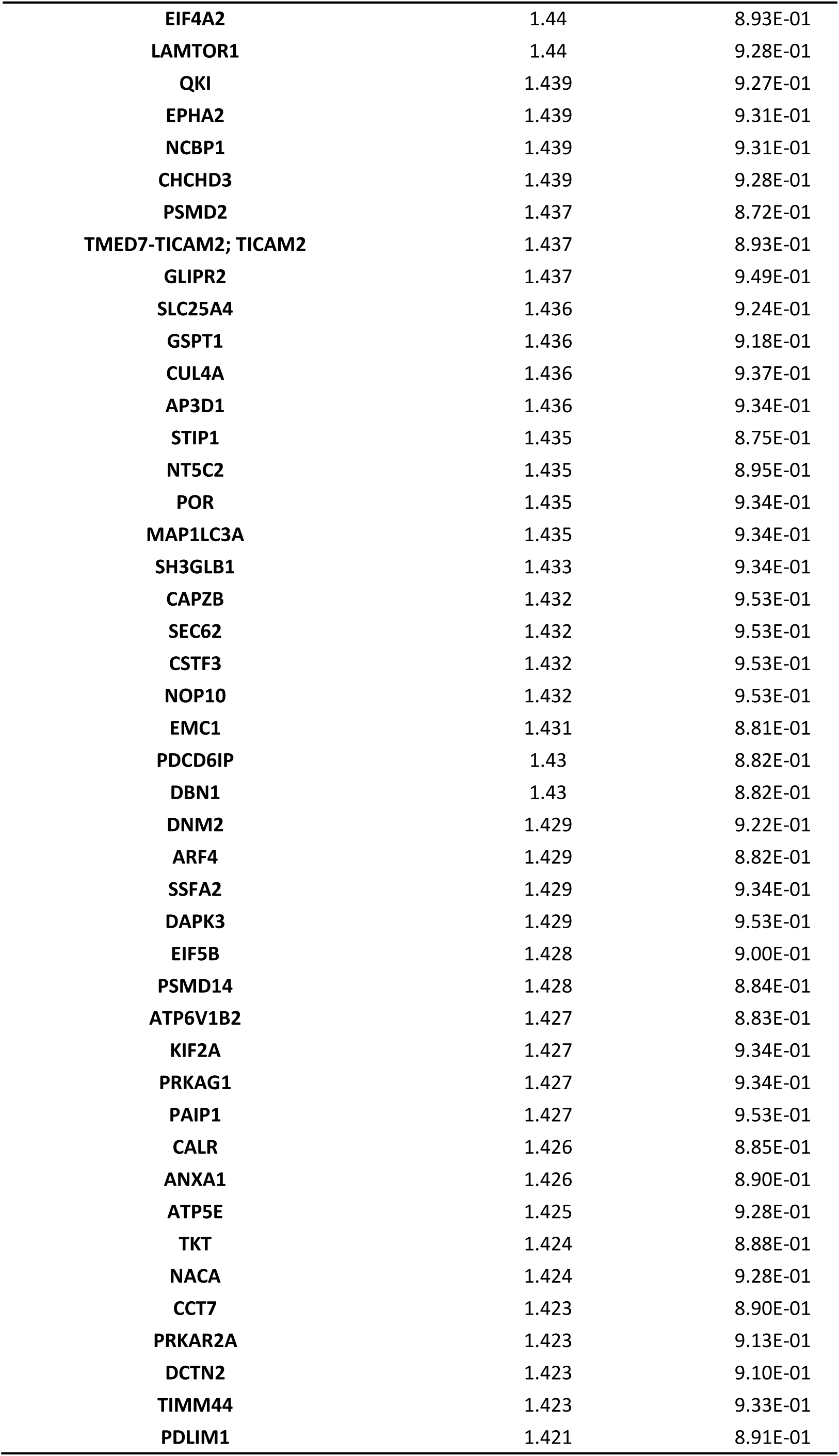
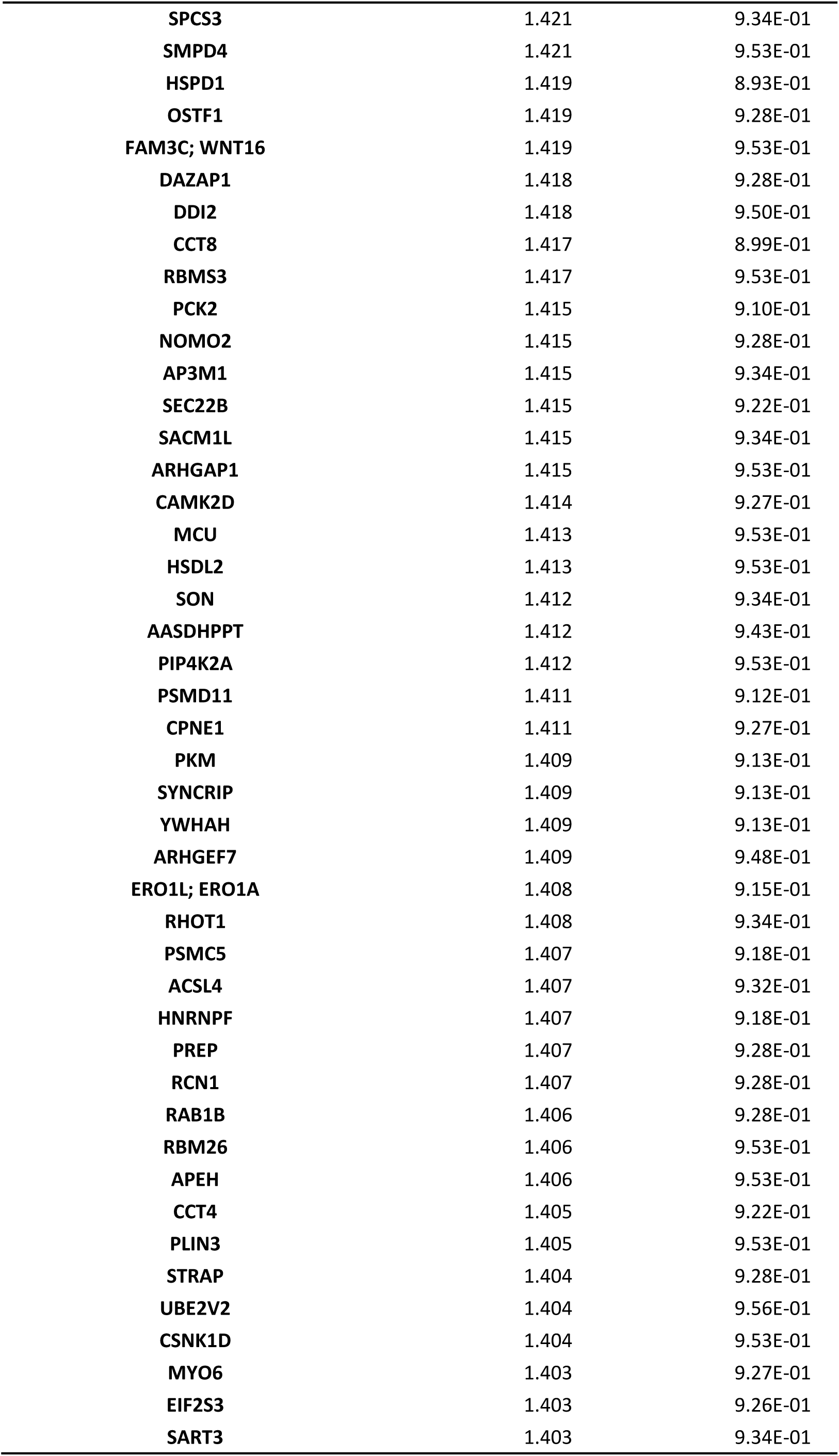
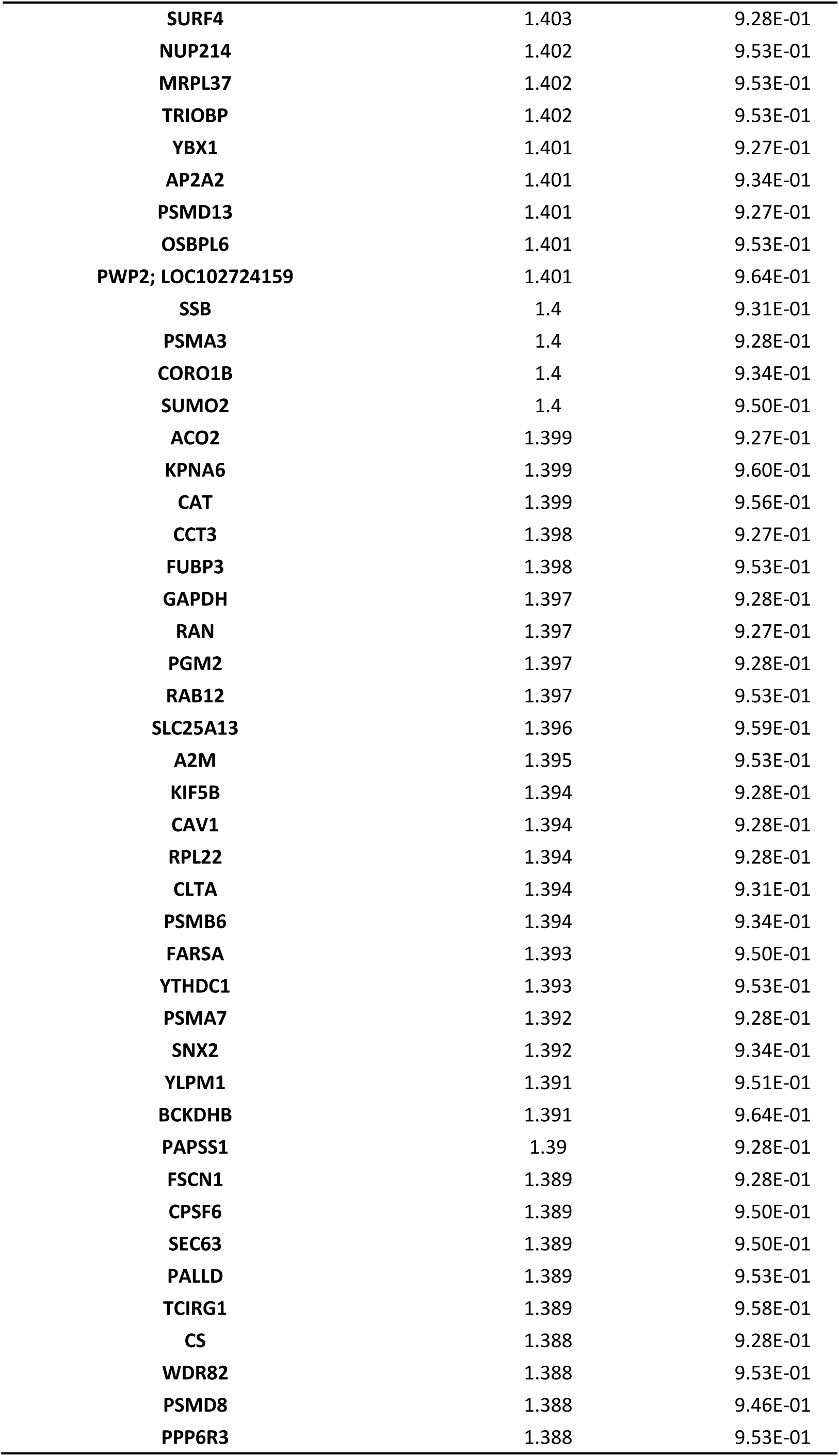
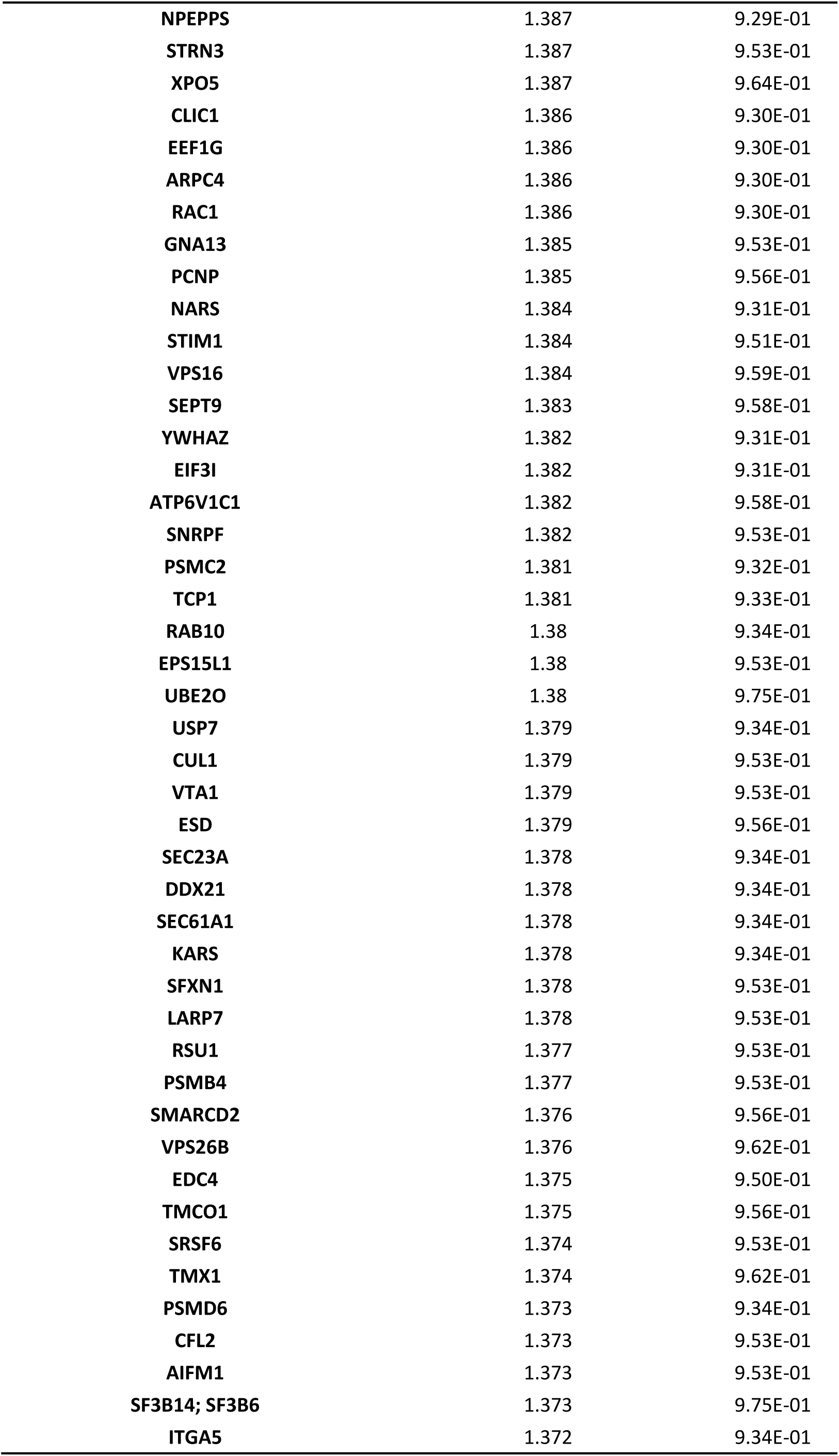
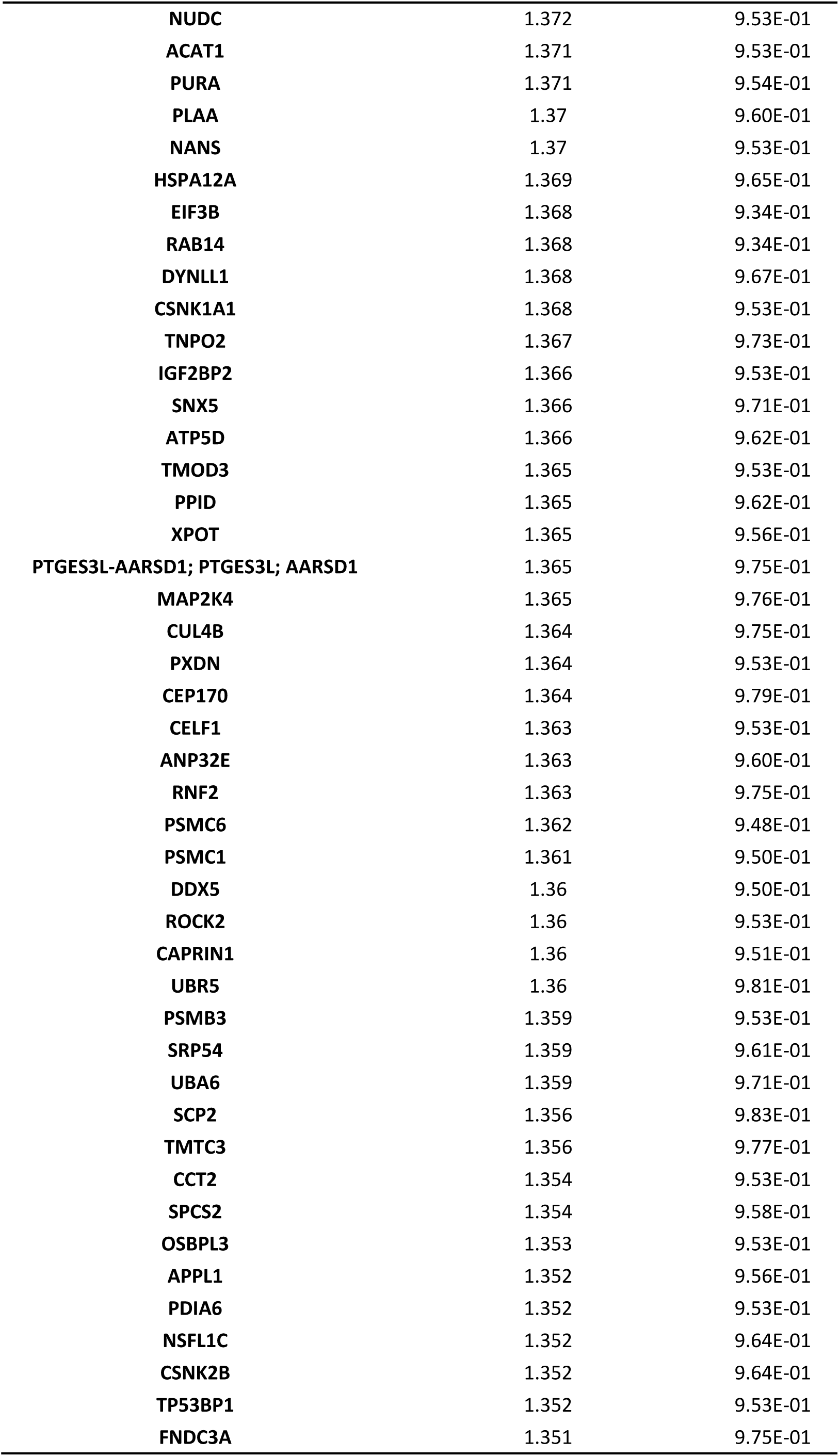
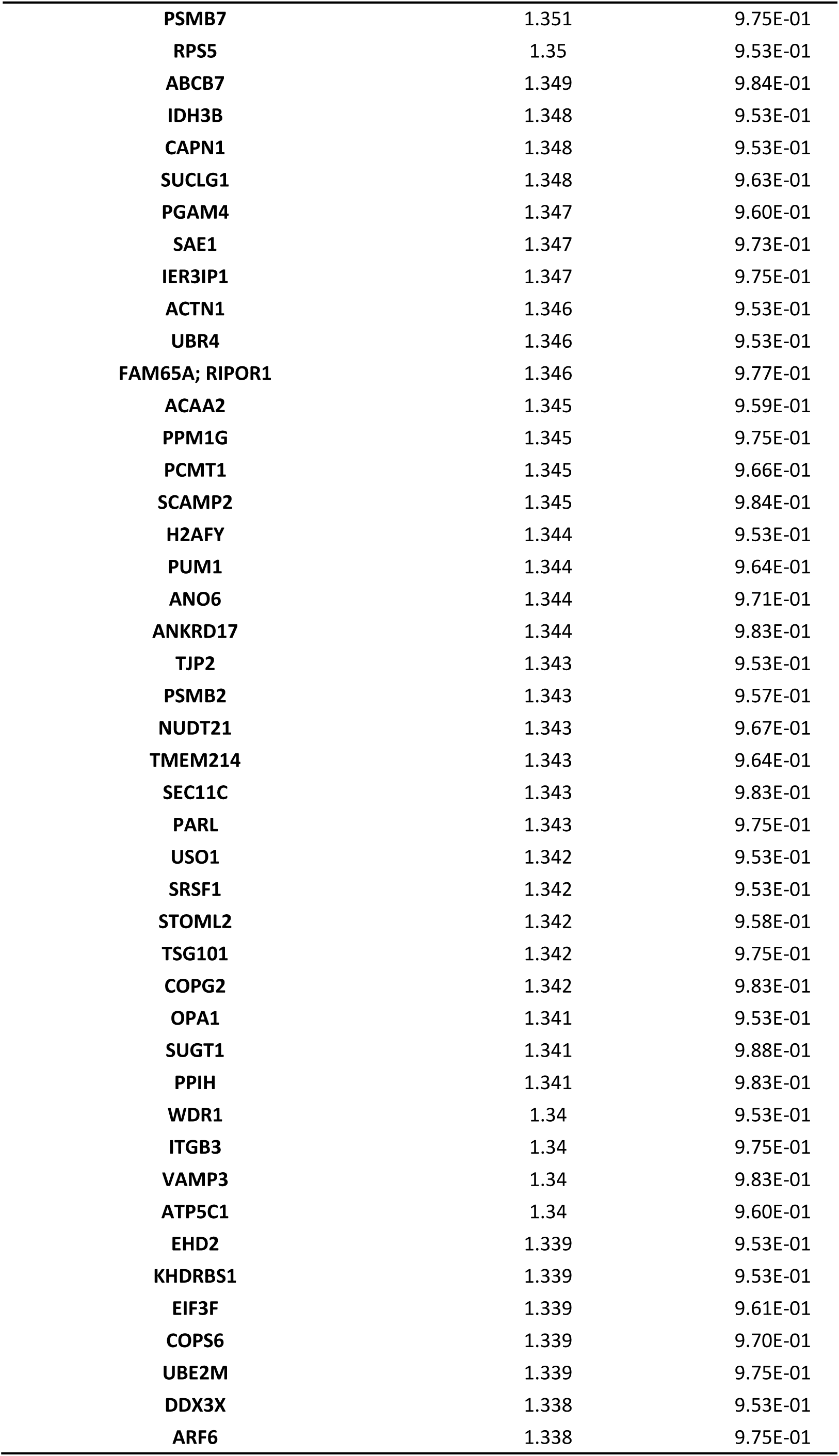
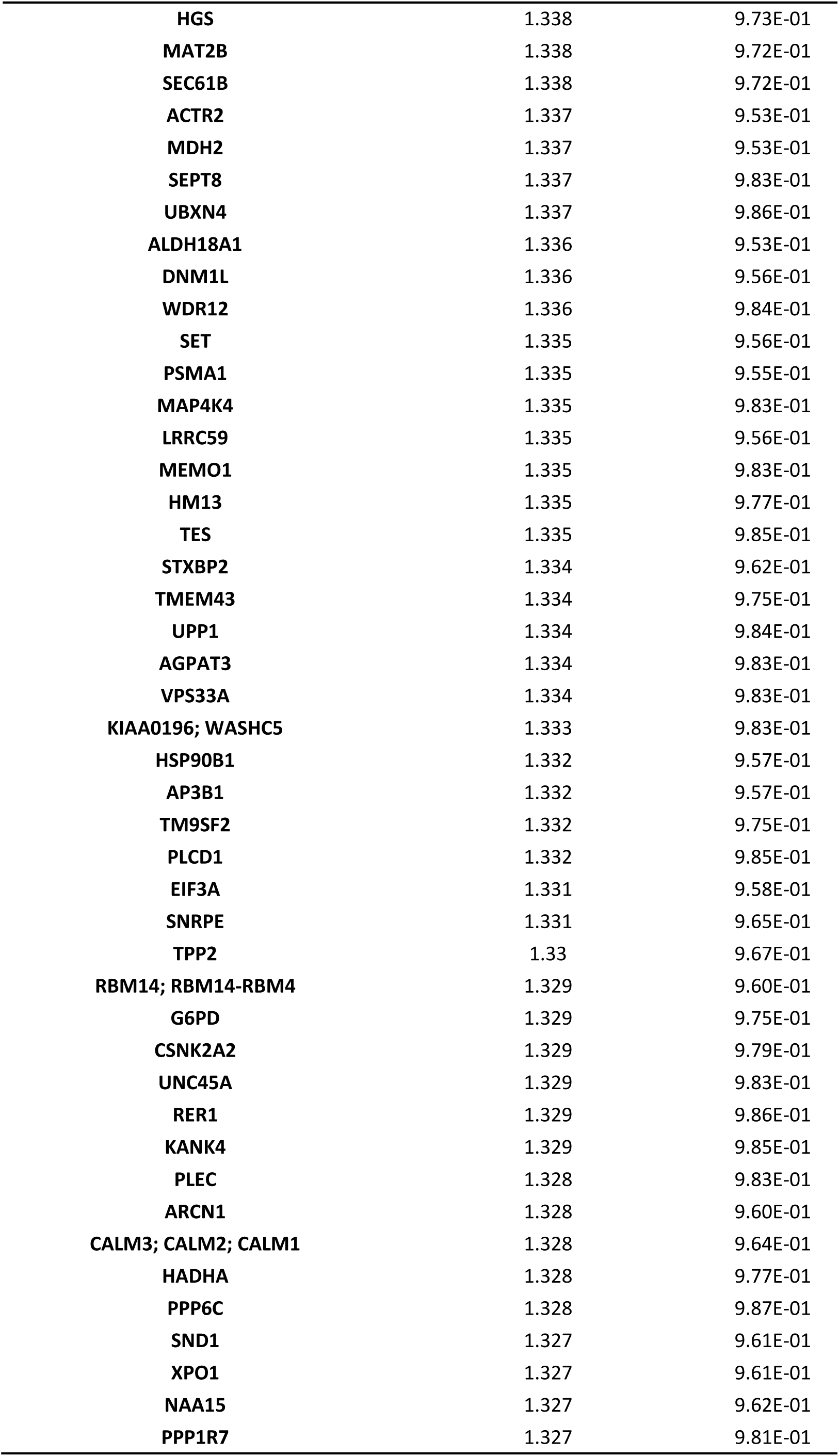
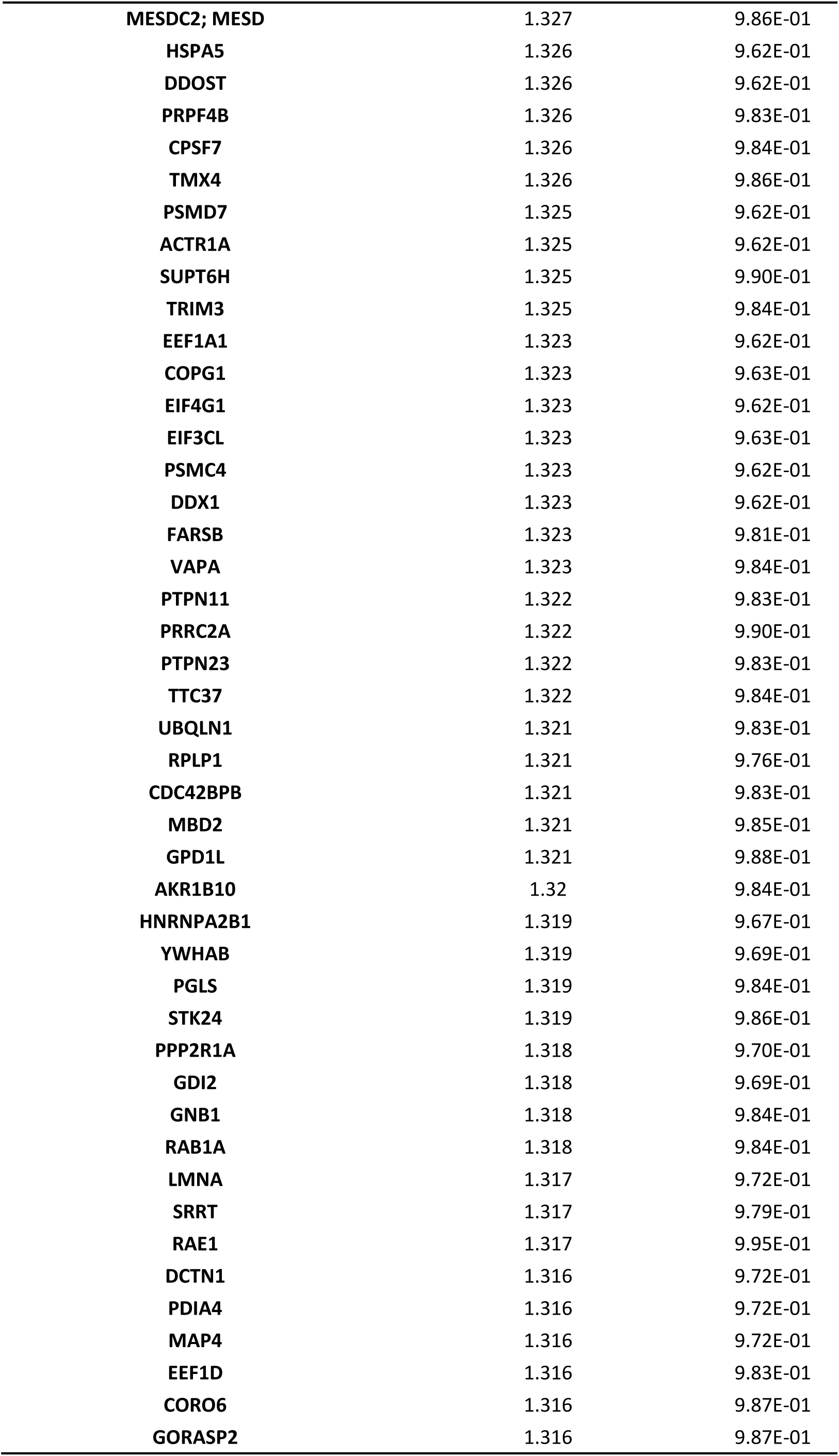
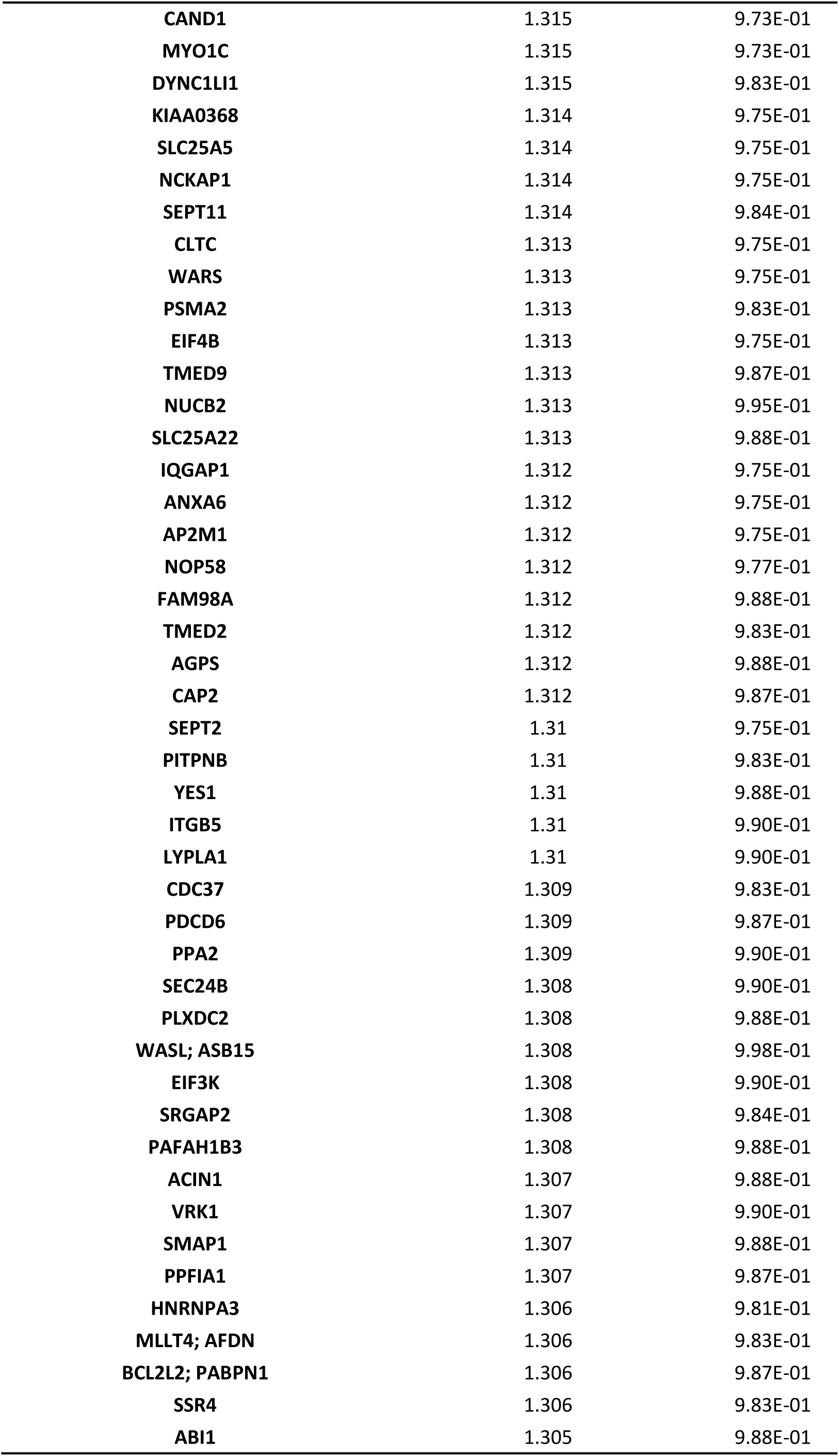
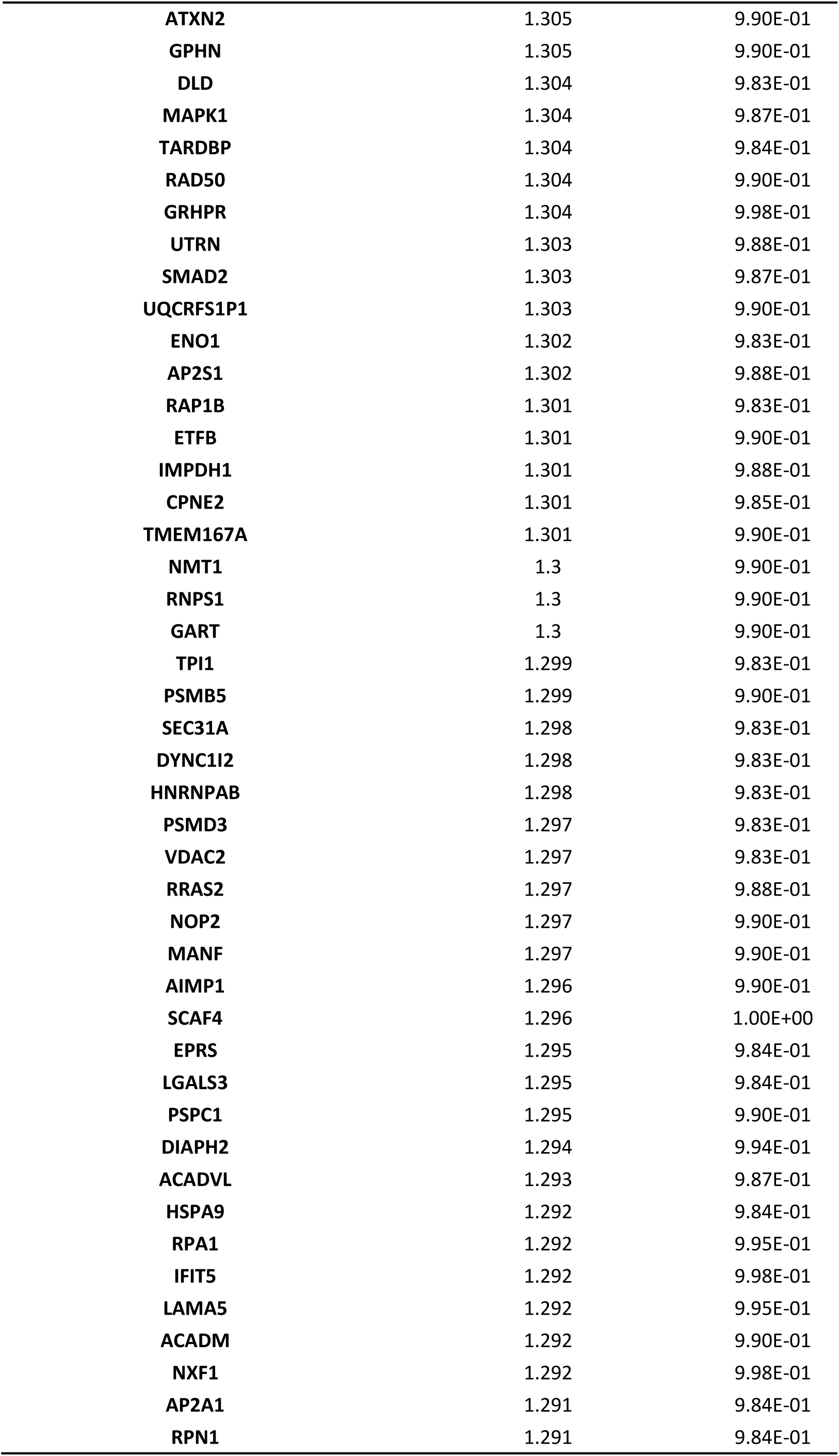
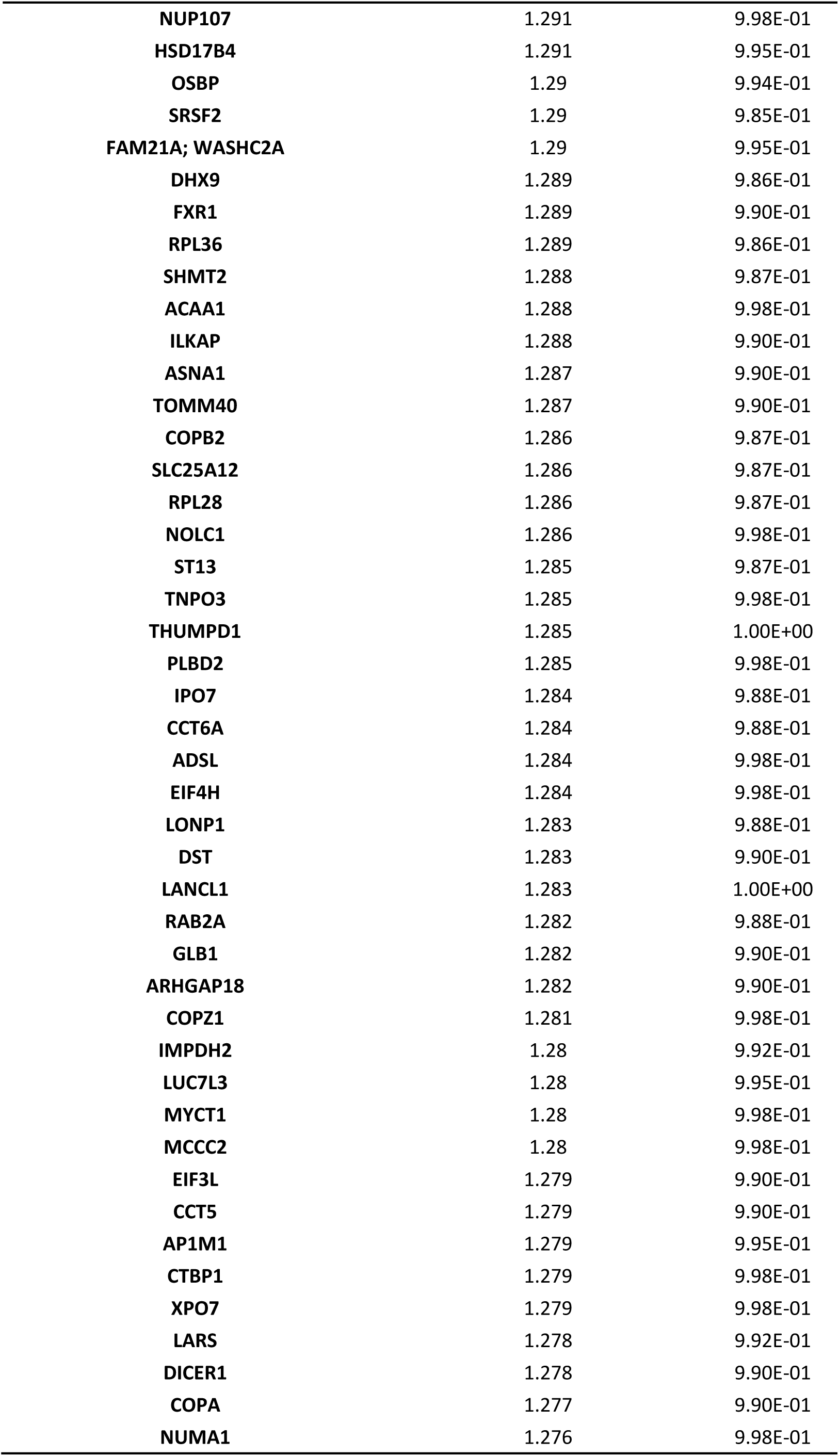
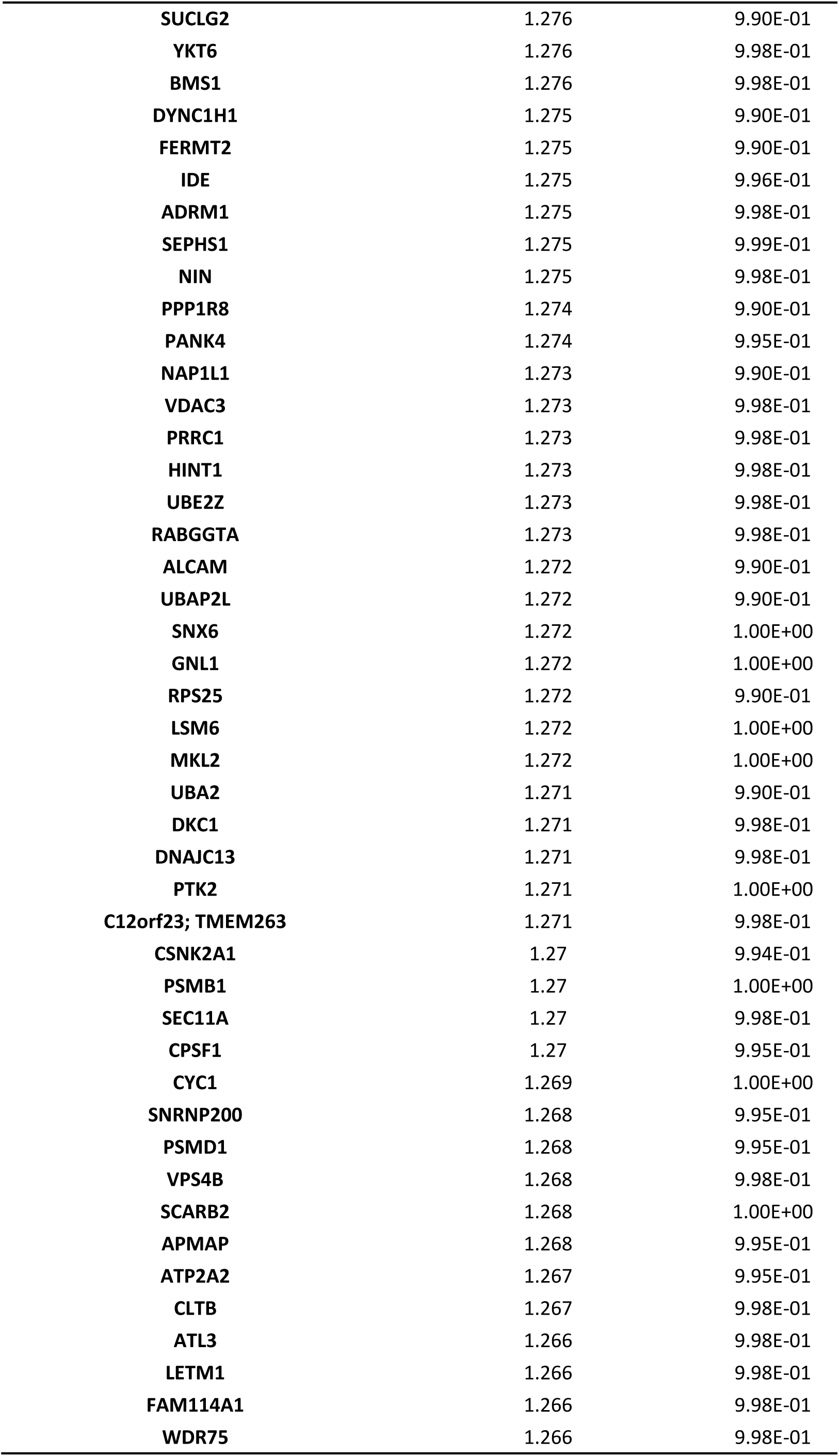
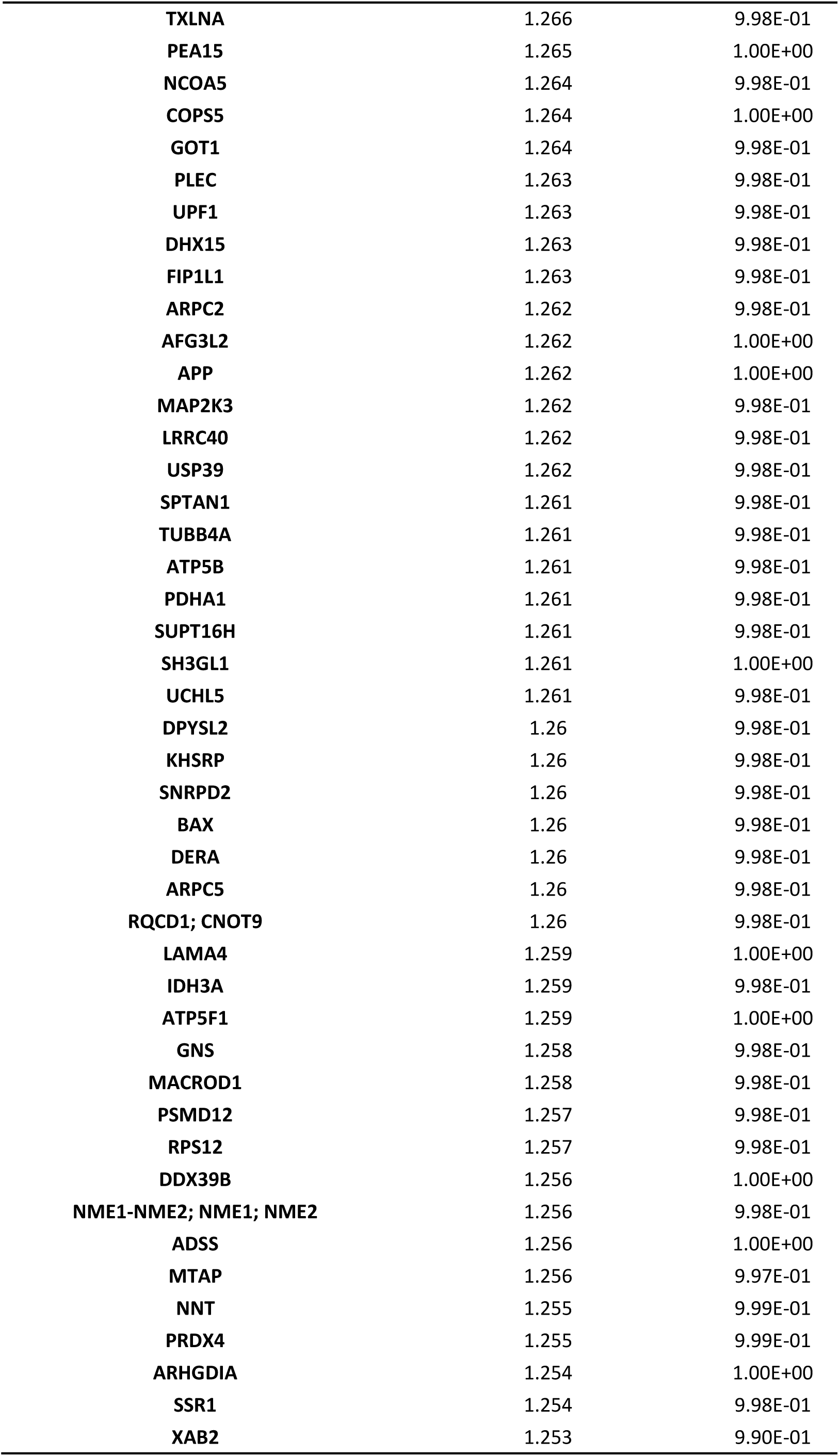
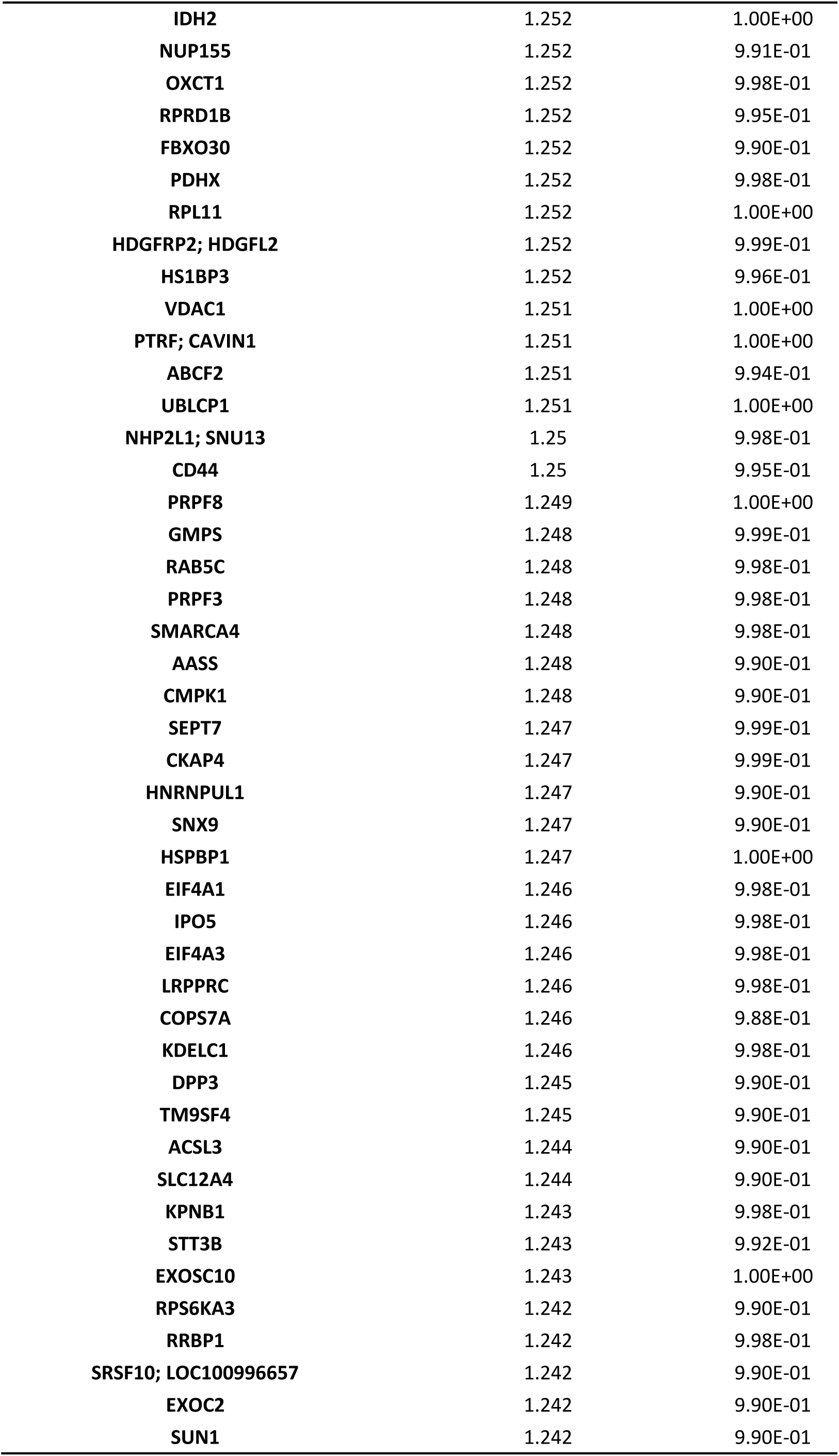
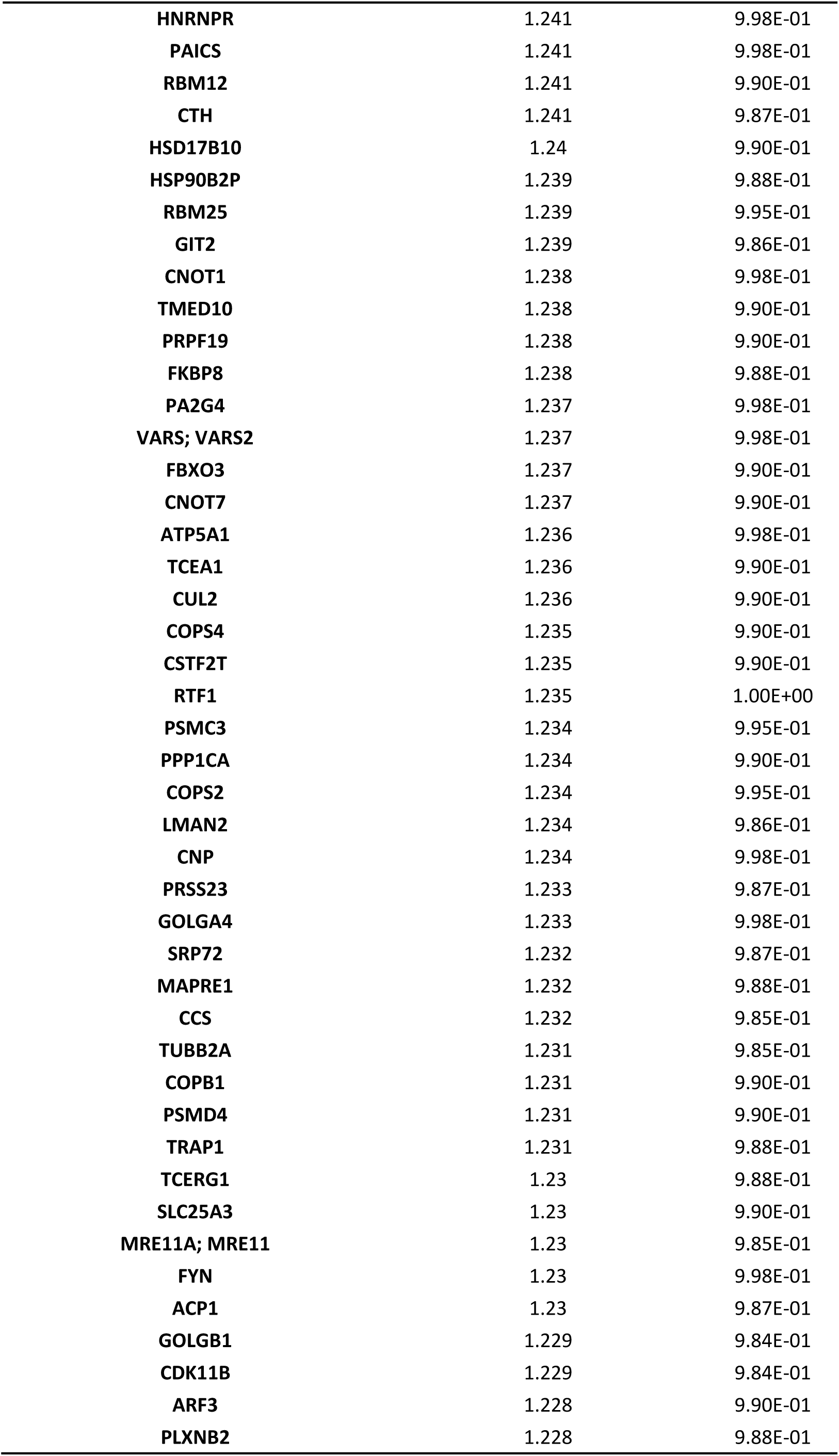
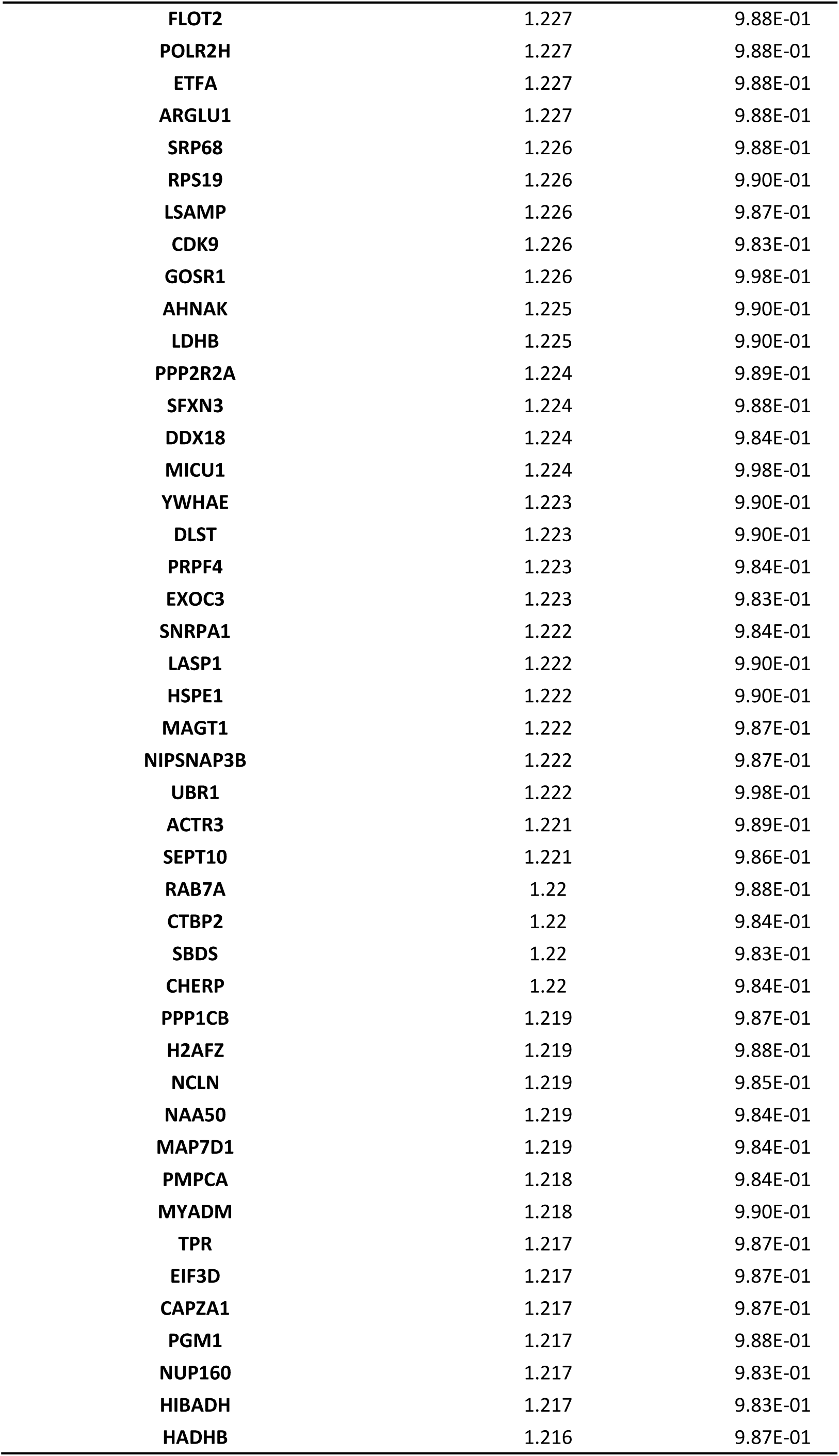
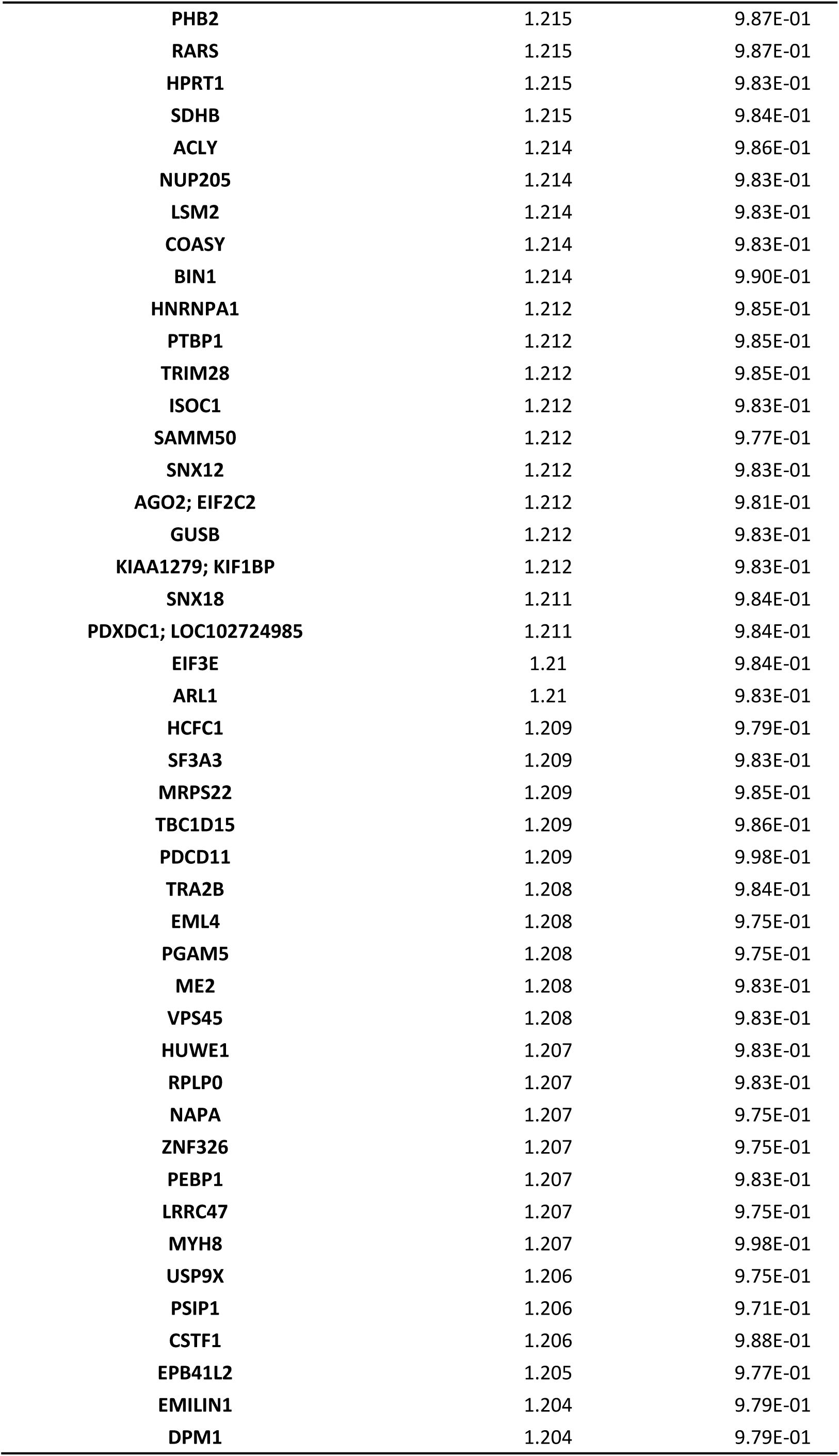
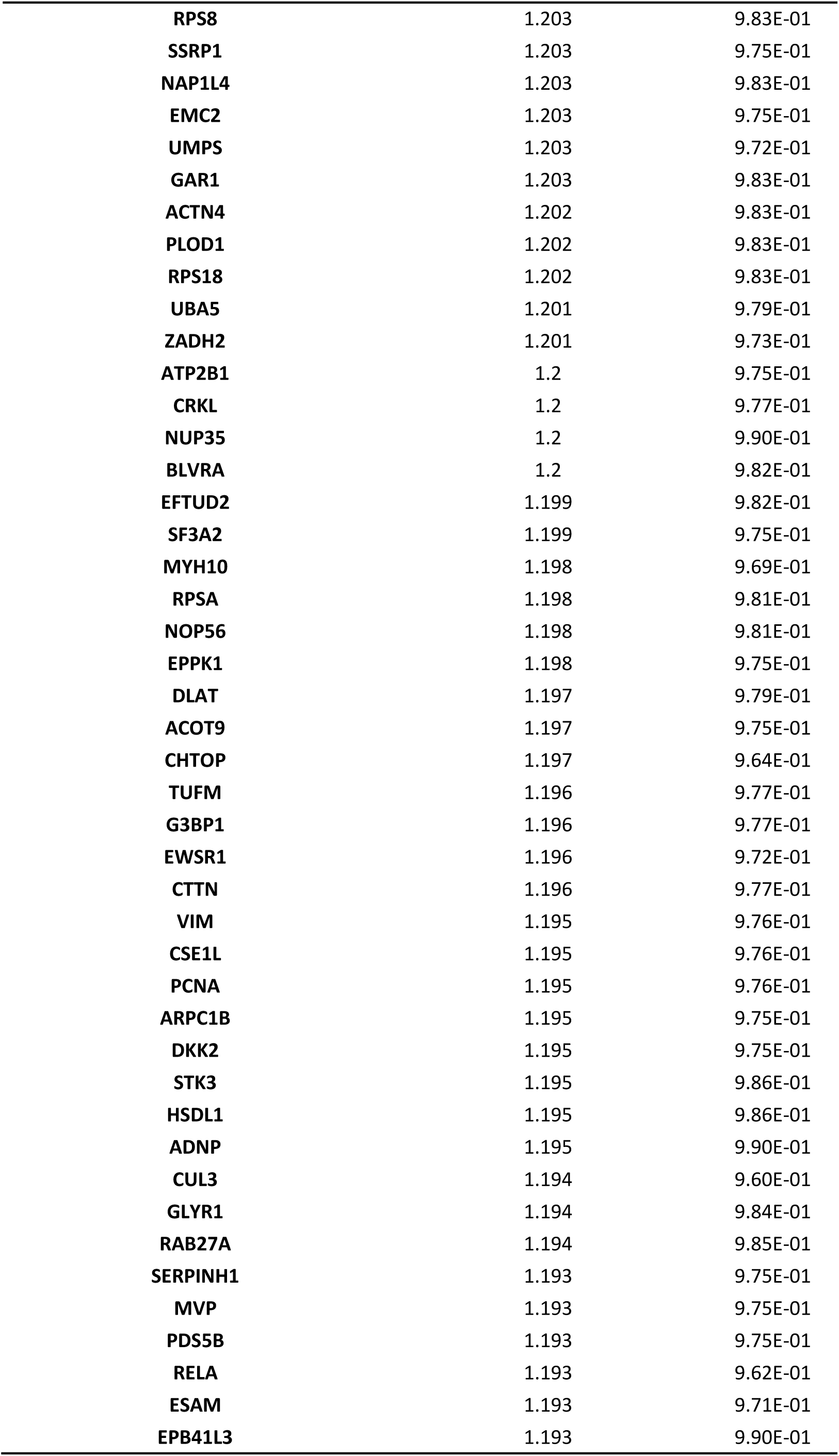
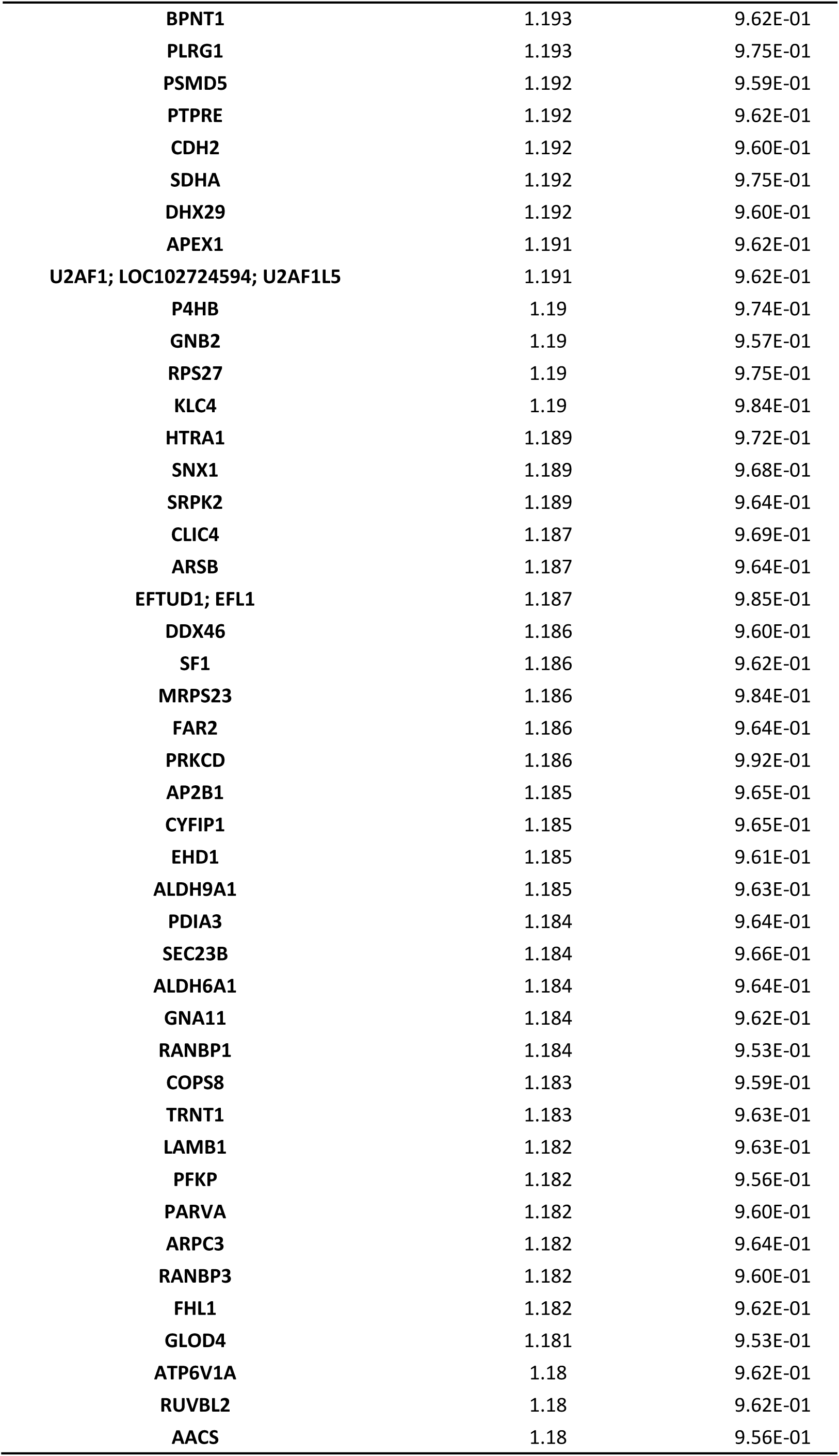
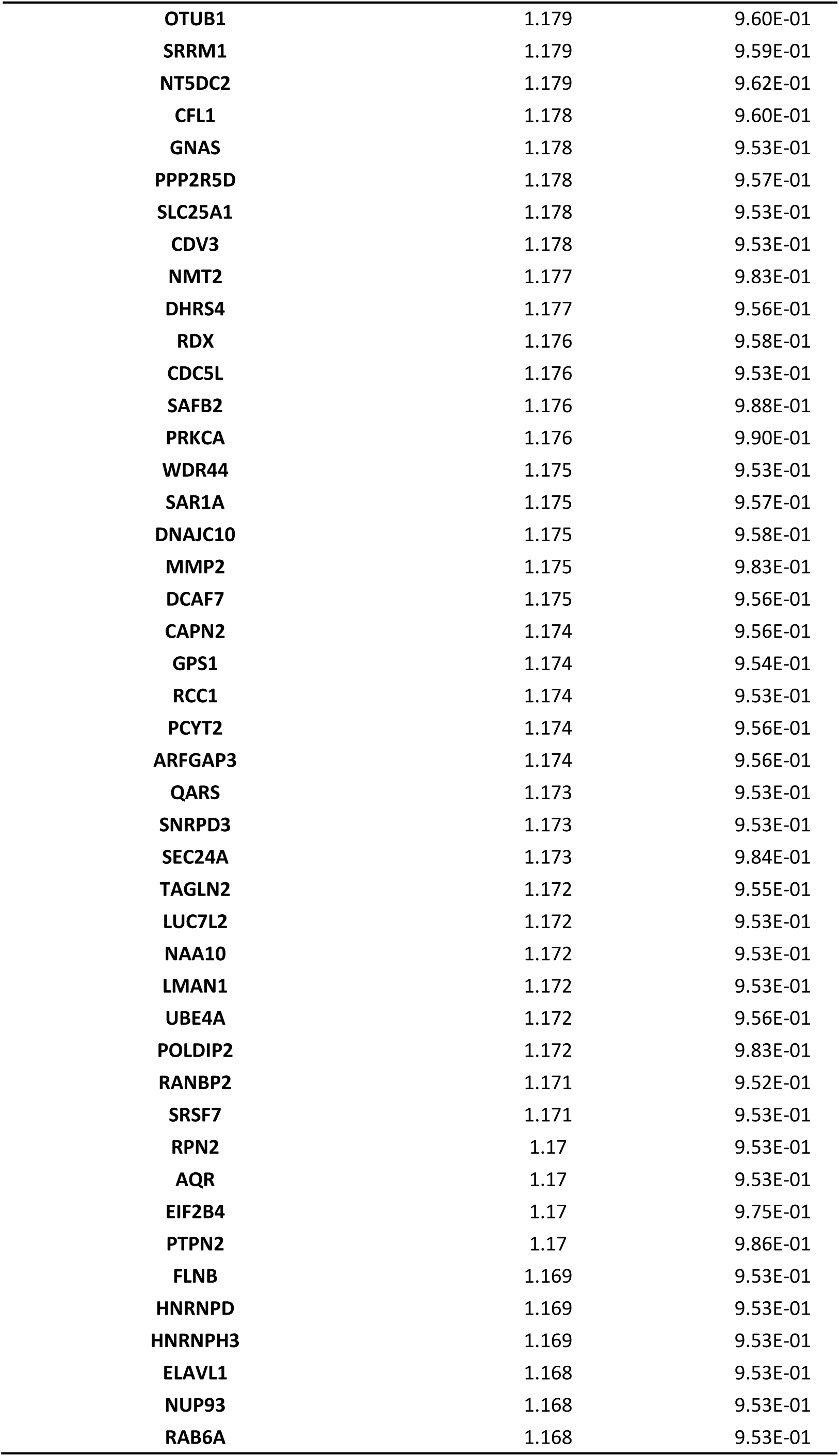
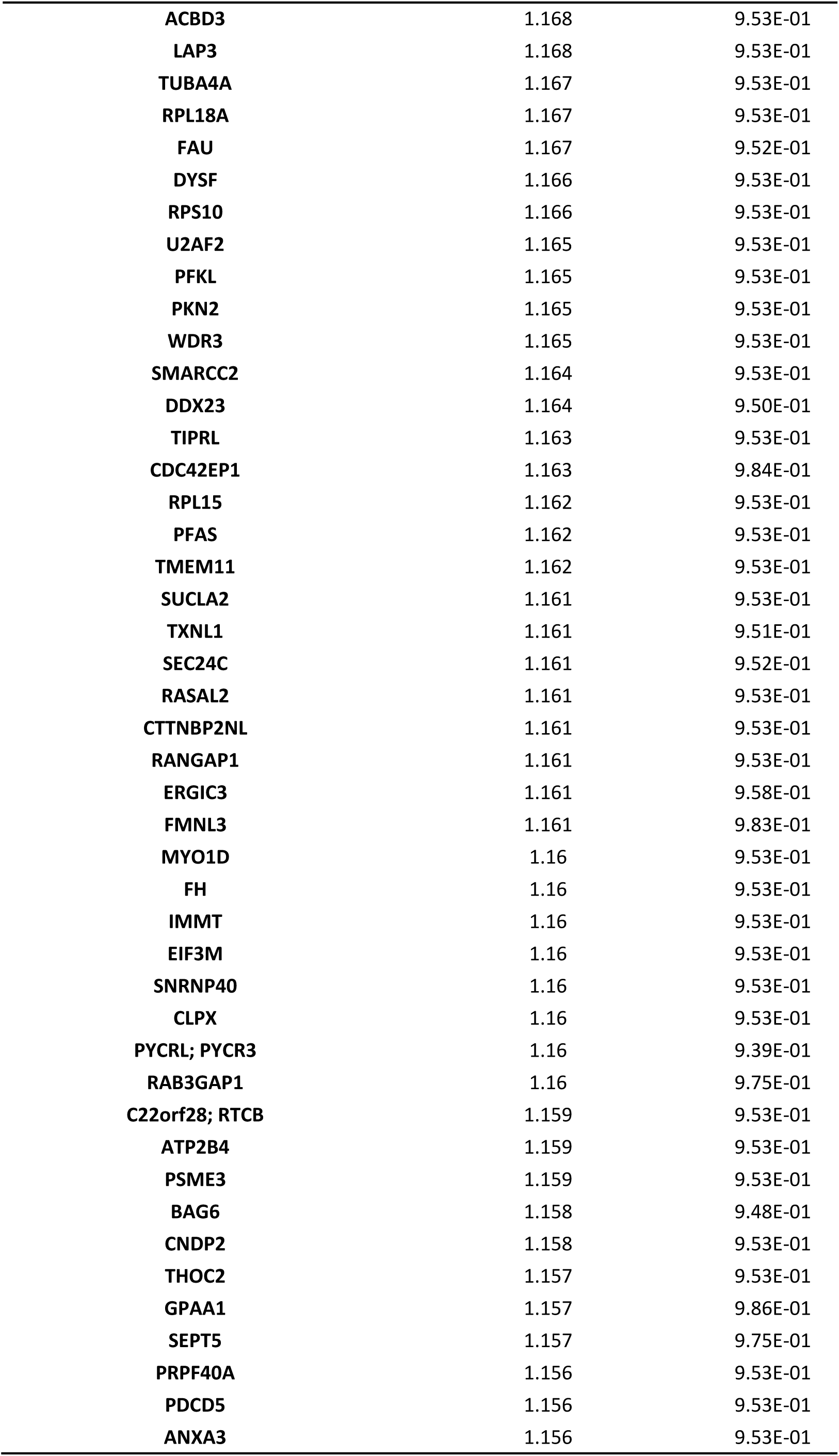
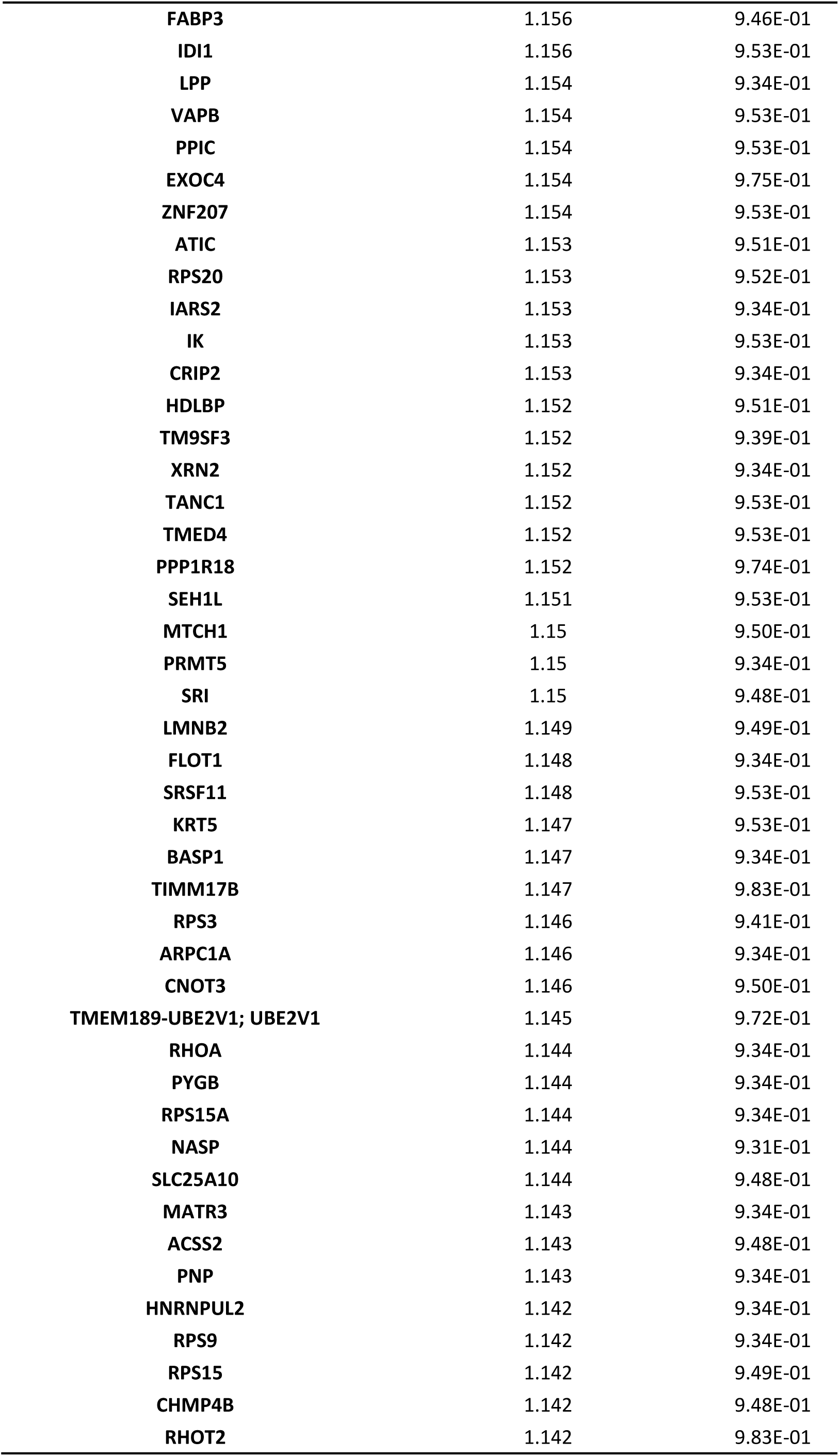
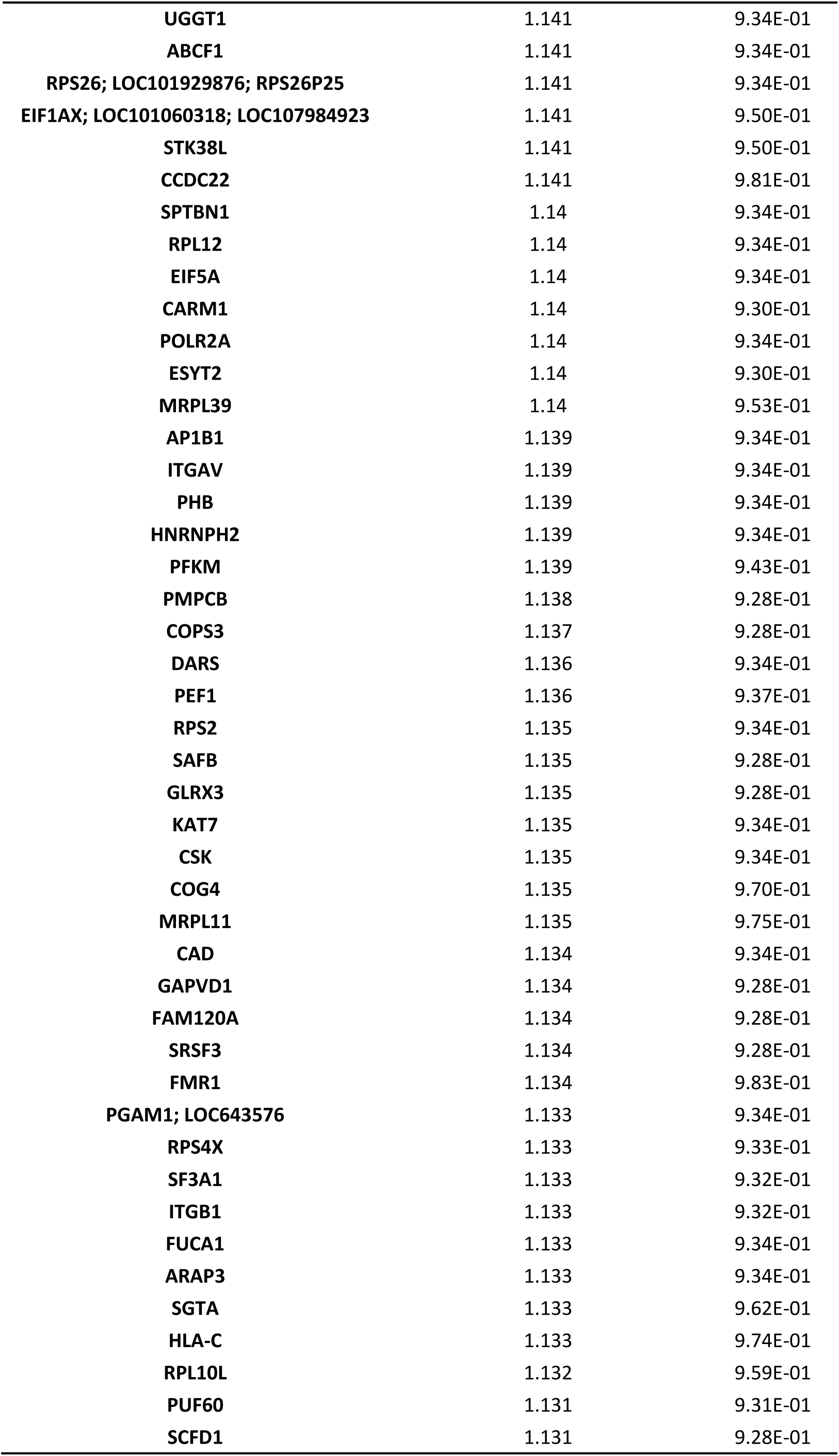
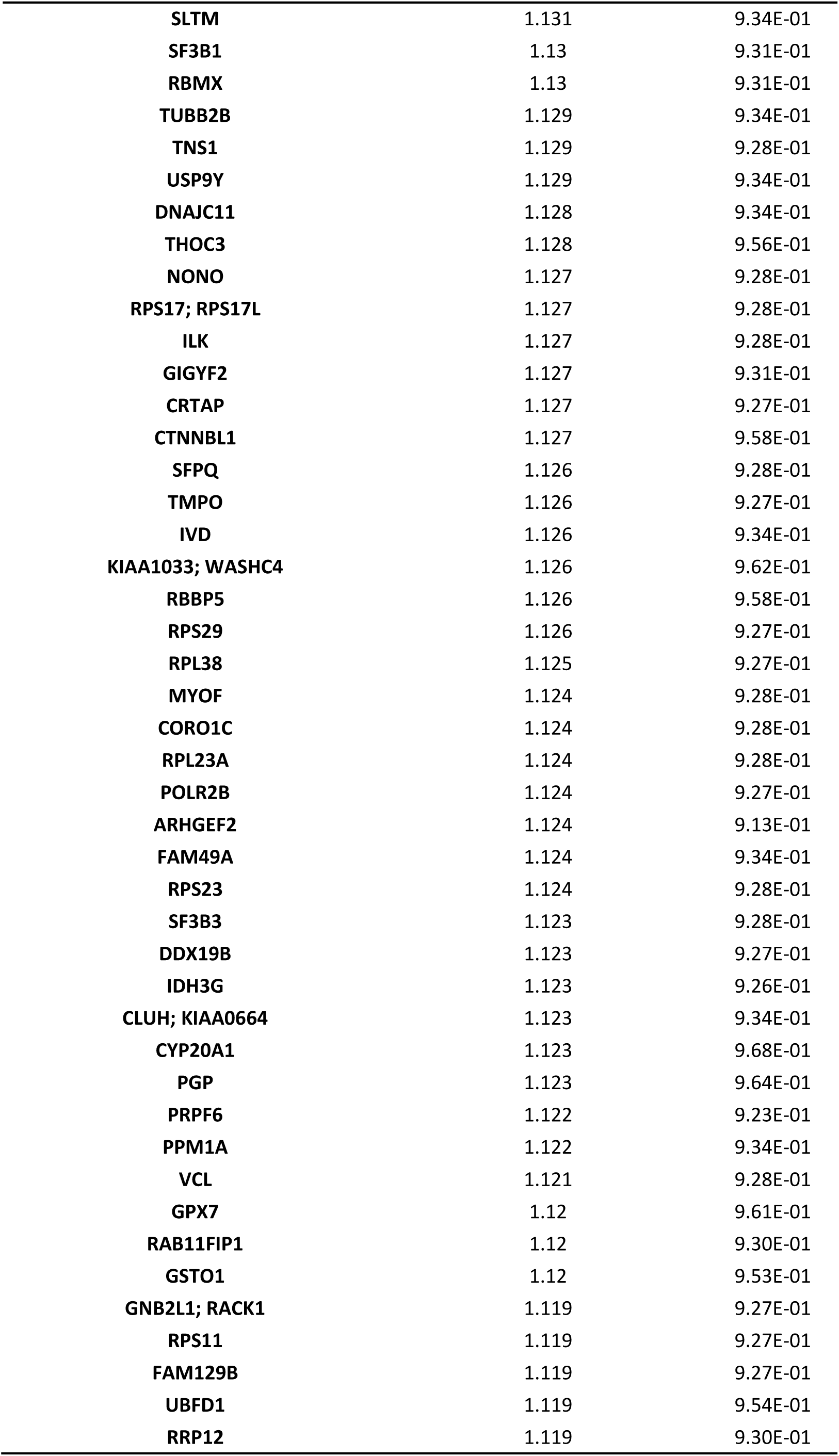
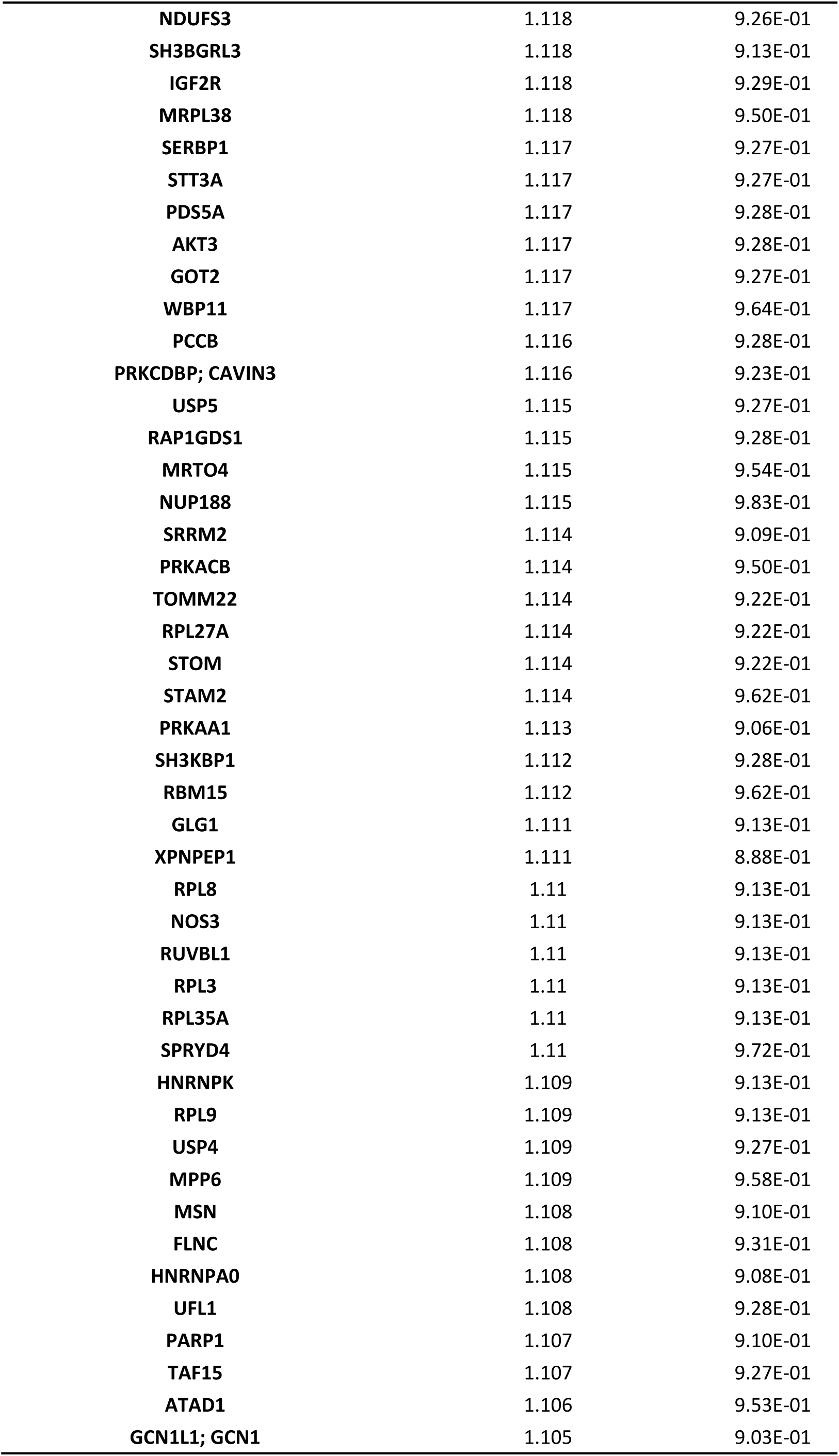
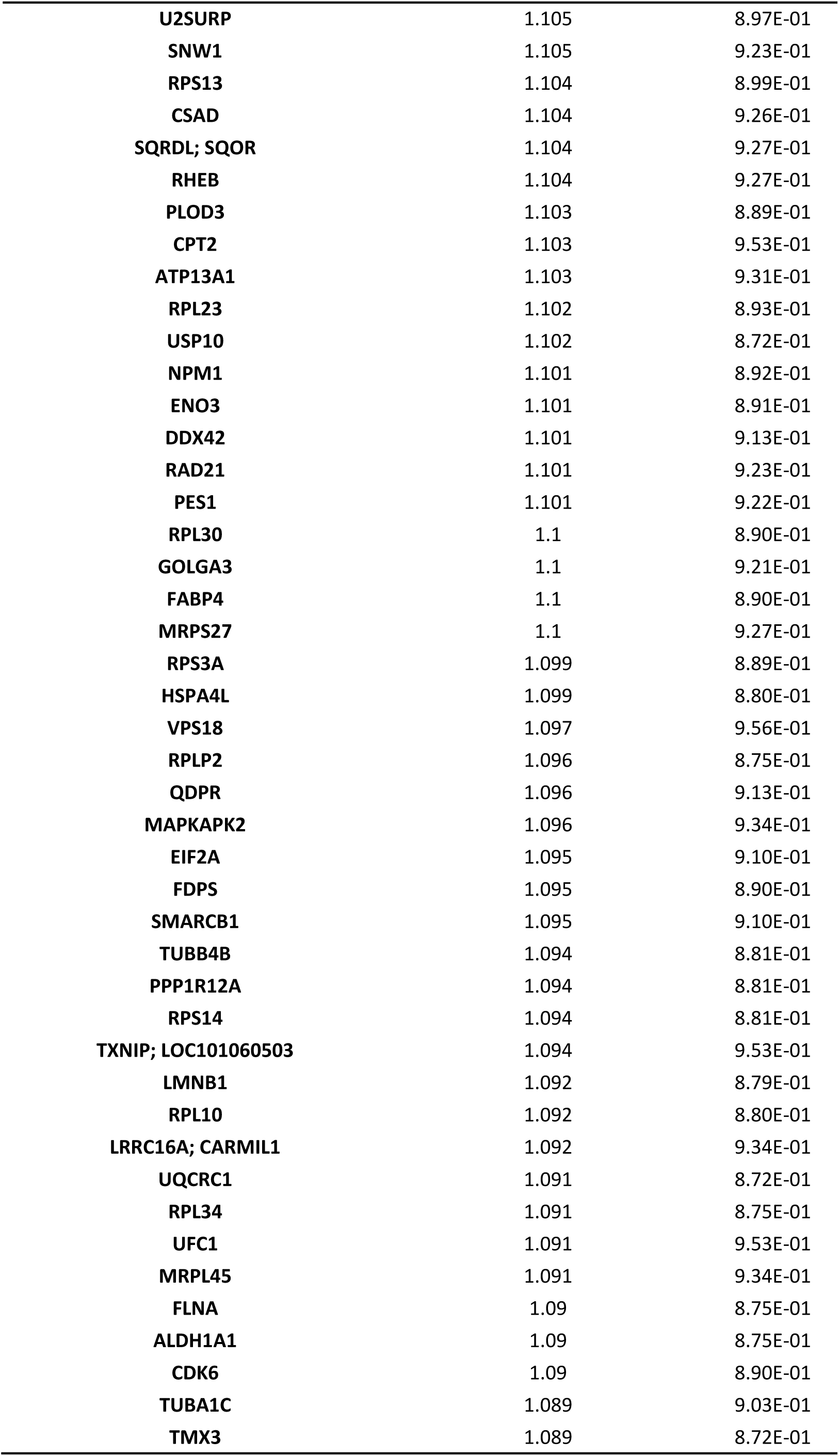
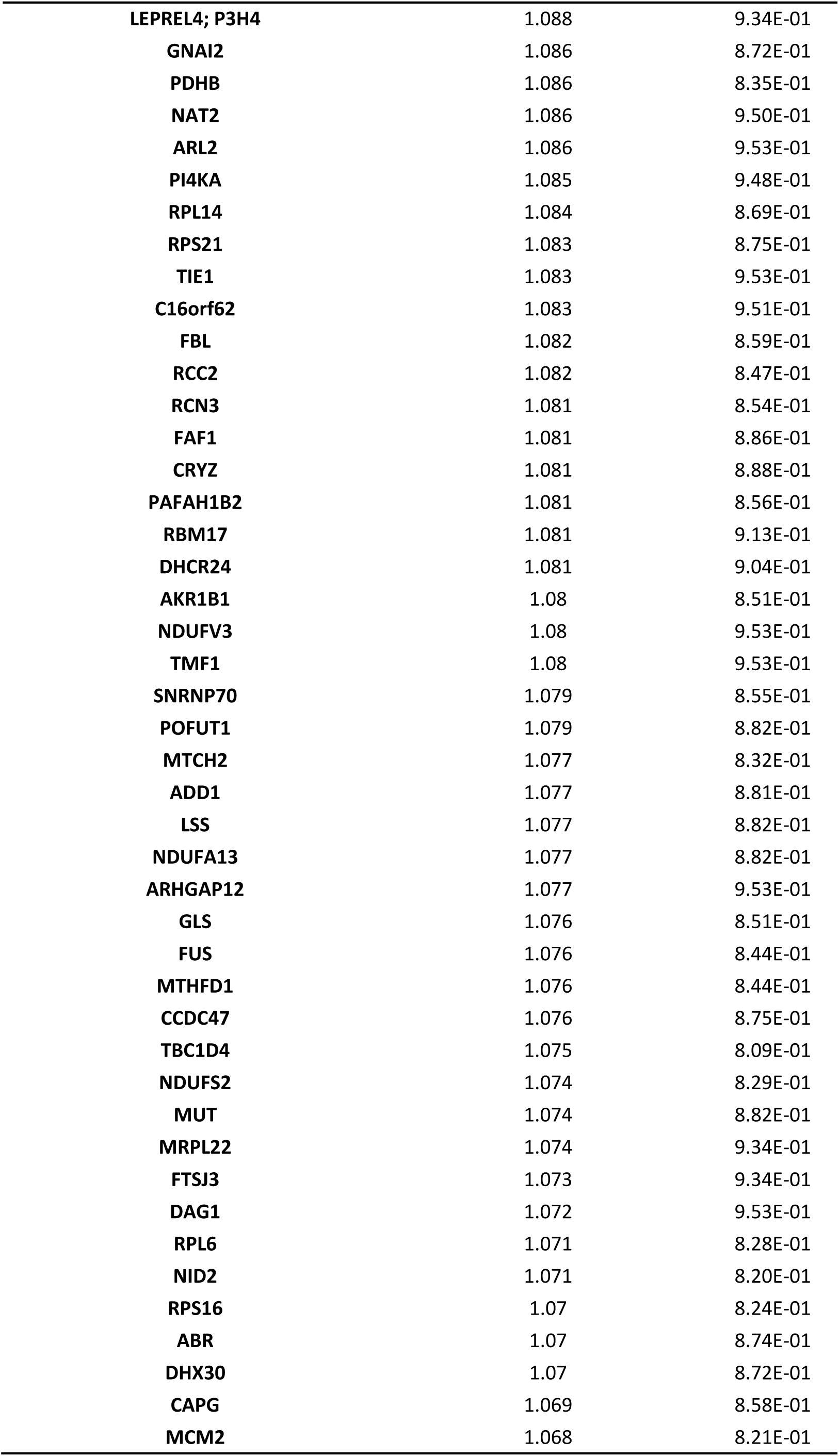
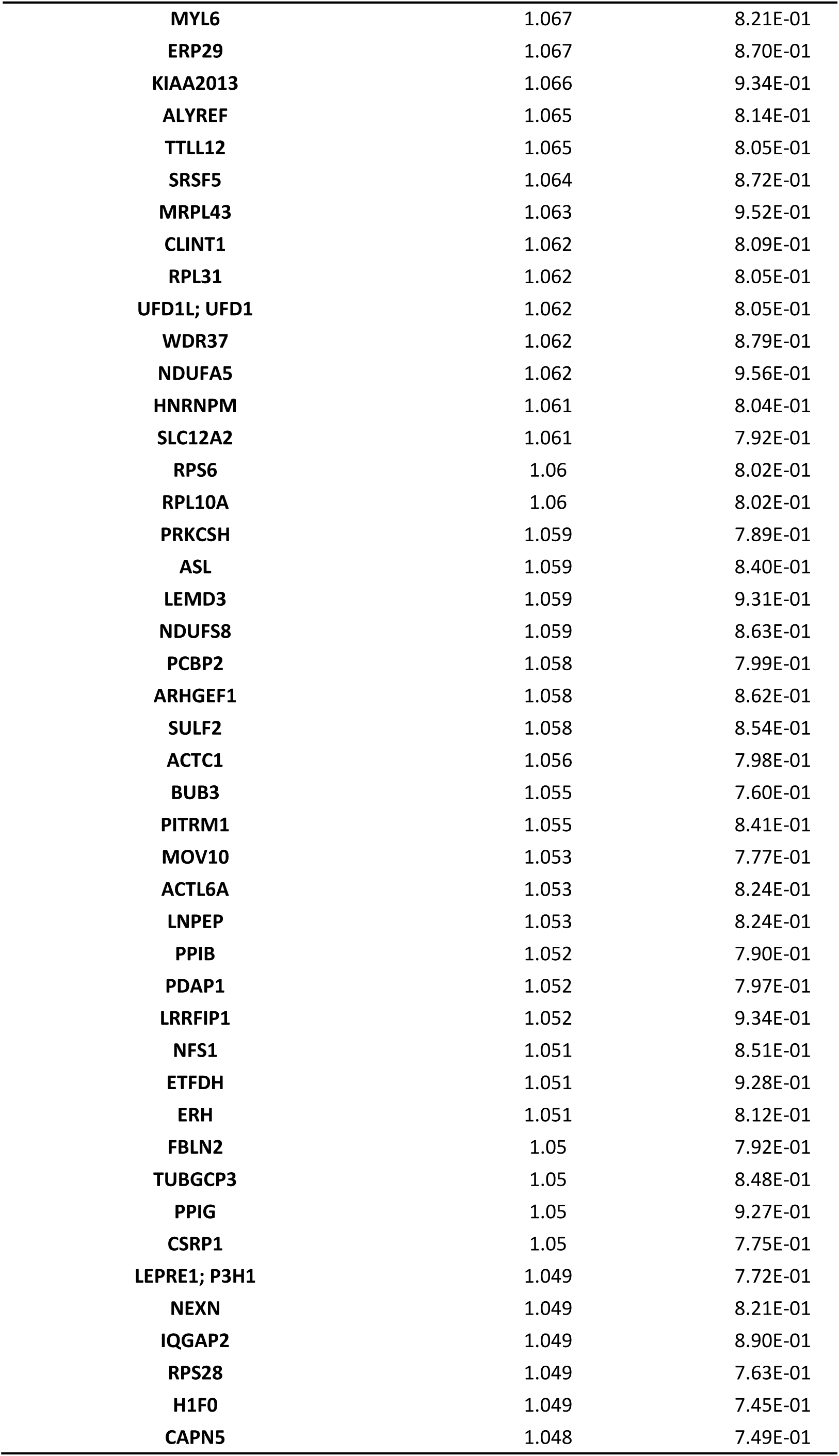
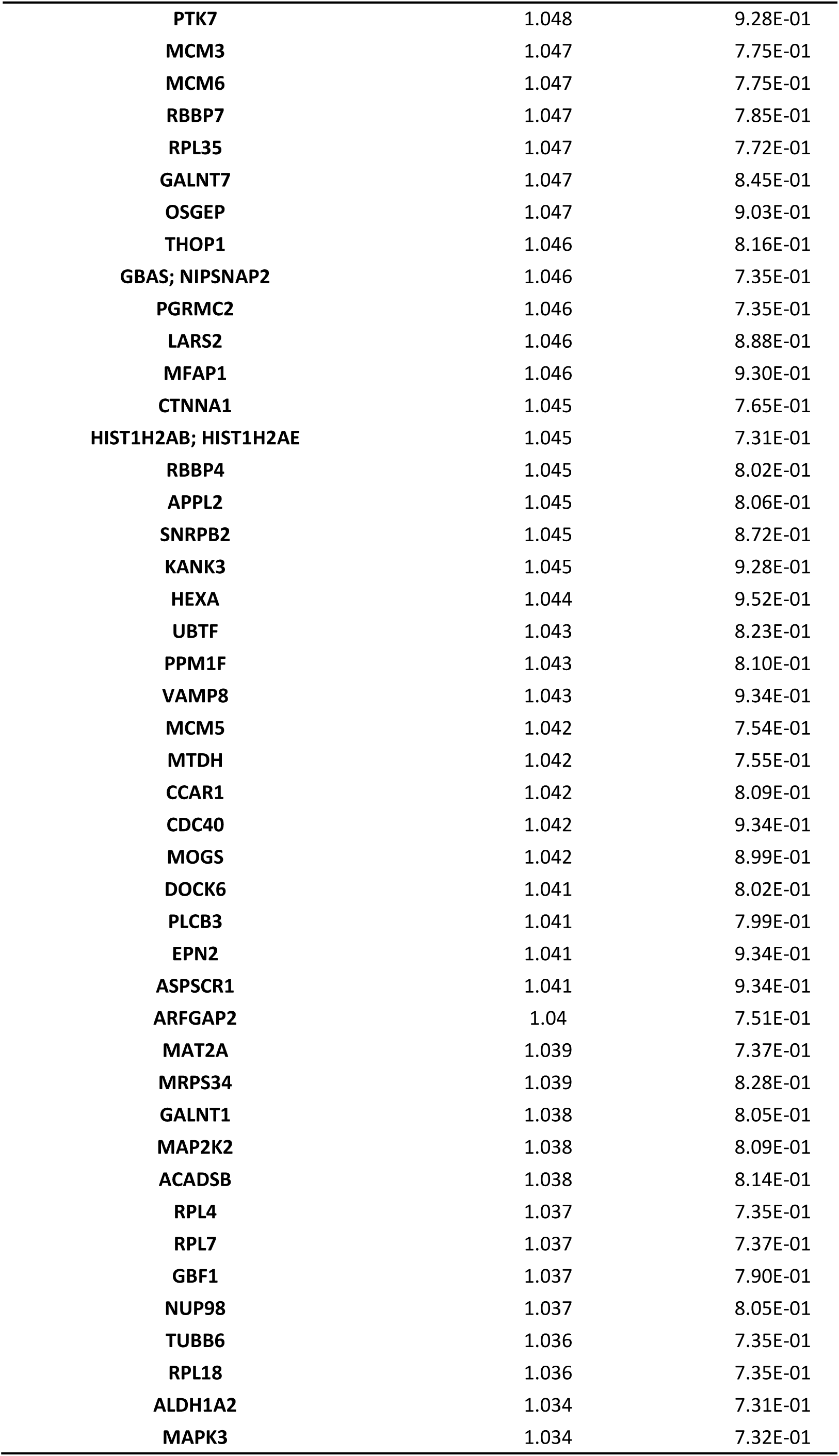
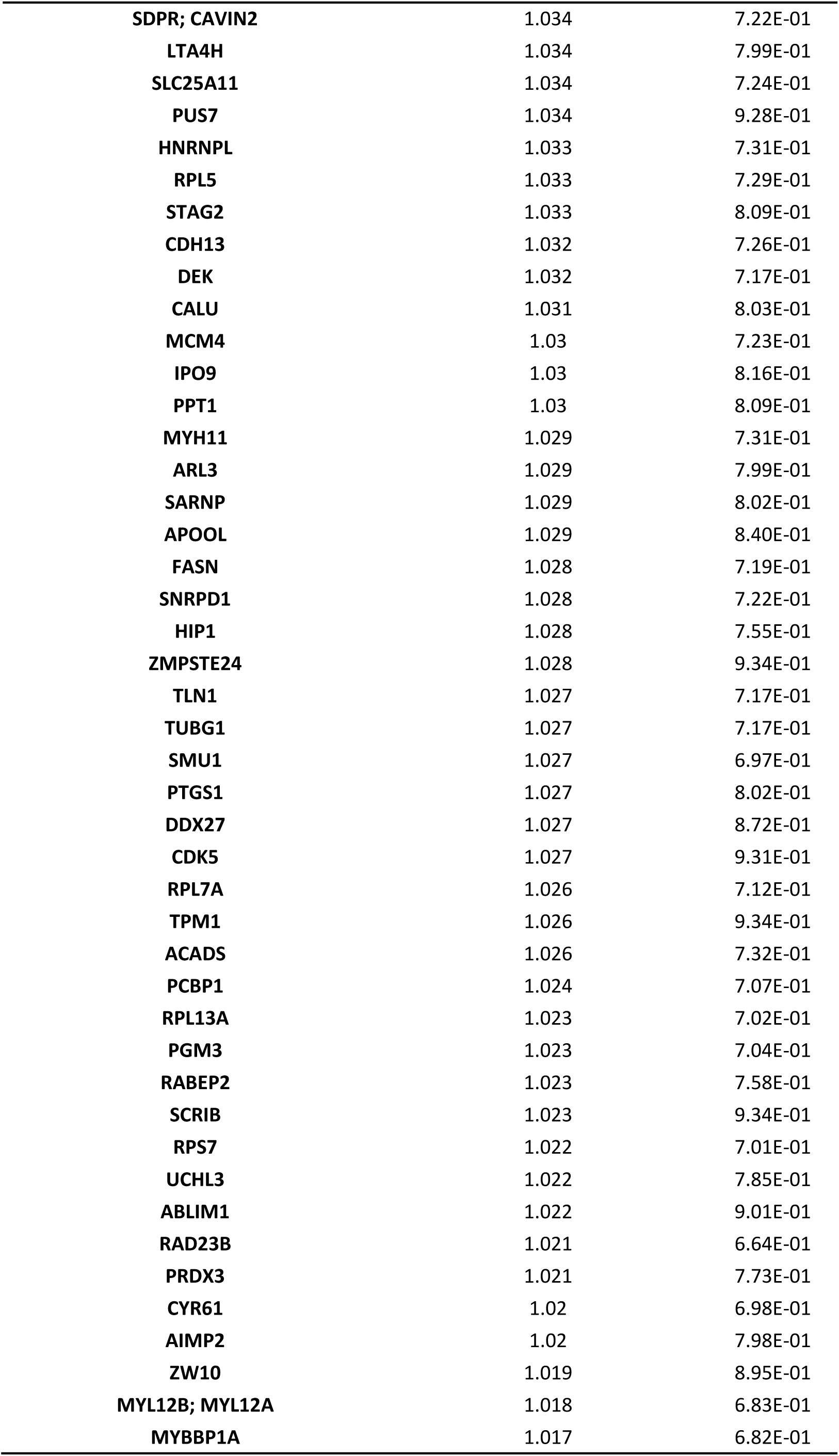
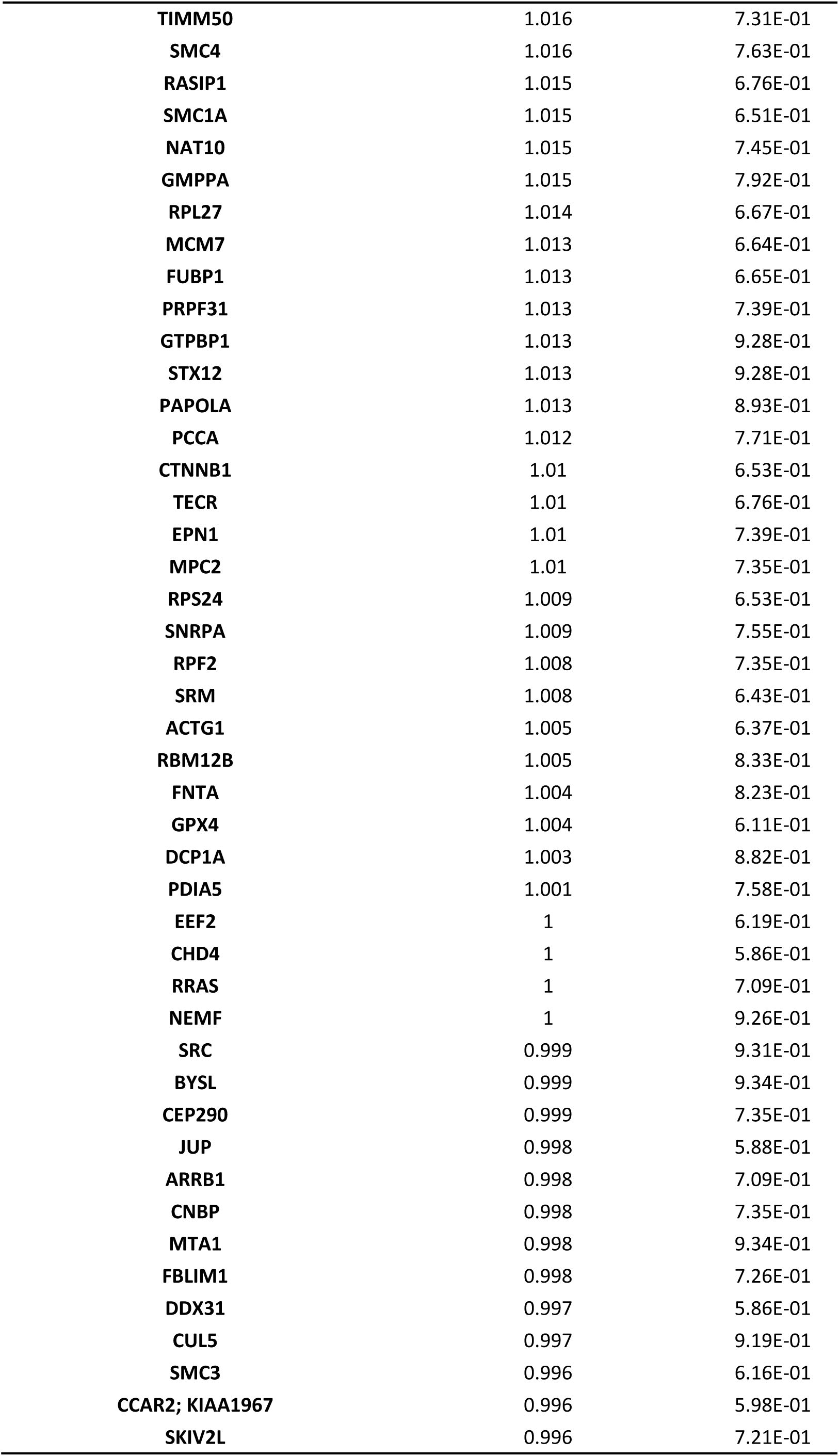
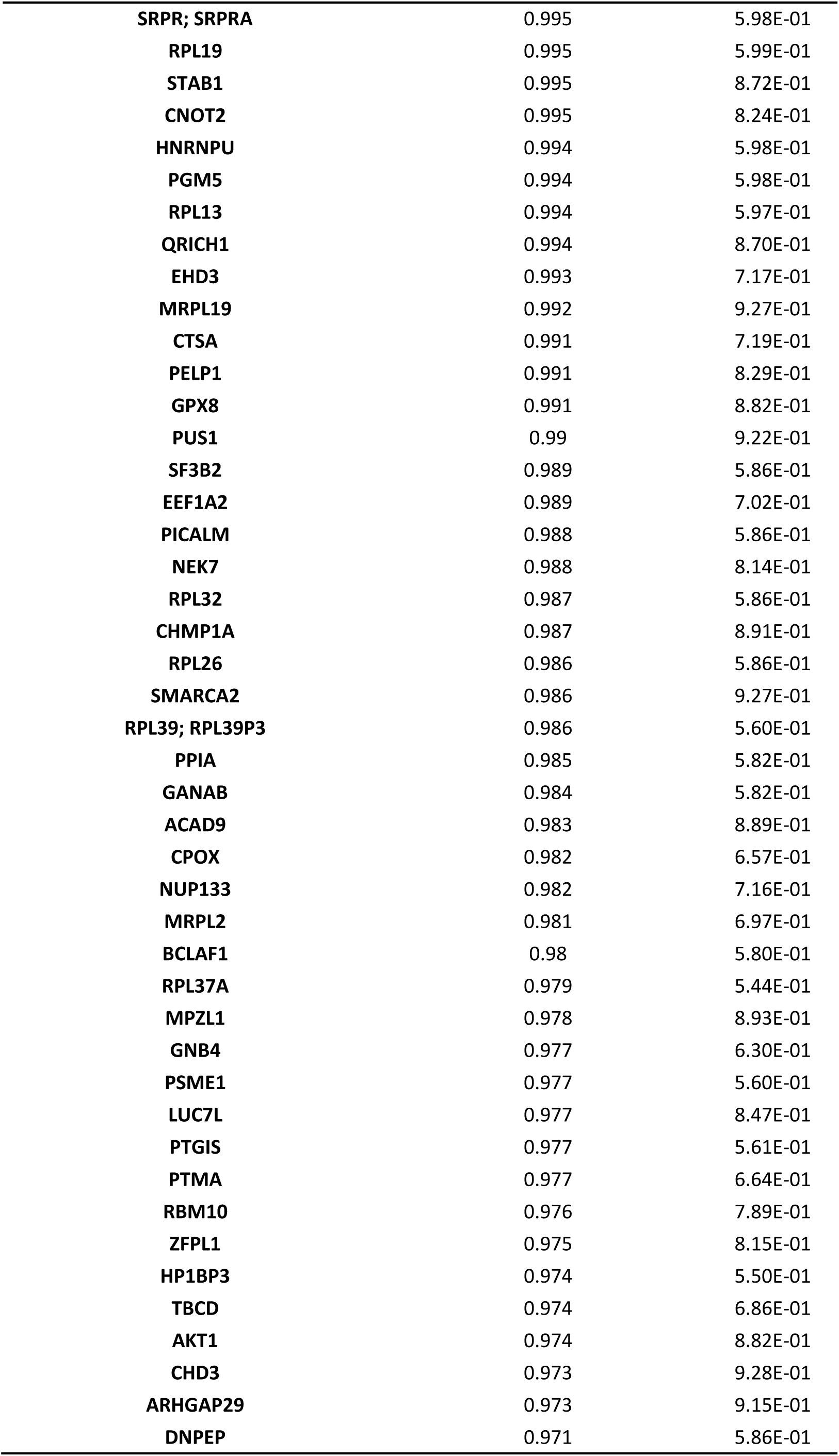
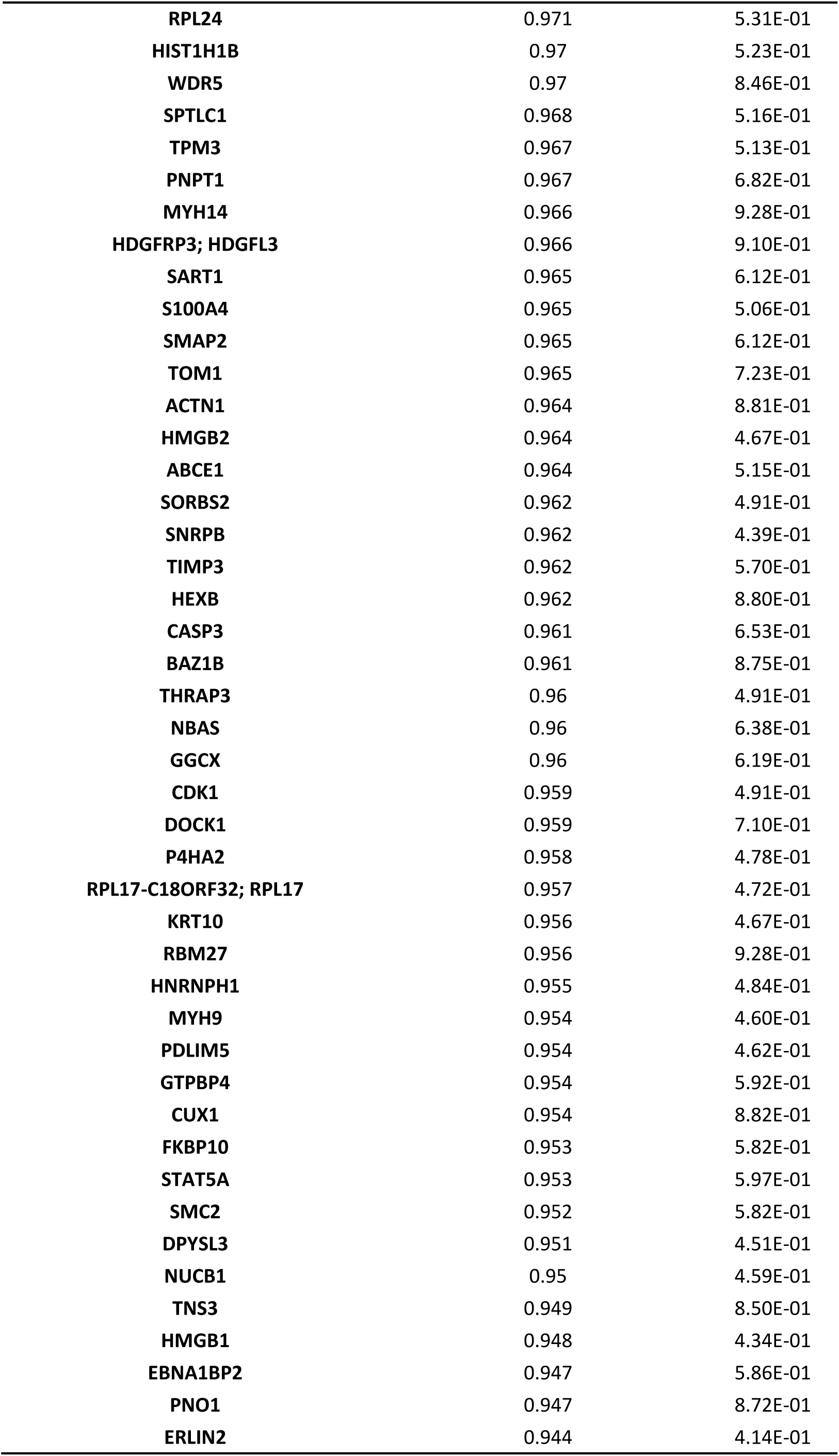
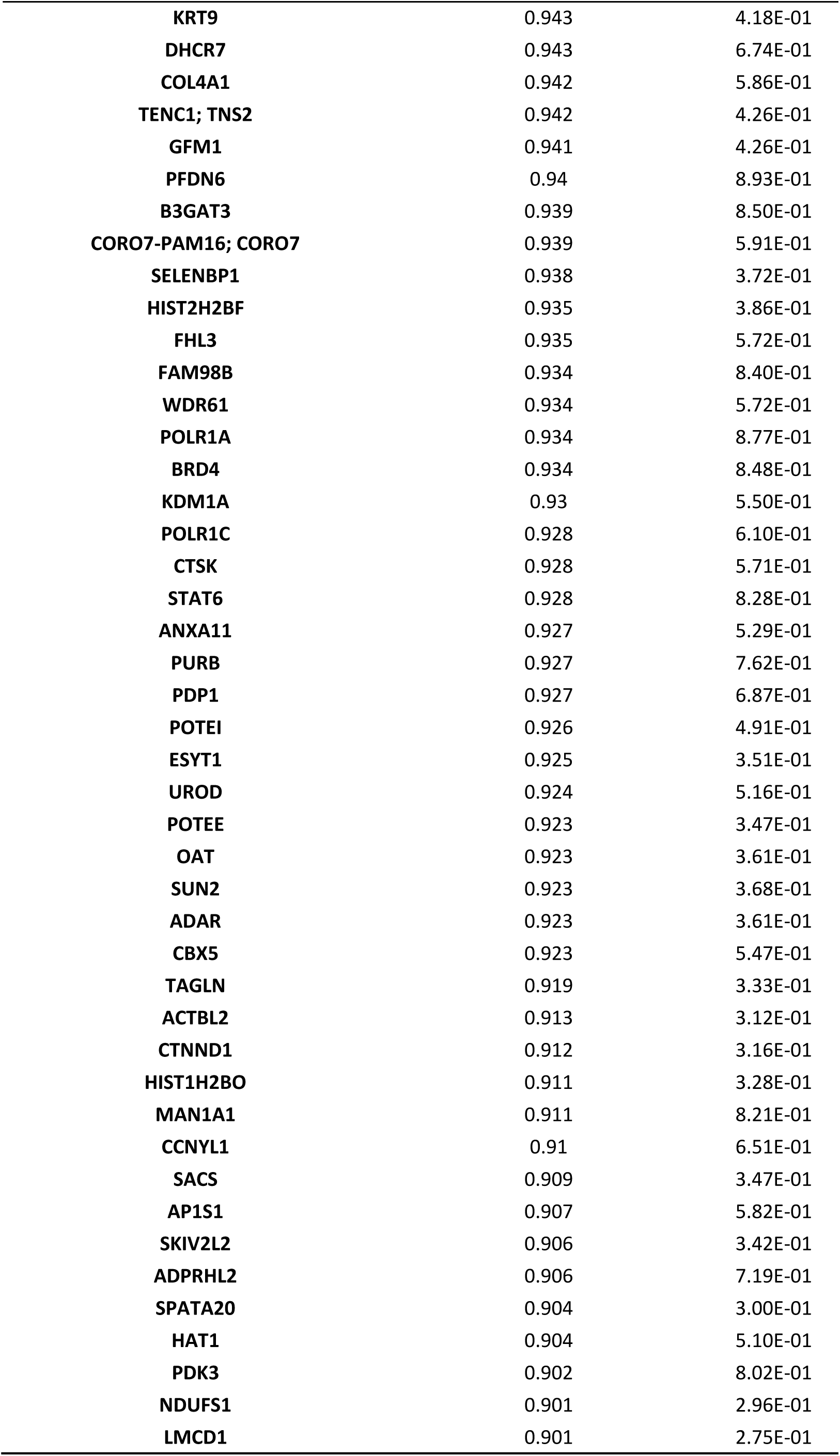
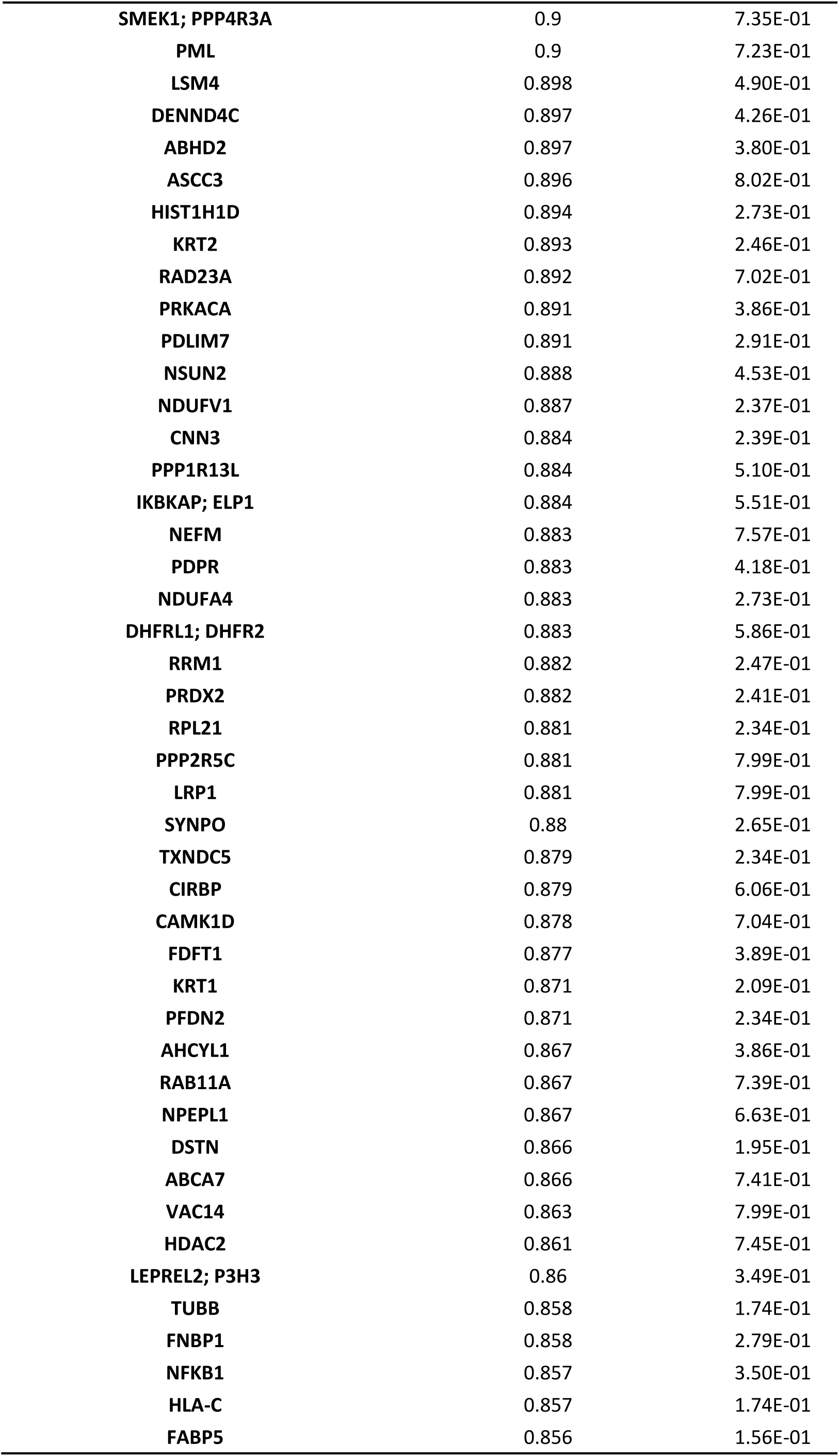
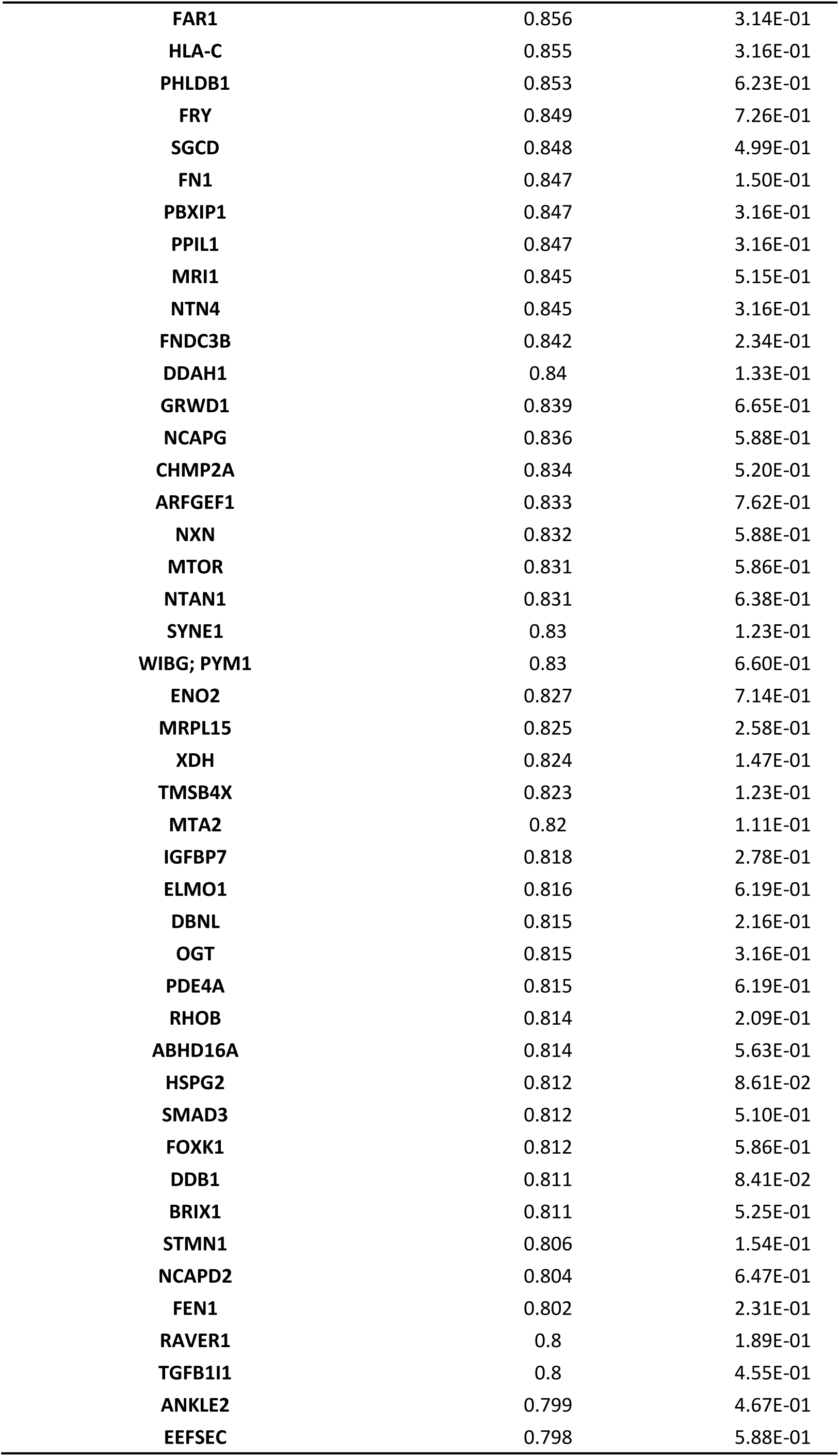
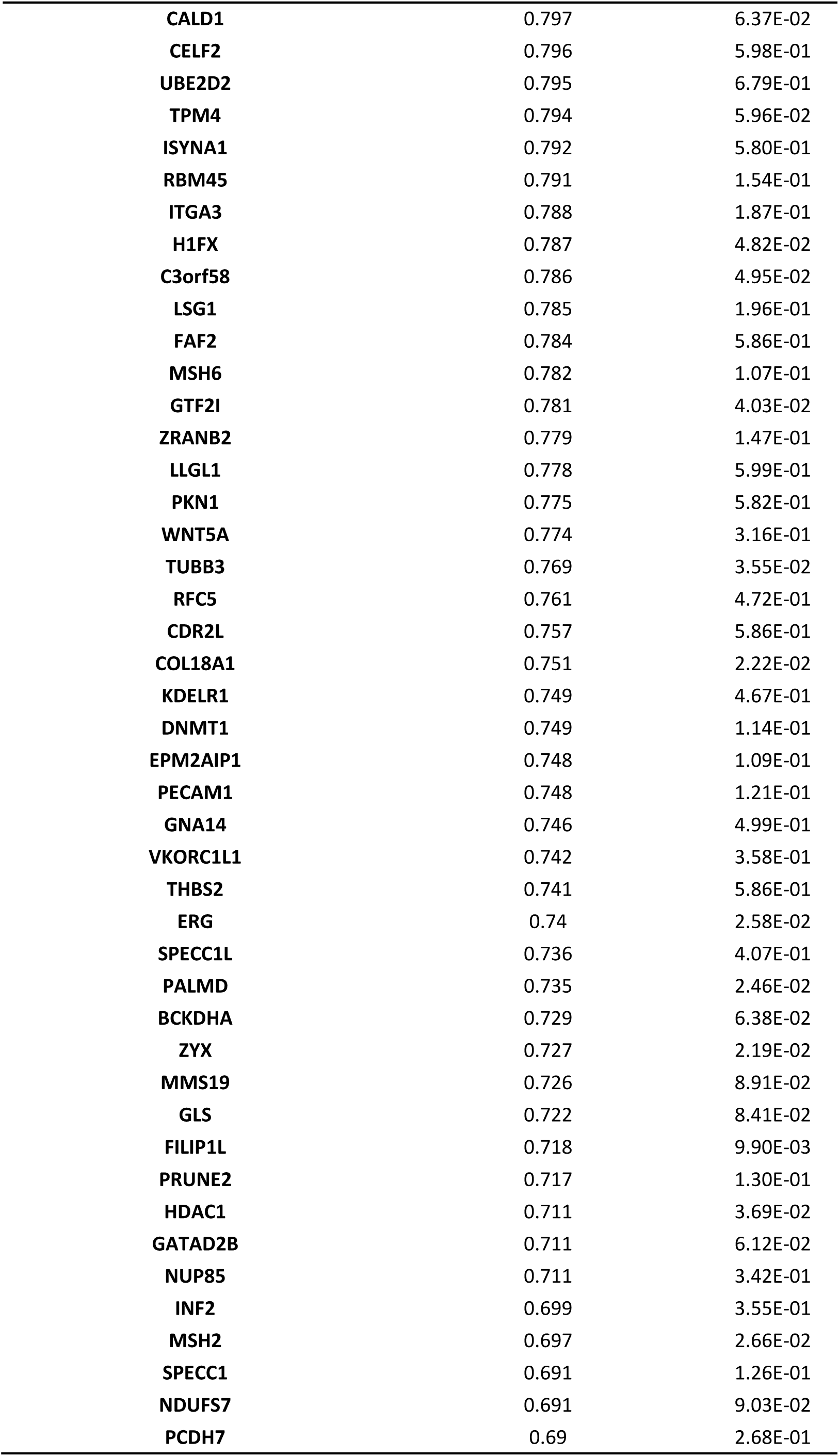
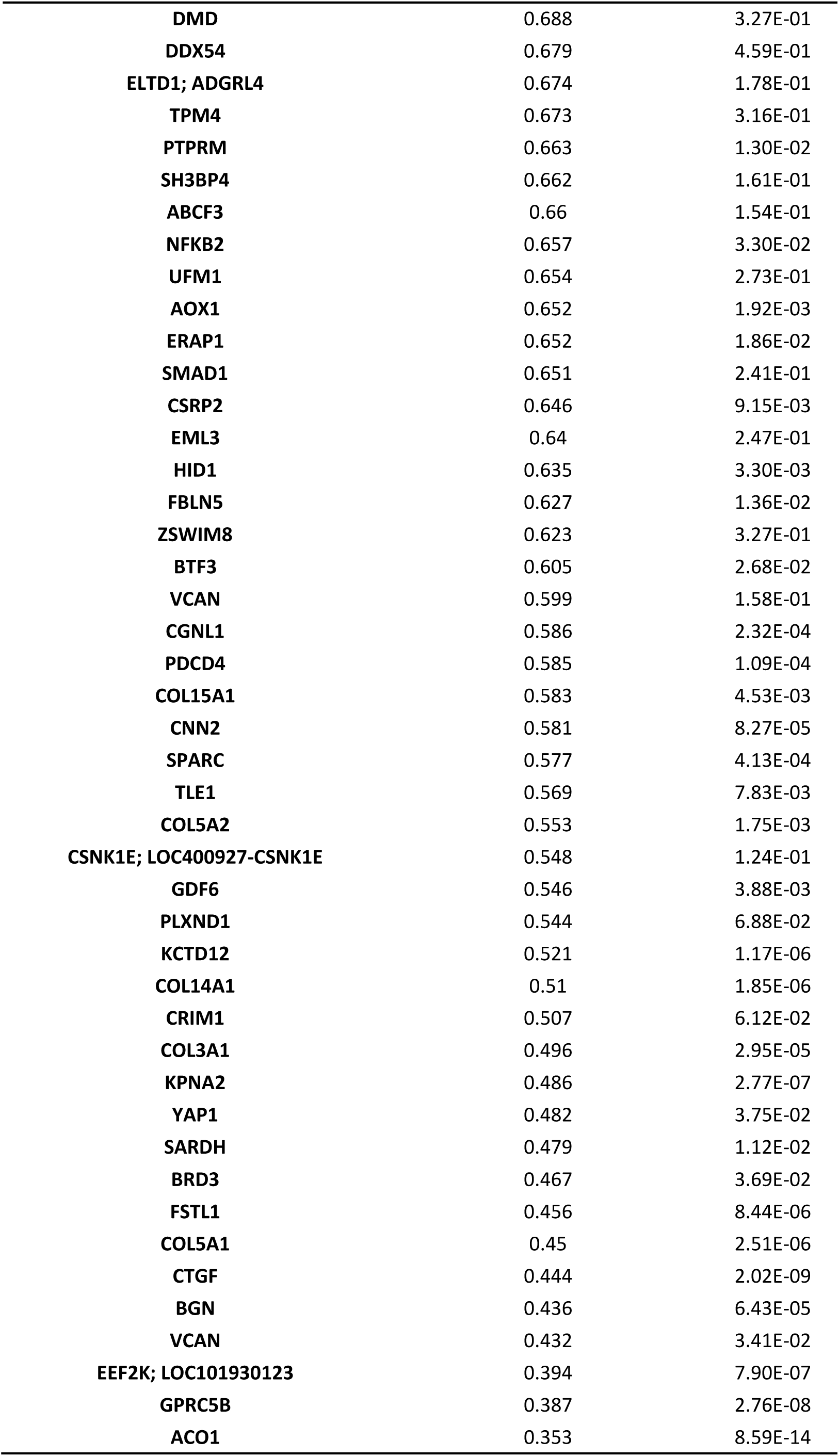
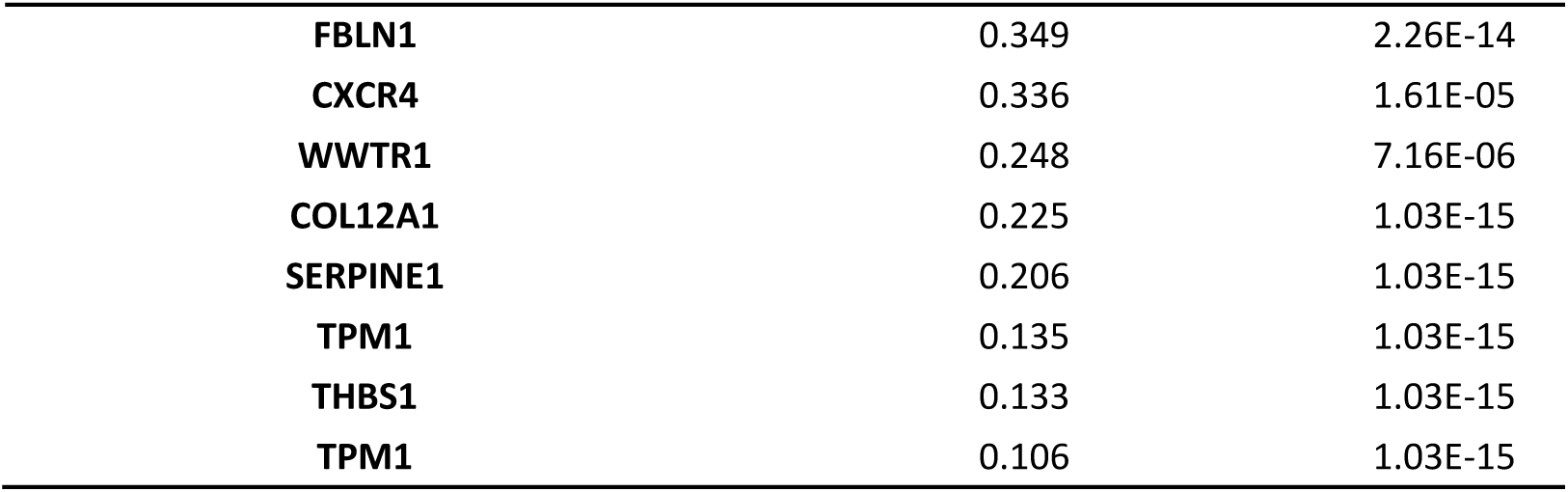
Total of proteins detected with the proteomics analysis in HMECs treated with 100 µM for 24 h.

**Supplementary Table S2.** Proteins detected with the proteomics analysis in HMECs with an abundance ratio (DMF/DMSO) > 1.5 and p-value < 0.01.

| Gene Symbol | Abundance Ratio (DMF/DMSO) | P-value |
| --- | --- | --- |
| HMOX1 | 26.203 | 1.03E-15 |
| GCLC | 7.12 | 1.03E-15 |
| ABHD4 | 7.001 | 1.53E-05 |
| GBE1 | 4.463 | 1.72E-11 |
| PSPH | 4.461 | 3.11E-11 |
| ARHGAP24 | 4.429 | 3.81E-04 |
| SLC2A14 | 4.063 | 7.13E-06 |
| DNM1 | 3.912 | 3.82E-03 |
| TXNRD1 | 3.904 | 2.96E-11 |
| SLC2A1 | 3.72 | 7.95E-03 |
| PCYT1A | 3.526 | 5.65E-05 |
| HSPA1B; HSPA1A | 3.497 | 2.73E-09 |
| DCD | 3.446 | 6.15E-06 |
| VTN | 3.359 | 9.10E-04 |
| VWA5A | 3.236 | 2.94E-04 |
| ASNS | 3.117 | 2.83E-07 |
| RTN4 | 3.107 | 6.39E-06 |
| SQSTM1 | 3.04 | 5.74E-04 |
| ALDH1L2 | 3.012 | 2.51E-05 |
| HSPH1 | 2.991 | 1.27E-06 |
| BAG3 | 2.932 | 4.46E-03 |
| CHORDC1 | 2.765 | 1.16E-03 |
| DNAJA1 | 2.76 | 6.39E-05 |
| SHANK3 | 2.755 | 5.95E-03 |
| ITGA2 | 2.752 | 8.27E-05 |
| SARS | 2.519 | 6.21E-04 |
| CACYBP | 2.516 | 1.62E-03 |
| HSPB1 | 2.51 | 2.97E-04 |
| CLIP1 | 2.496 | 8.65E-03 |
| PTGES3 | 2.445 | 1.62E-03 |
| TXN | 2.34 | 1.87E-03 |
| COTL1 | 2.332 | 8.80E-03 |
| HBD | 2.324 | 4.52E-03 |
| FKBP5 | 2.288 | 6.38E-03 |
| SNTB2 | 2.282 | 7.00E-03 |
| SNX3 | 2.189 | 9.15E-03 |
| HECTD1 | 2.183 | 8.80E-03 |

**Supplementary Table S3.** Proteins detected with the proteomics analysis in HMECs with an abundance ratio (DMF/DMSO) < 0.75 and p-value < 0.01.

| Gene Symbol | Abundance Ratio (DMF/DMSO) | P-value |
| --- | --- | --- |
| TPM1 | 0.106 | 1.03E-15 |
| THBS1 | 0.133 | 1.03E-15 |
| TPM1 | 0.135 | 1.03E-15 |
| SERPINE1 | 0.206 | 1.03E-15 |
| COL12A1 | 0.225 | 1.03E-15 |
| WWTR1 | 0.248 | 7.16E-06 |
| CXCR4 | 0.336 | 1.61E-05 |
| FBLN1 | 0.349 | 2.26E-14 |
| ACO1 | 0.353 | 8.59E-14 |
| GPRC5B | 0.387 | 2.76E-08 |
| EEF2K; LOC101930123 | 0.394 | 7.90E-07 |
| BGN | 0.436 | 6.43E-05 |
| CTGF | 0.444 | 2.02E-09 |
| COL5A1 | 0.45 | 2.51E-06 |
| FSTL1 | 0.456 | 8.44E-06 |
| KPNA2 | 0.486 | 2.77E-07 |
| COL3A1 | 0.496 | 2.95E-05 |
| COL14A1 | 0.51 | 1.85E-06 |
| KCTD12 | 0.521 | 1.17E-06 |
| GDF6 | 0.546 | 3.88E-03 |
| COL5A2 | 0.553 | 1.75E-03 |
| TLE1 | 0.569 | 7.83E-03 |
| SPARC | 0.577 | 4.13E-04 |
| CNN2 | 0.581 | 8.27E-05 |
| COL15A1 | 0.583 | 4.53E-03 |
| PDCD4 | 0.585 | 1.09E-04 |
| CGNL1 | 0.586 | 2.32E-04 |
| HID1 | 0.635 | 3.30E-03 |
| CSRP2 | 0.646 | 9.15E-03 |
| AOX1 | 0.652 | 1.92E-03 |
| FILIP1L | 0.718 | 9.90E-03 |

## Notes

### Competing Interest Statement

The authors have declared no competing interest.

